# Dual Adhesive Unipolar Polysaccharides Synthesized by Overlapping Biosynthetic Pathways in *Agrobacterium tumefaciens*

**DOI:** 10.1101/2021.04.22.440995

**Authors:** Maureen C. Onyeziri, Ramya Natarajan, Gail G. Hardy, Jing Xu, Ian P. Reynolds, Jinwoo Kim, Peter M. Merritt, Thomas Danhorn, Michael E. Hibbing, Alexandra J. Weisberg, Jeff H. Chang, Clay Fuqua

## Abstract

*Agrobacterium tumefaciens*, is a member of the Alphaproteobacteria that pathogenizes plants, and associates with biotic and abiotic surfaces via a single cellular pole. *A. tumefaciens* produces the unipolar polysaccharide (UPP) at the site of surface contact. UPP production is normally surface-contact inducible, but elevated levels of the second messenger cyclic diguanylate monophosphate (cdGMP) bypass this requirement. Multiple lines of evidence suggest that the UPP has a central polysaccharide component. Using an *A. tumefaciens* derivative with elevated cdGMP and mutationally disabled for other dispensable polysaccharides, a series of related genetic screens have identified a large number of genes involved in UPP biosynthesis, most of which are Wzx-Wzy-type polysaccharide biosynthetic components. Extensive analyses of UPP production in these mutants have revealed that the UPP is comprised of two genetically, chemically and spatially discrete forms of polysaccharide, and that each requires a specific Wzy-type polymerase. Other important biosynthetic, processing and regulatory functions for UPP production are also revealed, some of which are common to both polysaccharides, and a subset of which are specific to each species. Many of the UPP genes identified are conserved among diverse rhizobia, whereas others are more lineage specific.

**Plain language summary:** Bacteria attach to a wide variety of surfaces including host tissues, via externalized structures described as adhesins. We define a large set of genes involved in synthesis of a complex unipolar adhesin comprised of two distinct polysaccharides, that is required for surface attachment by the plant-associated pathogen *Agrobacterium tumefaciens*.

## Introduction

Many pathogenic bacteria stably associate with the tissues of their hosts as an obligate component of infection. This physical association is usually mediated by adhesins elaborated from the bacterial surface, including pili, flagella, other externalized proteins and exopolysaccharides (Berne *et al*., 2018). Adhesins can also drive bacterial attachment to abiotic surfaces. On both abiotic surfaces and host tissues, adherent cells may assemble into multicellular biofilms, physically and chemically recalcitrant communities that can protect the colonizing bacterial population (Flemming *et al*., 2016). Biofilm formation is driven by a combination of cell-to-surface interactions, intercellular cohesion, and extrusion of exopolymeric materials to build an extracellular matrix.

We discovered a polysaccharide-based adhesin in *Agrobacterium tumefaciens*, a member of the Alphaproteobacteria (APB) and causative agent of crown gall disease on a wide variety of dicotyledonous plants (Gelvin, 2003). Crown gall is a neoplastic disease that results from the transfer of a large segment of DNA (T-DNA) from the bacteria into plant cells at the site of infection via the activity of a type IV secretion system (Li & Christie, 2018). This interkingdom gene transfer process, both its regulation and the mechanism of transfer, has been the subject of intense study and T-DNA transfer has been extensively engineered for the generation of transgenic plants. There is, however limited, understanding of the initial colonization events that occur on the tissue surface, both for benign interactions and those that lead to the initiation of T-DNA transfer (Fuqua, 2008).

Micrographs dating as early as the 1940s suggest that *A. tumefaciens* often attaches to surfaces via a single cellular pole (Braun & Elrod, 1946, Tomlinson & Fuqua, 2009). Surfaces to which *A. tumefaciens* can attach include plant tissues and cells, other bacterial cells to form multicellular rosettes, and abiotic materials. We discovered that *A. tumefaciens* produces an adhesive structure at one of its poles that is predominantly comprised of polysaccharide, and hence we designated this the unipolar polysaccharide (UPP) (Tomlinson & Fuqua, 2009). Fluorescently-tagged lectins including wheat germ agglutinin (WGA) and *Dolichos biflorus* agglutinin (DBA) label the UPP as a small fluorescent focus at the pole that is in contact with abiotic surfaces, plant tissues, and other bacterial cells (Fig. 1) (Xu *et al*., 2012, Xu *et al*., 2013). The UPP structure shares facile and functional similarity to the adhesive holdfast localized to the tip of the cellular stalk for the prosthecate APB *Caulobacter crescentus* (Poindexter, 1981). In fact, production of the UPP is coordinated with the biphasic life cycle of *A. tumefaciens* in a similar fashion as the holdfast in *C. crescentus*, and is produced by mother cells, that are often adhered to surfaces, and will generate multiple progeny daughter cells via a polar budding mechanism (Brown *et al*., 2012). In both bacteria, production of the unipolar adhesin is stimulated by surface contact. In *C. crescentus*, this accelerates holdfast production to earlier in the cell cycle, but in *A. tumefaciens* surface contact is required for cells to produce the UPP (Li *et al*., 2012). The smaller size and strict surface-contact dependence of UPP production led to it being overlooked for many years, relative to the more microscopically conspicuous holdfast of *C. crescentus*. Several *A. tumefaciens* mutants have been characterized which exhibit a bypass of surface-contact dependence and elevated UPP production due to increased levels of the cytoplasmic second messenger cyclic diguanylate monophosphate (cdGMP) (Feirer *et al*., 2017, Feirer *et al*., 2015, Xu *et al*., 2013).

**Figure 1.**
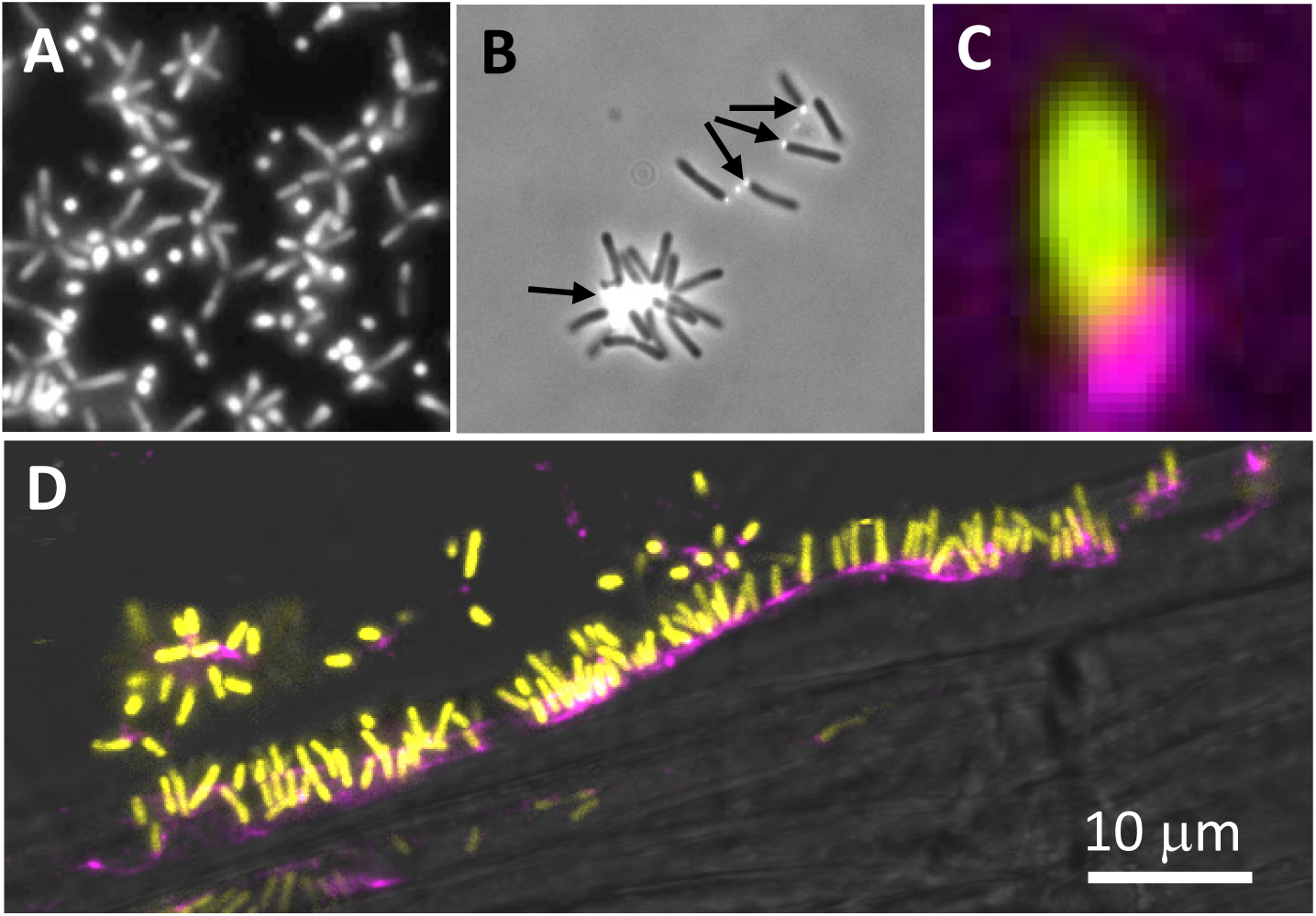
UPP-mediated polar attachment in *A. tumefaciens*. *A. tumefaciens* UPP on abiotic surfaces and plant tissues. Fluorescence microscopy using wheat germ agglutinin lectin labeled with FTIC (fl-WGA) and Alexafluor 594 (af-WGA). (A-C) Attachment assays on 22 x 22 mm glass coverslips. *A. tumefaciens* C58 cells incubated for the indicated time in ATGN minimal medium with coverslips prior to lectin labeling with 10 μg/ml of either fl-WGA or af-WGA for 20 min. (A) fl-WGA labeling of a 12 hr biofilm. Merged image from bright field and fluorescence microscopy. (B) af-WGA-labeling of a short-term attachment assay after 2 hrs incubation. Merged image from phase contrast and fluorescence. (C) af-WGA labeling of a single *A. tumefaciens* cell attached to glass expressing plasmid-borne *P_tac_*::*gfpmut3* (pJZ383). Scanning Disk Laser Confocal fluorescence microscopy; Merged green and red channels, pseudo-colored yellow and magenta, respectively. (D) Binding of *A. tumefaciens* (pJZ383) to *Arabidopsis thaliana* WS 1 cm root cutting incubated for 3 hrs in ATGN minimal medium. Merged green and red channels pseudo-colored yellow and magenta, respectively plus bright field. Microscopes: Nikon E800 fluorescence microscope and Photometrics Cascade cooled CCD camera; Spinning Disk Nikon TE2000U microscope Yokagawa CSU10 confocal scanner unit. Fluorescence settings: fl-WGA and GFP, excitation, 460-500 nm, emission 510-560 nm; af-WGA, excitation 510-560 nm, emission >610 nm.

A cluster of six predicted polysaccharide biosynthetic genes (ATU_RS06125-RS06100) was initially identified based on adhesion defects of a small number of mutants in *A. tumefaciens* (Fig. 2A) (Xu *et al*., 2012). These genes are homologous to six genes in a syntenic cluster required for synthesis of a glucomannan involved in recognition of host legumes during formation of symbiotic nodules in the related symbiotic nitrogen-fixing APB *Rhizobium leguminosarum* bv. *viciae* (Williams *et al*., 2008). Deletion of the entire six-gene cluster in *A. tumefaciens* led to complete loss of biofilm formation on abiotic surfaces and abolished lectin labeling of the UPP (Xu *et al*., 2012). In-frame deletion of a single gene in this cluster (ATU_RS06105) encoding a polyisoprenylphosphate hexose-1-phosphate transferase (PHPT) homologous to the HfsE glycosyltransferase from *C. crescentus* required for holdfast synthesis (Toh *et al*., 2008), also abolished surface attachment and visible UPP production. In addition to this PHPT protein, the other five genes in this cluster encode homologs to polysaccharide biosynthesis genes (Table S1), and we have tentatively designated this gene cluster as *uppABCDEF* (Thompson *et al*., 2018). In addition to the PHPT protein UppE, several of these genes encode components of a Wzx-Wzy-type polysaccharide pathway (Cuthbertson *et al*., 2009). Using the general polysaccharide biosynthetic nomenclature, UppE encodes a WbaP initiating glycosyltransferase, UppB (ATU_RS06120) encodes a predicted Wzb-type <u>p</u>eriplasmic <u>c</u>o-<u>p</u>olymerase (PCP), and UppC (ATU_RS06115) encodes a putative Wza-type outer membrane polysaccharide secretin (OPX). The other three genes are not restricted to Wzx-Wzy systems, but are still predicted polysaccharide biosynthesis genes (Table S1); UppA (ATU_RS06125) encodes a putative polysaccharide acetyltransferase with similarity to BcsL involved in bacterial cellulose synthesis, UppD (ATU_RS06110) encodes a Type 4 glycosyltransferase (GT4), and UppF (ATU_RS06100) is similar to WaaL O-antigen ligases that often transfer polysaccharides to the core of lipopolysaccharide (LPS), in formation of the O-antigen or LPS-conjugated polysaccharides (Romling & Galperin, 2015, Ruan *et al*., 2018).

**Figure 2.**
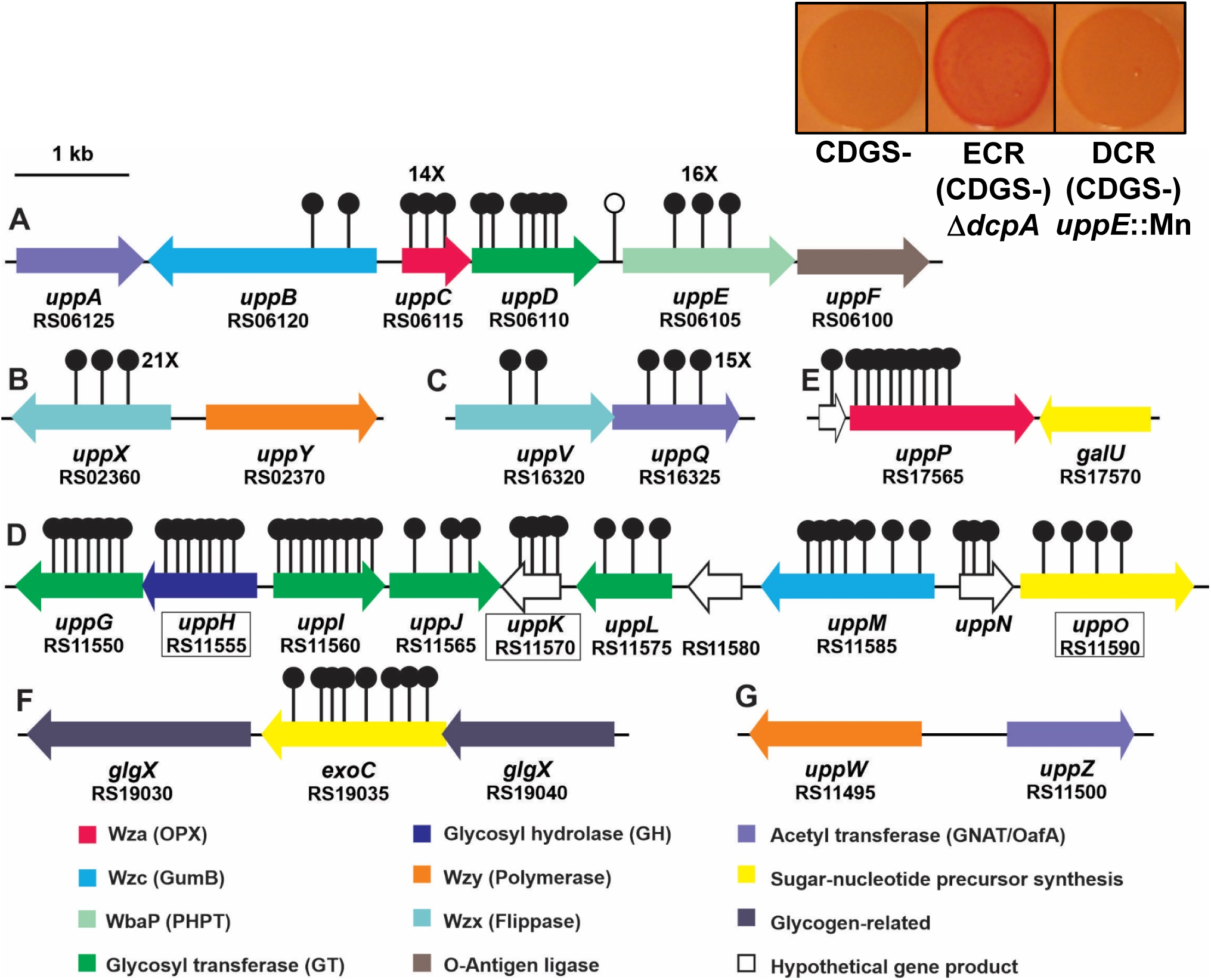
UPP biosynthesis gene clusters and Congo red colony phenotypes in *A. tumefaciens*. Gene clusters important for UPP biosynthesis that were identified via transposon screens for mutants with decreased Congo red (DCR) phenotypes. Stemmed circles represent independent, sequenced transposon insertion sites. Letters A-G designate specific genetic loci. The single unfilled marker upstream of *uppE* represents a transposon mutant isolated based on a biofilm deficiency (Xu *et al*., 2012). Numbers with X indicate 14 or more independent transposon insertions. Boxed gene names and designations indicate genes for which in-frame-deletions in wild type do not lead to attachment deficiencies. Predicted functions based in sequence similarity are described in the key below the gene map and are color-coded. Inset boxes on the upper right provide CR phenotypes of *A. tumefaciens* strains lacking all the known dispensable exopolysaccharides except UPP (CDGS-), the same strain elevated for cdGMP via a *dcpA* deletion and thus exhibiting an elevated Congo red (ECR) phenotype (middle), and the same strain with a Mariner transposon insertion in *uppE* conferring a DCR phenotype.

The *upp* gene cluster is completely or partially conserved among diverse members of the Rhizobiales, and mutations in several of these homologs have been shown to impact production of a unipolar polysaccharide and rosette formation in *Rhodopseudomonas palustris* (Fritts *et al*., 2017). Thus, in at least three members of the Rhizobiales, homologs of the *A. tumefaciens uppA-F* genes are required for synthesis of polar polysaccharides, and it seems likely that this function is conserved in these bacteria. However, notably absent from these gene clusters are homologs of the Wzx-type polysaccharide flippase enzymes and the Wzy-type polysaccharide polymerases, two of the defining components of these pathways (Cuthbertson *et al*., 2009). In this study, we use several distinct forward genetic approaches along with targeted mutational analysis to expand and deepen our prior analyses on *A. tumefaciens* UPP biosynthesis, identifying over 16 additional genes in 6 genomic clusters that are required for UPP production and attachment to surfaces. We combine this genetic analysis with microscopy and functional assays to reveal that there are at least two distinct polysaccharides that both contribute to the *A. tumefaciens* UPP, and that the biosynthetic pathways are genetically separable, but also overlapping and co-regulated.

## Results

### Mutations in the *uppABCDEF* gene cluster disrupt attachment and UPP production

While performing a biofilm deficiency screen, we initially isolated a Mariner (*Himar1)* transposon mutant of *A. tumefaciens* C58 that was completely deficient for biofilm formation on PVC coverslips relative to wild type (Xu *et al*., 2012). The transposon in this mutant was immediately upstream of the gene we now designate as *uppE* (Fig. 2A, unfilled transposon marker). Creation of a mutant with an in-frame deletion of *uppE* (ATU_RS06105) resulted in an equivalent loss of biofilm formation and WGA labeling as in the transposon mutant (Figs. 3, 4). To test the other genes in the *uppABCDEF* cluster (ATU_RS06125-RS06100), we created in-frame deletions of each gene individually and observed significant decreases in biofilm formation on PVC coverslips (Fig. 3, Table S1). The *uppC* and *uppE* deletions exhibited the most profound defects, but the *uppA*, *uppB*, *uppD* and *uppF* deletions also were substantially decreased biofilm formation, although still detectable. Short term attachment assays with cells incubated on PVC coverslips revealed weak attachment in all mutants and loss of labeling with Alexafluor594-tagged WGA (af-WGA) for all of the mutants (Fig. 4, Table S1). Attachment and biofilm formation deficiencies of deletion mutants were fully complemented by ectopic expression with plasmid-borne copies of these genes, with the exception of the *uppC* mutant, which was only partially complemented with the *P_lac_-uppC* plasmid (Fig. 3). We realized that the *uppC* deletion also impacted the overlapping downstream *uppD* coding sequence, and provision of a plasmid expressing both *uppC* and *uppD* fully reversed the Δ*uppC* mutant phenotype. As described above, the six *upp* genes are all homologs of polysaccharide biosynthetic functions (Table S1). However, it was clear that based on their predicted functions in a Wzx-Wzy-type pathway, they would not be sufficient and that there must be additional genes required for UPP synthesis. Most notably, homologs of both the Wzx flippase and the Wzy polymerase are not found in this gene cluster nor in proximal genes.

**Figure 3.**
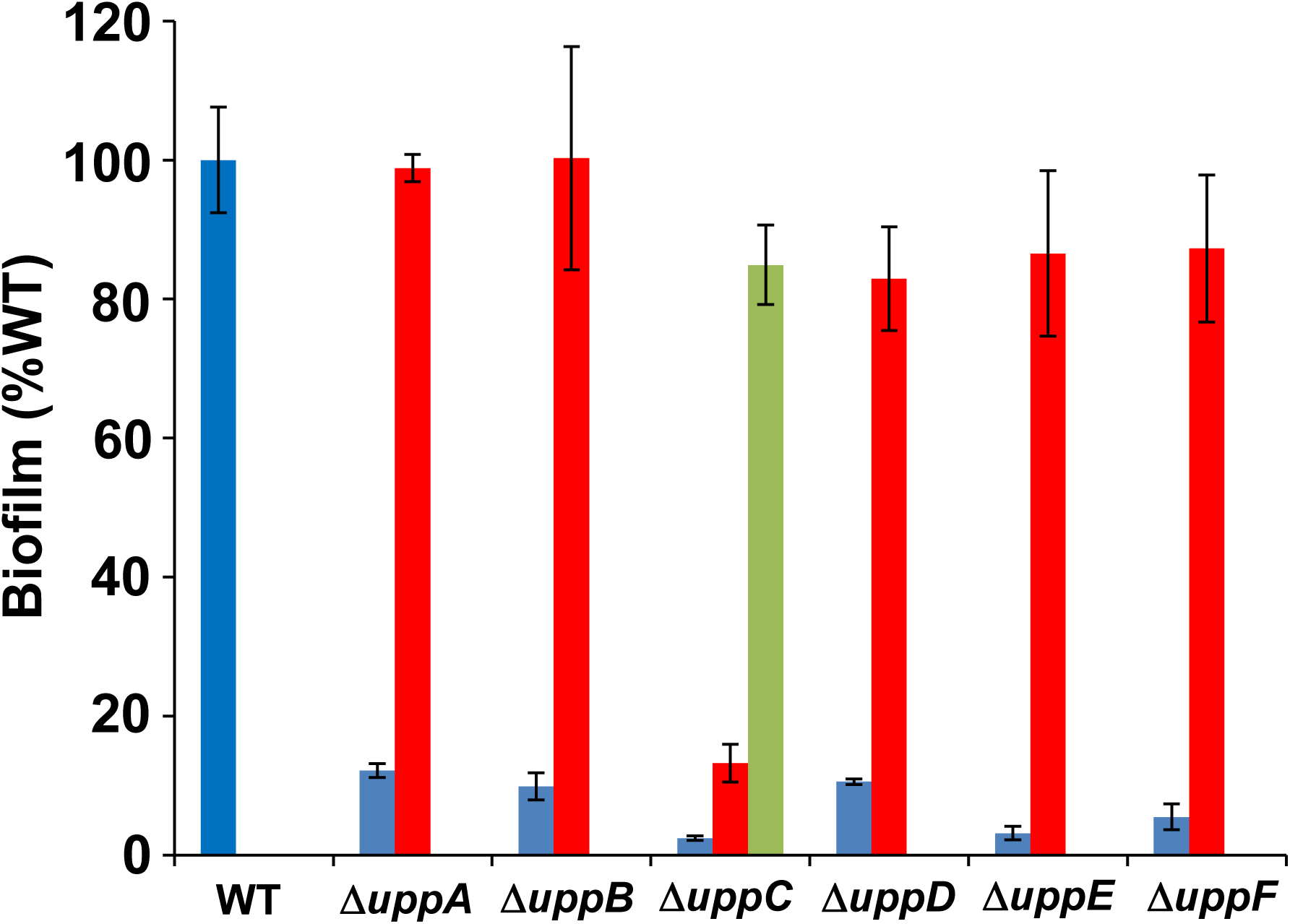
Biofilm phenotypes of *uppA-F* mutants. Ratio of acetic acid-solubilized CV absorbance (A_600_) from 48 h biofilm assays normalized to the OD_600_ turbidity measured from the planktonic phase of the same culture. Ratio for wild type set to 100% and mutant derivatives expressed as percent wild type. Blue bars are the strains harboring the pSRK-Gm vector, red bars are expressing pSRK-Gm derivatives with the corresponding gene, and the single green bar is the Δ*uppC* strain expressing a pSRK-Gm derivative with *uppC* and *uppD*. All assays were performed with 500 μM IPTG. Bars represent the mean of triplicate assays and error bars are standard deviation.

**Figure 4.**
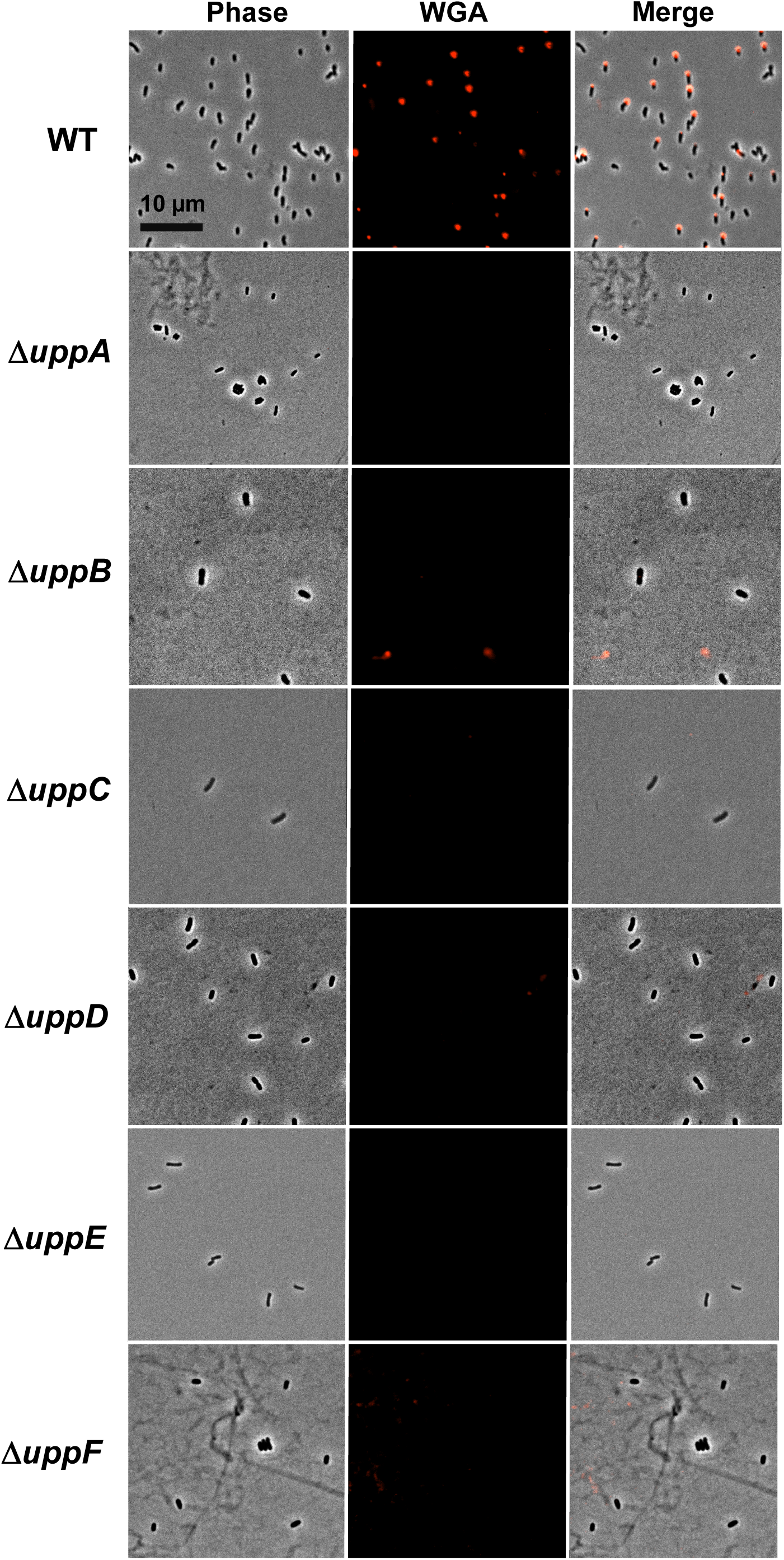
af-WGA labeling of UPP in *uppA-F* mutants. Short term binding assays of *A. tumefaciens* C58 derivatives on glass coverslips stained with af-WGA. Phase images, fluorescence images (510-560 nm excitation, emission >610 nm), and merged images. Microscopy was performed as described in experimental procedures on a Nikon Eclipse Ti-E microscope with a Nikon E800 camera with the 60X oil immersion objective and NIS-Elements Software, and analyzed with Fiji (Schindelin *et al*., 2012).

### A facile genetic screen for UPP biosynthetic genes

In prior studies, we had identified a set of mutants in four different loci (*visR* ATU_RS02585, *claR* ATU_RS08010*, pruA* ATU_RS05580, and *dcpA* ATU_RS16190), that manifested elevated production of the UPP coincident with elevated cellular cdGMP levels (Feirer *et al*., 2017, Feirer *et al*., 2015, Xu *et al*., 2013). The screen was based on the polysaccharide-binding, azo-dye Congo red (CR, which can also bind to β-amyloid proteins), and in all cases elevated CR pigmentation correlated with increased UPP production and hyper-adherence to surfaces. Wild type *A. tumefaciens* forms very pale orange colonies when cultured on agar plates of a standard minimal medium (ATGN) supplemented with CR (ATGN-CR) (Fig. 2). Elevated production of either the UPP or cellulose results in red pigmentation of bacterial colonies grown on ATGN-CR, which we designated the ECR (<u>E</u>levated <u>C</u>ongo <u>R</u>ed) phenotype (Xu et al., 2013). In an *A. tumefaciens* mutant disabled for cellulose production (Cel-), the ECR phenotype reflects UPP production. In the current study, we utilized this same CR-based screen, but performed it with the four different ECR mutants, each with in-frame deletions in the relevant regulatory genes (Δ*visR*, Δ*claR,* Δ*pruA*, and Δ*dcpA*). These strains were also genetically disabled for other known dispensable polysaccharides (<u>c</u>ellulose, cur<u>d</u>lan, β,1-2 <u>g</u>lucans, <u>s</u>uccinoglycan) but retain UPP production (UPP+); for ease of description we generally designate these strains as ECR (CDGS-) derivatives (the specific deleted regulator resulting in the ECR phenotype is indicated as required).

Mariner (*Himar-1*) transposon mutant libraries of each strain were screened for colonies with a notable decrease in CR pigmentation (the DCR phenotype), reasoning that we would likely isolate mutations in UPP biosynthetic functions, as well as positive regulatory elements required for the ECR phenotype. Overall, we screened close to 50,000 colonies of these derivatives for the DCR phenotype. Once confirmed as DCR mutants, the transposon insertion site junctions were amplified and sequenced. From this initial screen 140 independent mutants were identified in 19 discrete annotated genes (Fig. 2, Table S2), most predicted to be involved in polysaccharide biosynthesis or clustered with such genes (other DCR mutants were in known or predicted regulatory functions, see Xu et al., 2013). All but one of the 19 disrupted loci were identified multiple times, and mutations in all but two genes were found in more than one of the mutants used in the screen, with 9 genes identified in at least three different mutant backgrounds (Table S2). Subsequent analyses revealed that most of the genes disrupted by Mariner insertions in the DCR mutants, when deleted and retested in new genetic backgrounds, resulted in decreased or absent UPP production and decreased biofilm formation (Fig. S1, Table S1). Of the genes identified, four were in the previously characterized *uppA-F* cluster (number of mutants); *uppB* (2)*, uppC* (14), *uppD* (6), and *uppE* (16) for a total of 38 (27%) of the independent mutants (Fig. 2A). No DCR mutations were identified in *uppA* and *uppF.* The identification of four of the genes in this cluster however, validated the ability of our DCR screen to detect defects in UPP biosynthesis.

### Identification of two Wzx flippase homologs that impact UPP production

One of the required Wzx-Wzy pathway functions that had not been identified previously was a flippase (Wzx) homolog. The DCR screen identified two different Wzx-type genes (Fig. 2B, 2C; Tables S1, S2). In all four mutant backgrounds used in the screen, 21 independent mutations were obtained in a Wzx-type flippase homolog, ATU_RS02360, that we now designate *uppX*. This gene encodes a 426 aa protein, with 10-12 predicted transmembrane helices, as is consistent with many flippase homologs (Hong *et al*., 2018). In-frame deletion of *uppX* in the wild type C58 strain resulted in a dramatic loss of biofilm formation (Fig. 5A) and attachment, as well as UPP production as judged by WGA and DBA labeling (Fig. 5B). These phenotypes were fully complemented by expression of a plasmid-borne copy of *uppX*. Divergently oriented from *uppX* is ATU_RS02370 (Fig. 2B), encoding a Wzy-type protein predicted to function as a polysaccharide polymerase, but no DCR transposon mutants were obtained in this gene. Interestingly, DCR mutations in a second flippase homologue were also identified, ATU_RS16320, now designated *uppV* (Fig. 2C, Table S1). UppV is a 477 aa Wzx-type protein, also with 12 predicted transmembrane domains. Only two mutations in *uppV* were isolated, but one each in the Δ*pruA* and Δ*dcpA* mutants (Table S2). Deletion of *uppV* in wild type C58 significantly decreased but did not abolish biofilm formation relative to wild type (Fig. 5A), although weak attachment and no substantial WGA lectin binding was observed (Fig. 5B).

**Figure 5.**
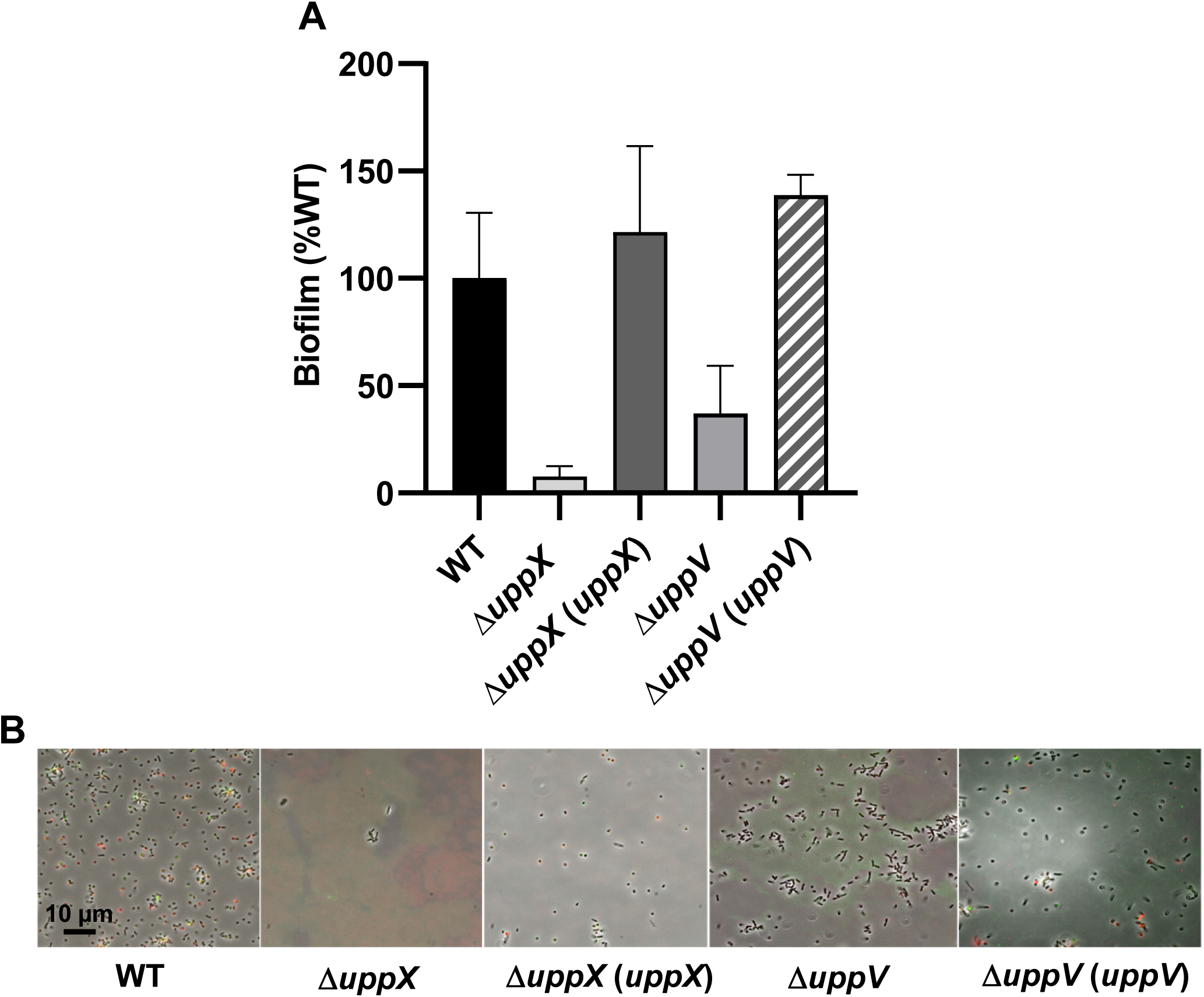
Biofilm and lectin-binding phenotypes of putative flippase deletion mutants. (A) Biofilm assays performed as in Fig. 3 for flippase deletion mutants and complementation strains. WT biofilm was set to 100%. Plasmid-borne copies of genes were expressed from the *P_lac_* promoter in pSRK derivatives induced with 500 μM IPTG. Bars represent the mean of triplicate assays and error bars are standard deviation. (B) Short term attachment assays as in Fig. 4, co-incubated with af-WGA (red) and fl-DBA (green). Only merged images shown. Microscopy as described in Fig. 4. af-WGA (excitation 510-560 nm, emission >610 nm); fl-DBA (excitation 460-500 nm, emission 510-560 nm).

The *uppV* coding sequence terminates 8 bp upstream of another reading frame, ATU_RS16325, that we have designated *uppQ* (Fig. 2C). Fifteen independent DCR mutants were isolated with insertions in *uppQ*, which is oriented convergently to an adjacent unidentified reading frame (ATU_RS16330), strongly suggesting that *uppQ* itself is required for UPP production. UppQ is homologous to acetyltransferases of the GNAT family of enzymes (Vetting *et al*., 2005), and paralogous to UppA (ATU_RS06125) (Table S1). Deletion of *uppQ* in the wild type results in loss of WGA lectin binding and has a strong biofilm defect (Fig. S1, Table S1) suggesting it plays a crucial role in UPP biosynthesis.

### A gene cluster of multiple polysaccharide biosynthetic genes with roles in UPP biosynthesis and biofilm formation

Strikingly, 47 of the DCR mutants (33.6%) had transposon insertions in a single gene cluster ATU_RS11550-RS11590, with mutations represented in all four screening backgrounds, and multiple independent mutations identified for all of these genes (Fig. 2D). Seven of these annotated genes have predicted polysaccharide-related functions, and the others genes encode hypothetical functions. Nine out of ten genes in this contiguous cluster were identified in the DCR screen, with the exception of ATU_RS11580, a hypothetical gene (Fig. 2D). The genes are organized into at least 4 different predicted transcription units. Multiple lines of evidence for most of these genes indicate roles in UPP biosynthesis and hence we have designated them *uppG* through *uppO* (Fig. 2D, Fig. S1, Table S1). Four of these genes are predicted to encode cytoplasmic glycosyl transferases (GTs) *uppG* is in the Carbohydrate Active Enzymes (CAZy) group GT4, *uppI* and *uppJ* in CAZy group GT2, and *uppL* (CAZy group 26) (Lombard *et al*., 2014). The *uppH* gene encodes a predicted periplasmic Class 10 glycoside hydrolase, *uppM* encodes a periplasmic co-polymerase (PCP-2b), and *uppO* encodes a phosphomannomutase. Two of the genes interrupted among the DCR transposon mutants, *uppK* and *uppN* are predicted to encode proteins of unknown function, and in fact the short 59 amino acid coding sequence we designate as *uppN* is not annotated as a gene in the current *A. tumefaciens* genome annotation (although it was designated as Atu2378 in an earlier genome annotation).

Independent in-frame deletions were generated in all of these predicted genes in the wild type C58 background, and tested for biofilm formation defects. Profound defects were observed for Δ*uppG*, Δ*uppI*, Δ*uppJ*, Δ*uppL*, Δ*uppM*, and Δ*uppN* (Fig. S1 and Table S1). Ectopic expression of the disrupted gene complemented the deletion mutants for *uppG*, *uppI*, *uppJ*, and *uppL* (Table S1, Fig. S2A). In frame deletion of *uppH* did not have an impact on attachment, but the gene overlaps by 4 bp with the *uppG* coding sequence and multiple lines of evidence suggest that the *uppH* transposon mutations are polar on *uppG*. However, subsequent analysis revealed an important activity of UppH relevant to the UPP (described below).

In-frame deletion of the *uppN* gene with a 59 aa translation product resulted in a dramatic loss of biofilm formation, but complementation did not effectively restore this phenotype (Fig. S1A, S2A). Initially, we thought this might be due to polarity on the phosphomannomutase *uppO*, although it is 155 bp downstream of the *uppN* gene (Fig. 2D). However, in frame deletion of *uppO* did not result in a biofilm defect in a wild type background and seemed to result in greater adherence (Fig. S1A), despite the isolation of four DCR transposon mutants with insertions in this gene. Thus, any unexpected polarity on *uppO* apparently does not explain the Δ*uppN* phenotype, and its weak complementation. Similar to *uppO*, in-frame deletion of the hypothetical gene *uppK* did not impact biofilm formation, in the wild type (Fig. S1A). We hypothesized that the difference might be due to the elevated cdGMP in the mutants used for DCR screening, whereas the *uppK* deletion was in the wild type. However, introduction of the Δ*pruA* deletion that dramatically increases cdGMP and in which we isolated all 4 *uppK* transposon mutants, increased adherence in the wild type, and this was not diminished by the Δ*uppK* mutation (Fig. S2b).

### Mutations in *uppM* and *uppB* have additive defects

The UppM protein is homologous to Type 2b periplasmic co-polymerases (PCP-2b), and is paralogous to UppB. In-frame deletion of *uppM* in the wild type resulted in a significant, yet incomplete loss of biofilm formation (Fig. S1A, Table S1). Given the similarity of UppM and UppB, we speculated that a functional *uppB* gene might partially mask the full attachment defect, as the Δ*uppB* mutant also exhibits an incomplete loss of biofilm formation (Fig. 3). A mutant with both *uppB* and *uppM* deleted shows a complete loss of biofilm formation (Fig. S2C), lower than either mutation alone, consistent with their similar roles in UPP biosynthesis.

### The UppG glycosyl transferase catalytic site is required for full UPP synthesis

Careful examination of the Δ*uppG* mutant revealed that it does in fact bind to the WGA lectin, but that the signal is significantly weaker than the wild type (Fig. S3A, S3B). We recreated this deletion in the Δ*pruA* CDGS-screening strain, and found it to be fully complemented by a plasmid-borne copy of the gene as evaluated by CR colony staining (Fig. S3C). GT4-Class glycosyltransferases similar to UppG have a conserved DXD catalytic motif (Breton *et al*., 2006). Mutation of the predicted DXD active site motif (pos. 142-4) at the first D residue (DXD>AXD) abolished complementation (Fig. S3C), confirming a requirement for its GT activity in this and related phenotypes.

### A predicted outer membrane protein required for UPP synthesis

Nine independent DCR mutants had transposon insertions within or immediately upstream of the ATU_RS1765 gene and we have designated this gene as *uppP* (Fig. 2E). One of the nine transposon insertion mutants mapped to the short annotated 106 aa upstream hypothetical coding sequence ATU_RS1760 which terminates 6 bp upstream of the predicted *uppP* start codon. This single transposon insertion is likely polar on *uppP*. UppP is 497 aa in length, has an N-terminal secretion signal and encodes an outer membrane β-barrel-type II porin protein (Table S1). UppP is much larger than the predicted polysaccharide secretin UppC (190 aa), does not match well with Wza homologues, and unlike UppC, lacks a lipidation motif. The *uppP* gene is convergently oriented to a GalU homologue (ATU_RS17570), predicted to encode a UTP-glucose-1-phosphate uridylyltransferase, that could synthesize a UDP-glucose precursor, but no DCR mutants were isolated that had insertions in this gene (although it is implicated by other findings, described below). An in-frame deletion of *uppP* in wild type resulted in complete loss of biofilm formation and was fully complemented by plasmid-borne expression of the gene (Fig. S1A, Table S1). The Δ*uppP* mutant does not produce UPP as evaluated with af-WGA staining (Fig. S1B).

### Mutations in the *exoC* gene result in the DCR phenotype

Eight independent DCR mutants were found to have transposon insertions within ATU_RS19035 (Fig. 2F), encoding a gene previously designated as *exoC* (Kamoun *et al*., 1989). The *exoC* gene product encodes a predicted phosphoglucomutase that would catalyze conversion of glucose-6-phosphate to glucose-1-phosphate, a precursor of sugar nucleotide substrates for polysaccharide biosynthesis. Mutants in *exoC* have been reported to be pleiotropic with defects in succinoglycan production, motility and loss of virulence on plants (Cangelosi *et al*., 1987, Kamoun *et al*., 1989). An *exoC Himar1* transposon mutant independently isolated in an otherwise wild type background harboring a plasmid-borne copy of the diguanylate cyclase *pleD* (ATU_ RS06385) revealed this mutant to be unable to produce UPP or cellulose (as judged by CR staining) and deficient for lectin labeling when induced with IPTG (Fig. S4A, S4B). The *exoC* mutant was also deficient for the exopolysaccharide succinoglycan and was non-motile (Fig. S4C, D). The *exoC* gene in *A. tumefaciens* is located within an apparent operon involved in glycogen biosynthesis and utilization (Fig. 2F). However, none of the other genes in this cluster, other than *exoC,* were isolated in the DCR screen.

### Apparent redundancy for Wzy-type polymerase function(s)

The DCR screen described above revealed a large number of additional genes required for UPP biosynthesis. Despite the wide range of polysaccharide-related functions identified, it was surprising that no DCR mutants were isolated with mutations in Wzy-type polymerase homologs, despite the divergent orientation from *uppX* of a gene encoding a Wzy-type protein (ATU_RS02370) (Fig. 2B). We reasoned that the crucial Wzy polymerase activity required for a Wzx-Wzy-type pathway, might be fulfilled by functionally redundant proteins. Examination of the C58 genome revealed a Wzy-type protein paralogous to ATU_RS02370, ATU_RS11495 to be the closest match, (37% identity, 57% similar). As described below, we have designated these genes as *uppY* (ATU_RS02370) and *uppW* (ATU_RS11495) (Fig. 6A). Single, in-frame deletions of *uppY* and *uppW* did not result in any discernable change in CR colony pigmentation nor for adhesion based on *in vitro* biofilm assays (Fig. 6B, 6C). However, combination of the *uppY* and *uppW* deletions resulted in the DCR phenotype in the Δ*pruA* CDGS-mutant (Fig. 6B) and abolishment of biofilm formation in an otherwise wild-type strain (Fig. 6C). Expression of plasmid-borne copies of either gene complemented the Δ*uppY* Δ*uppW* mutant for adhesion. UppY and UppW are both Wzy-type RfaA homologs with 10 predicted transmembrane domains, that could function as polysaccharide polymerases in the periplasm. Our observations suggested that either UppY or UppW can compensate for the absence of its paralog, for CR interactions and for adhesion in our *in vitro* assays. Initially, the explanation for this functional redundancy was unclear, but we now recognize that each protein plays a specific role in forming the UPP adhesin.

**Figure 6.**
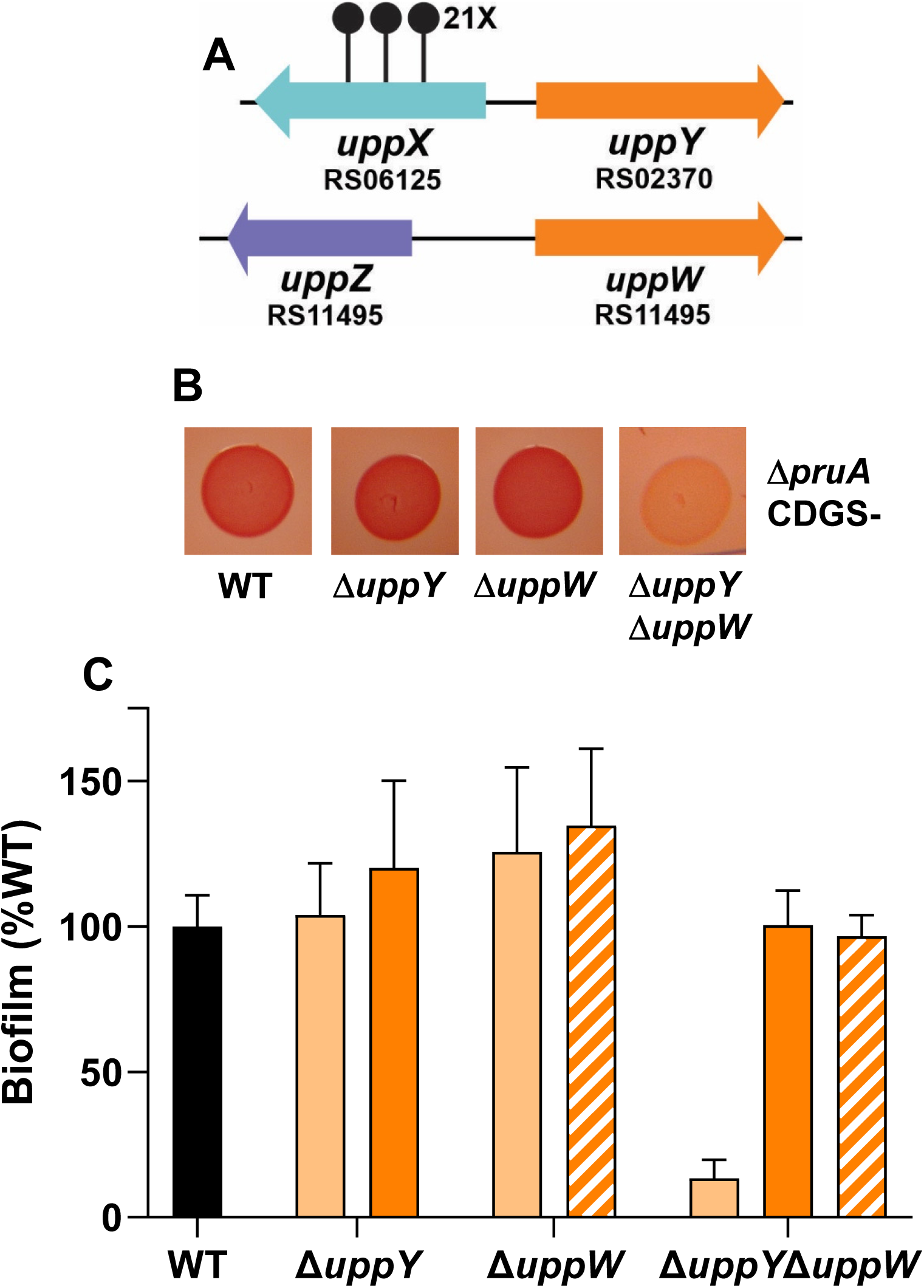
Two Wzy-type polymerase proteins are redundant for attachment and biofilm formation. (A) Genetic loci of *uppY* and *uppW*. Stalked circles represent independent, sequenced transposon insertion sites and numbers with X indicate 14 or more independent transposon insertions. (B) ATGN CR colony phenotypes of the indicated deletion mutants in the Δ*pruA* CDGS-background. (C) Quantitative biofilm assay as performed in Fig. 3 with the ratio of A_600_ of acetic acid-solubilized CV-stained coverslips to planktonic OD_600_ normalized to percent wild type for the indicated in-frame deletion mutants in otherwise wild type C58 after 48 h incubation in ATGN. The indicated strains with solid black and light orange bars have the pSRK-Gm vector control, whereas the strains represented by dark orange bars harbor a *P_lac_*-*uppY* plasmid and the strains with cross hatched bars harbor the *P_lac_*-*uppW* plasmid. Cultures grown in 400 μM IPTG to induce *P_lac_*. Error bars are standard deviation of assays performed in triplicate.

### The UPP adhesin is composed of two chemically and genetically distinct polysaccharides

The large number of putative UPP genes we identified from the DCR screen was striking given an apparent duplication of function for many of the requisite steps involved in Wzx-Wzy-type polysaccharide synthesis (Fig. 2, Table S1). Previously, *A. tumefaciens* UPP synthesis was examined using multiple lectins with different oligosaccharide specificities (Xu *et al*., 2012). In addition to GlcNAc residues recognized by WGA, the lectin from the legume *Dolichos biflorus* (DBA) has specificity for *N*-acetylgalactosamine residues (GalNAc) and effectively labels the *A. tumefaciens* UPP. We performed dual labeling experiments with af-WGA and fl-DBA of *A. tumefaciens* (Δ*pruA* CDGS-) cells adhered to coverslips. Both lectins labeled the poles for a large proportion of adhered cells (Fig. 7). The pattern of labeling was striking with the fl-DBA fluorescence forming a tight focus at the point of contact, and the af-WGA labeling more diffusely bound in a second ring around the DBA-labeled sector. This suggested that there were two discrete populations of polysaccharide at the pole, one identifiable using the GlcNAc-specific WGA and a second labeled with the GalNAc-specific DBA. We tentatively designated these two polysaccharide species as UPP_GlcN_ and UPP_GalN_, respectively. The basis for the clear spatial segregation of the lectin labeling pattern was initially unknown.

**Figure 7.**
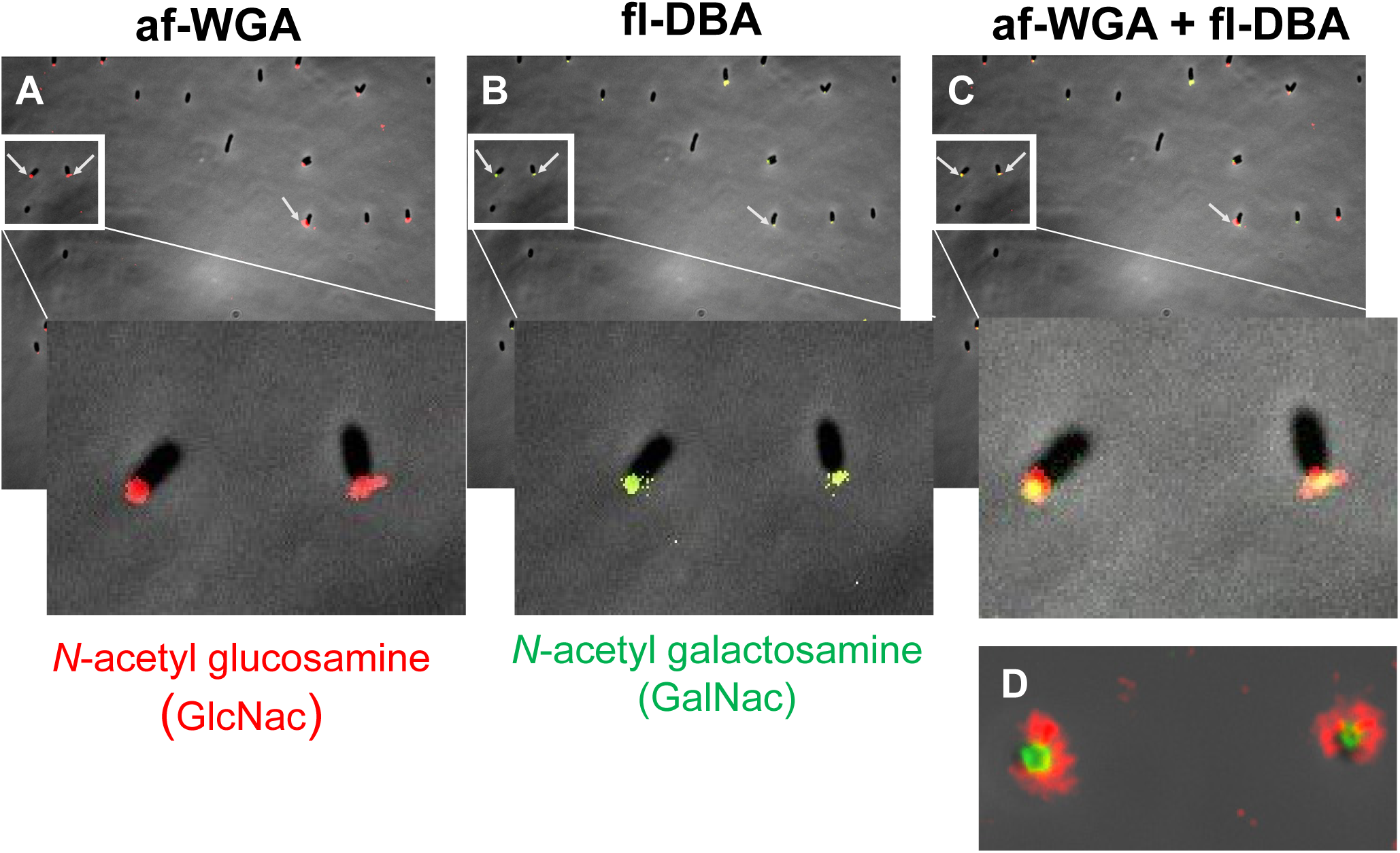
Dual lectin binding of wild type UPP with WGA and DBA. Short term attachment assay on a glass coverslip co-incubated with lectins. (A) UPP stained with af-WGA showing red fluorescent polar foci. (B) UPP stained with fl-DBA showing green fluorescent polar foci. (C) UPP stained with both af-WGA and fl-DBA showing the sectored pattern of dual red and green fluorescence. Arrows indicate clearly labeled UPP. Insets – magnification of boxed area. (D) Image of two attached dual labeled cells at the focal plane of surface contact. Microscopy as in Fig. 5.

Our findings suggested that the two Wzy-type proteins, UppY and UppW, were functionally redundant for adhesion and for CR labeling. Lectin labeling of the Δ*uppY* and Δ*uppW* mutants background revealed reciprocal specificity for each polymerase and a corresponding polysaccharide (Fig. 8). In short-term coverslip binding assays probed simultaneously with both lectins, the Δ*uppY* mutation caused loss of all fl-DBA labeling, but did not affect that of af-WGA, whereas the Δ*uppW* mutation caused loss of the af-WGA labeling but retained that of fl-DBA. The double mutant, which abolished elevated CR binding and biofilm formation, was unable to bind either lectin. This suggested that production of the DBA-labeled UPP_GalN_ requires UppY, whereas elaboration of the WGA-labeled UPP_GlcN_ requires the function of UppW.

**Figure 8.**
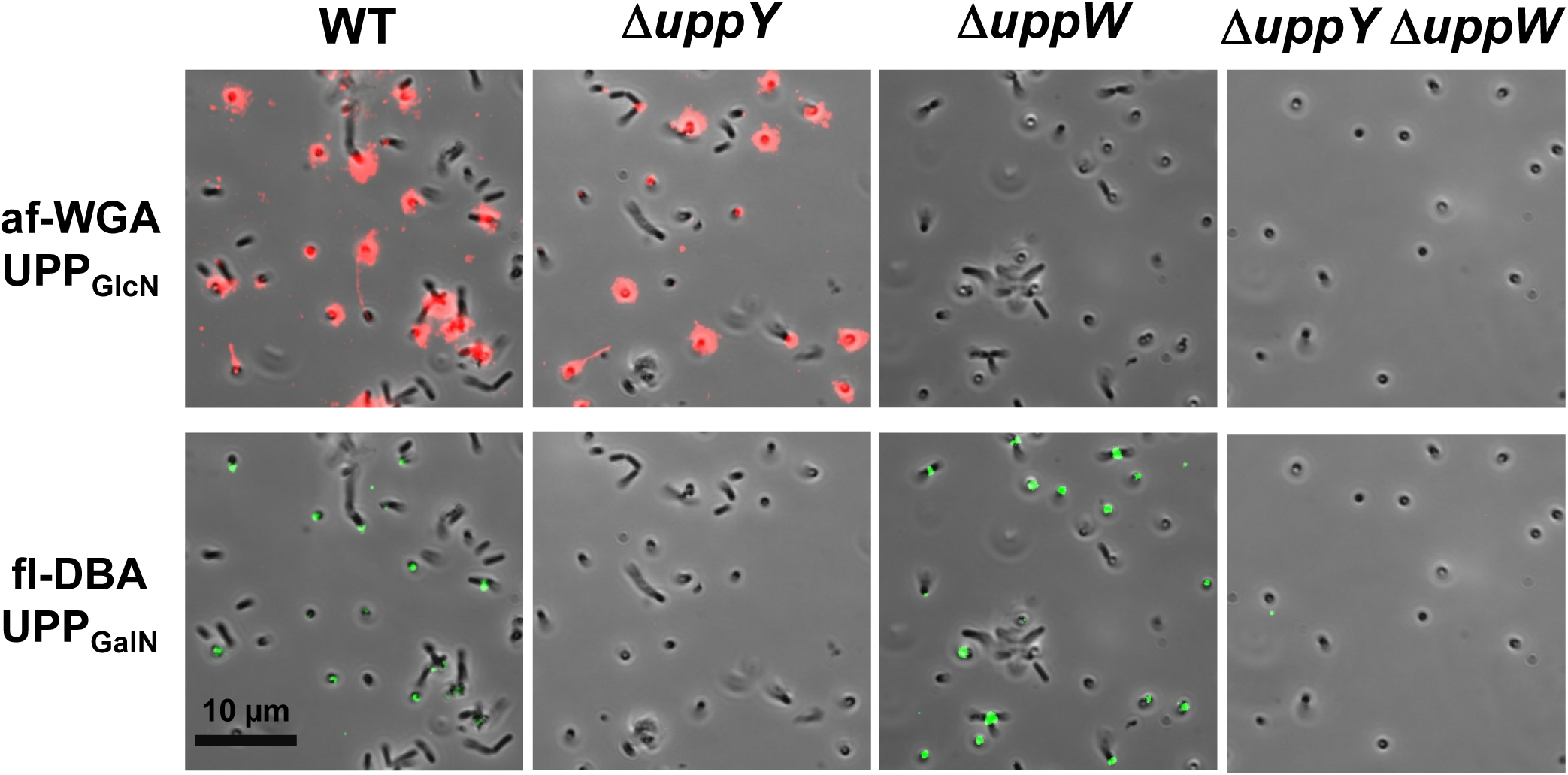
Dual lectin binding in *ΔuppW* and *ΔuppY* mutants shows reciprocal specificity. Short term attachment assay on a glass coverslip co-incubated with WT, Δ*uppY, ΔuppW,* and *ΔuppY ΔuppW* mutants co-incubated with af-WGA and fl-DBA showing reciprocal fluorescence of the polymerase mutants. Microscopy as in Fig. 5, but only merged images are shown.

UPP polysaccharide production was examined using transmission electron microscopy (TEM) and lectins conjugated with colloidal gold beads. The *A. tumefaciens* Δ*pruA* CDGS-strain was used for these experiments to eliminate additional polysaccharides that can obscure visualization of the UPP and because the surface-contact dependence of the wild type strain makes TEM of adhered cells challenging. The appropriate cultures were grown and probed with each lectin preparation individually, and then incubated with the general TEM reagent uranyl acetate. In the strain with functional copies of both *uppY* and *uppW*, the UPP could be observed as fibrous material at a single pole of the cell, that was absent in the Δ*uppY* Δ*uppW* mutant (Fig. 9). The gold-labeled WGA (20 nm) bound this fibrous material quite effectively and no binding was observed for the Δ*uppY* Δ*uppW* mutant. Fibrous polar material was visible for the single Δ*uppY* and Δ*uppW* mutants, but the labeled WGA only bound efficiently to the fibers observed in the Δ*uppY* mutant, and not those on the Δ*uppW* mutant (Fig. 9A). Initial experiments with the gold-labeled DBA lectin, revealed it to bind very inconsistently to the UPP (data not shown). We found that the lectin from peanut (PNA) that also binds to polysaccharides with GalNAc residues, gave more consistent results (Fig. 9B). The PNA lectin bound to the fibrous material for the cells that were wild type for both *uppY* and *uppW,* and the Δ*uppW* mutant, but no binding to the Δ*uppY* mutant or the double mutant. Although this pattern was observed consistently, it was clear that this lectin binds much less efficiently in this experiment to the UPP, than does WGA. The basis for this weaker binding remains to be determined.

**Figure 9.**
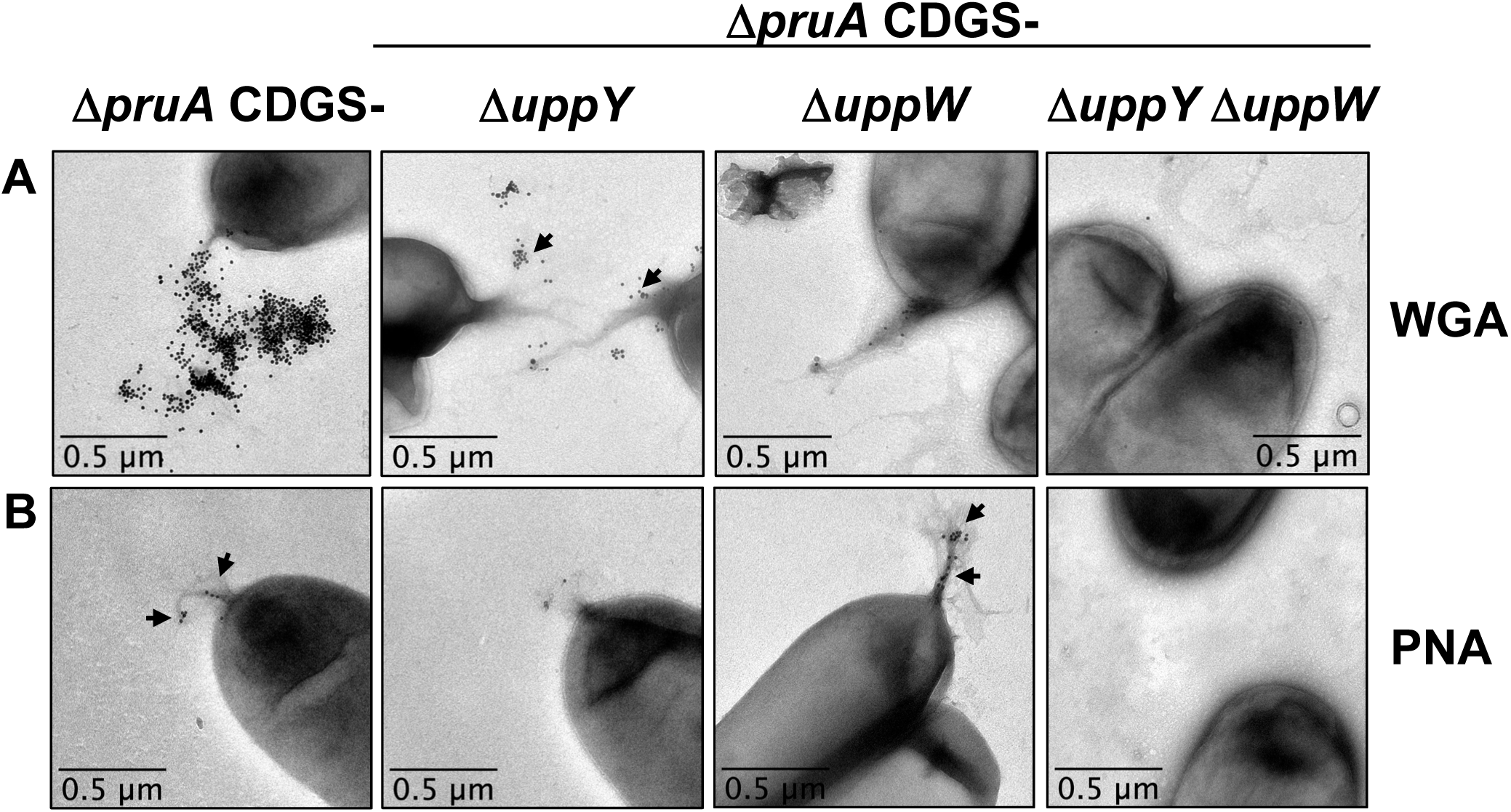
Transmission electron microscopy (TEM) of UPP from polymerase deletion mutants. TEM analysis of *A. tumefaciens* Δ*pruA* CDGS-mutant derivatives on a JEOL JEM-1010 Transmission Electron Microscope at 80 kV. (A) TEM using WGA conjugated with gold beads (20 nm). (A) TEM using PNA conjugated with gold beads (10 nm).

### UppH encodes a glycoside hydrolase capable of destabilizing the UPP

The *uppH* gene (ATU_RS1155) was found to be largely dispensable for adhesion and UPP production (Fig. S2A, Table S1). However, it was predicted to be a periplasmic Class 10 glycoside hydrolase, and its position within the *uppG-O* cluster implicated it in UPP production. Periplasmic glycoside hydrolases are often encoded within polysaccharide biosynthetic gene clusters, and are thought to hydrolyze glycosidic bonds within the associated polysaccharide as it transits this cellular compartment (Oglesby *et al*., 2008). Recently it was reported that exogenous addition of the purified PelAh and PslGh periplasmic glycoside hydrolases from *Pseudomonas aeruginosa* enzymes can disrupt Pel and Psl-dependent biofilms through cleavage of these polysaccharides (Baker *et al*., 2016). A derivative of UppH with its N-terminal secretion signal replaced by a hexahistidinyl tag (His_6_) was purified. His_6_-UppH was added in standard culture media (1 µM final concentration) to coverslips on which *A. tumefaciens* (*ΔpruA* CDGS-) had been allowed to adhere for 2 h, and incubated for an additional 1 h, followed by dual lectin labeling with af-WGA and fl-DBA. In the untreated samples, many cells were polarly attached and visibly labeled with one or both lectins (Fig. 10). Addition of the UppH enzyme to these attached populations did not dislocate the cells, but removed visible lectin-labeled polysaccharide. In parallel, we generated and purified a site-specific UppH mutant in which two predicted catalytic glutamate codons were mutated to glutamine codons to create UppH** (E152Q E266Q). Similar addition of exogenous, purified UppH** at 1 μM had no impact on lectin labeling (Fig. 10). We also examined UppH activity using TEM, and found that pre-incubation of *A. tumefaciens* Δ*pruA* CDGS-strain with the same concentration of purified UppH, effectively removed the unipolar fibrous material, resulting in poles with no visible fibers, whereas incubation with the UppH** mutant did not affect these fibers. Similar to studies using the *P. aeruginosa* glycoside hydrolases (Baker et al., 2016), addition of increasing concentrations of purified His_6_-UppH added at the beginning of cultivation inhibited *A. tumefaciens* biofilm formation (Fig. S5), whereas the catalytic site mutant had no effect (data not shown).

**Figure 10.**
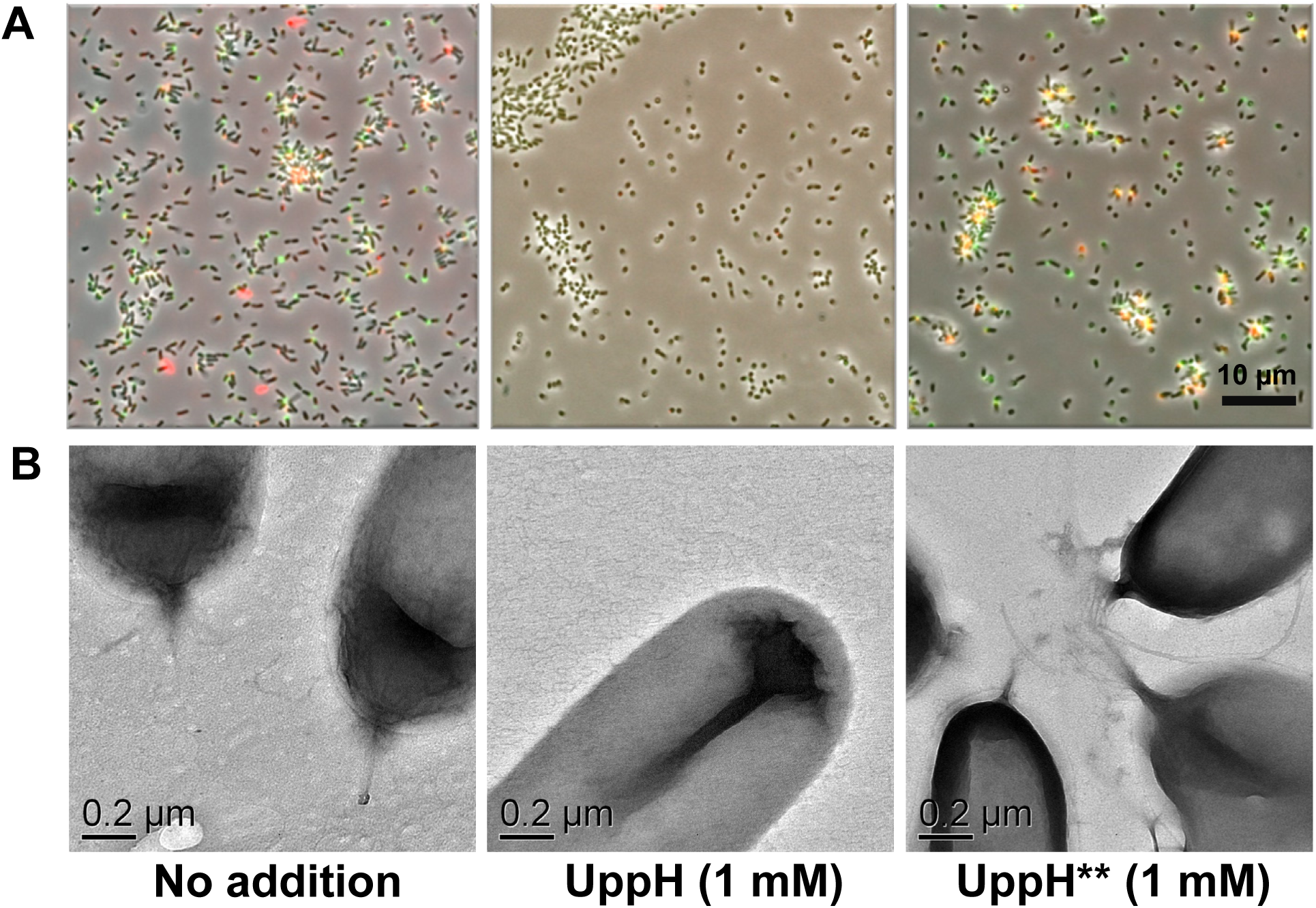
Purified UppH breaks down UPP in *A. tumefaciens.* Purified His_6_-UppH and His_6_-UppH** at 1 mM were incubated for 1 h with *A. tumefaciens* Δ*pruA* CDGS-. (A) Cells were pre-attached to glass coverslips prior to incubation with UppH preparations, and subsequently labeled with af-WGA and fl-DBA. Microscopy performed as in Fig. 5, with only the merged phase contrast and fluorescence images shown. (B) TEM performed on cells following incubation with UppH preparations as in Fig. 9, but without the addition of gold-labeled lectins.

### A second genetic screen to distinguish between UppY- and UppW-dependent UPP biosynthetic pathways

Our findings suggest that there are two distinguishable polysaccharides that comprise the UPP, UPP_GalN_ and UPP_GlcN_, and that each polysaccharide species requires a specific Wzy-type protein, UppY and UppW, respectively. The individual Δ*uppY* and Δ*uppW* mutants do not manifest pronounced deficiencies in CR staining nor in standard biofilm assays. However, there are a large number of other single mutants in presumptive UPP biosynthetic genes identified that do manifest these deficiencies. For example, deletion mutants in *uppC* (Wza, OPX, outer membrane secretin), *uppX* (Wzx, flippase), *uppE* (WbaP, PHPT initiating glycosyltransferase) and *uppQ (*GNAT acetyltransferase), all have very pronounced phenotypes. This suggests that certain UPP genes have functions that are required for production of both polysaccharides, whereas others, such as UppY and UppW, are specific to each polysaccharide. To identify UPP-related functions specific to each polysaccharide and those that play a role for both, a variation of the initial DCR was developed. Neither the individual *uppY* or the *uppW* mutant has a DCR phenotype in the Δ*pruA* CDGS-background. We reasoned that additional loss of the other UPP polysaccharide in these single mutants would cause a significant DCR colony phenotype as observed for the Δ*uppY ΔuppW* mutant (Fig. S6). Two strains were transposon mutagenized in parallel: the *A. tumefaciens* Δ*pruA* CDGS^-^ strain deleted for either the *uppY* gene or the *uppW* gene. We then screened these transposon mutant libraries for DCR mutants, examining roughly 30,000 colonies per lineage. We obtained and confirmed about 100 independent DCR mutants from each lineage. These mutants were pooled, total genomic DNA extracted and barcoded to specify each screening background, and then sequenced *en masse* using *Himar-1* transposon-specific amplification and Illumina sequencing ([dataset] Onyeziri and Fuqua, 2021). We expected to identify DCR mutations in the *uppY* gene for the Δ*uppW* derivative, and mutations in *uppW* in the Δ*uppY* derivative, and multiple mutants in these genes were isolated as predicted (Fig. 11, Table S3). Other previously identified genes from the initial DCR screen were also re-isolated. For example, multiple independent mutants in *uppE* (Atu1236) and *uppC* (Atu1238) were isolated in both mutant backgrounds. However, no transposon insertions in any of the other genes in the *uppABCDEF* cluster were isolated in either screen, despite the isolation of mutants in *uppB* and *uppD* in the prior screen (Fig. 11). The two flippase homologs, *uppX* and *uppV* were isolated in both screening backgrounds, although as observed before, considerably more mutants were isolated in *uppX*. Overall, multiple transposon mutations were re-isolated in both the Δ*uppY* and Δ*uppW* screening backgrounds in 12 different *upp* genes that had been identified in the prior screen, (Fig. 11, paired green and red bars; Table S3). Several other genes were identified in both screens that had not been previously associated with UPP biosynthesis, notably a *galE*/*exoB* homolog (ATU_RS19470), a *traA* plasmid conjugation gene homolog (ATU_RS22800, located on the linear chromosome), several predicted regulators (see below), and two genes of unknown function (ATU_RS15030 and ATU_RS22740).

**Figure 11.**
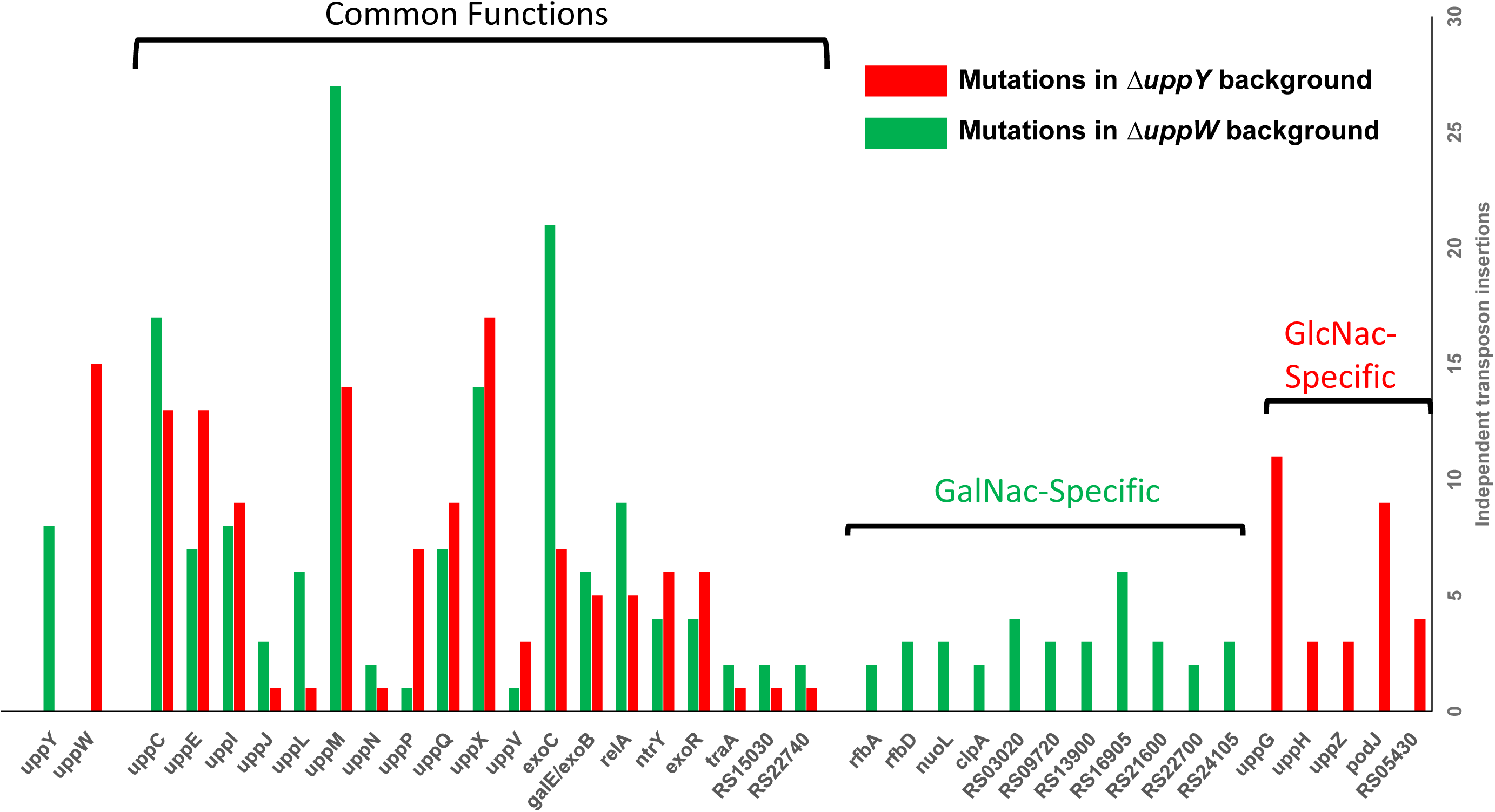
Genes isolated in UPP pathway-specific screens. On the y-axis are number of independent transposon insertions that resulted in a scorable DCR phenotype for the Δ*pruA* (CDGS-) polymerase deletion mutants (Δ*uppY* and Δ*uppW*), and on the x-axis are the genes identified with transposon insertions. Red bars indicate genes disrupted in the *ΔuppY* background and therefore predicted to be important for UPP_GlcN_ synthesis. Green bars indicate genes identified in the *ΔuppW* background and thus predicted to be important in the synthesis of UPP_GalN_. Red and green bars together indicate genes identified in both mutant screening backgrounds.

### Genes identified in only one of the polymerase mutants

The only *upp* genes other than *uppY* and *uppW* that had been identified before, which were isolated in only one of the pathway-specific screens, were *uppG* and *uppH.* They were identified multiple times in the Δ*uppY* mutant but not once in the Δ*uppW* mutant (Fig. 11, single red bars, Table S3). As described, UppG encodes a glycosyltransferase in the GT4 family, whereas *uppH* is a Class 10 glycoside hydrolase. Given the polarity of transposon insertions in *uppH on uppG* that we had already confirmed, it is very likely that the DCR phenotype in the *uppH* transposon mutants was due to their effect on *uppG* expression. Thus, *uppG* is probably the only established UPP biosynthesis function in addition to the *uppW* polymerase that is required specifically for UPP_GlcN_ production and does not appear to play a prominent role in UPP_GalN_ production (at least in the Δ*uppY* background). However, another polysaccharide related gene we have now designated *uppZ* (ATU_RS11500) encoded convergently from the *uppW* Wzy-type gene, was disrupted in three independent sequences. UppZ is an OafA homolog similar to ExoZ, required for production of acetylated succinoglycan (Reuber & Walker, 1993). Two additional genes were identified only in the *ΔuppY* screen. Four mutants were isolated in ATU_RS05430, predicted to be a fatty acid biosynthesis gene. Nine independent mutations were isolated in the *podJ* gene (ATU_RS02460), a polarity determination factor known to be required for normal polar localization of the *C. crescentus* holdfast (Hinz *et al*., 2003) and required for functions related to polar cell division in *A. tumefaciens* (Anderson-Furgeson *et al*., 2016). No *podJ* mutants were identified in the Δ*uppW* background.

Transposon mutants in 11 different genes were isolated specifically in the Δ*uppW* mutant that resulted in loss of the CR phenotype imparted by the UPP_GalN_ polysaccharide (Fig. 11, single green bars, Table S3). Although these genes were independently disrupted in multiple DCR transposon mutants, the majority were isolated in fewer than 5 mutants. Two genes with transposon insertions from the Δ*uppW*-specific screen are adjacent, and annotated as *rfbA* (ATU_RS21635) and *rfbD* (ATU_RS21640), encoding a glucose-1-phosphate thymidylyltransferase and a dTDP-4-dehydrorhamnose reductase. They are located in an operon of genes predicted to be involved in synthesis of the dTDP-L-rhamnose, often a component of O-antigen synthesis for LPS in members of the *Rhizobiaciae* (De Castro *et al*., 2008), and thus they may be involved in polysaccharide synthesis. The *nuoL* gene (ATU_RS06310) and *clpA* (ATU_RS06730) encode a NADH ubiquinone oxidoreductase subunit, and the ATP-binding subunit of the Clp protease. Seven of the genes identified are hypothetical reading frames.

### Genes involved in sugar-nucleotide precursor synthesis

Multiple independent mutants were isolated in the *exoC* gene (ATU_RS19035), in both screening backgrounds (Fig. 11, Table S3), consistent with the prior screen (Fig. 2F). As described earlier, *exoC* encodes a phosphoglucomutase, predicted to convert glucose-6-phosphate to glucose-1-phosphate, a precursor of sugar nucleotide substrates for polysaccharide biosynthesis (Kamoun *et al*., 1989). However, a gene annotated as *galE/exoB* (ATU_RS19470) was not isolated in the original screen, but was isolated with 5 or more independent transposon mutants in both pathway-specific screens. This gene is predicted to encode a UDP-glucose epimerase, similar to GalE, that catalyzes the reversible epimerization reaction of the sugar nucleotide precursor UDP-GlcNAc to UDP-GalNAc, and has an NADP Rossman domain (Holden *et al*., 2003). Given that we distinguish the two polysaccharides by the presence of GlcNAc and GalNAc, the role for this enzyme may be particularly relevant to the separation between pathways.

### Additional regulators identified in the pathway-specific screen

Several probable regulatory genes were identified in the UPP_GlcN_ and UPP_GalN_ specific screen (Fig. 11, Table S3). Mutations in three of these were identified in both screens. The *exoR* gene (ATU_RS08400), encodes a periplasmic regulator of the ChvG-ChvI two-component system (Heckel *et al*., 2014). In *A. tumefaciens*, *exoR* mutations dysregulate hundreds of genes, including those for synthesis of the polysaccharide succinoglycan. Mutations in *exoR* were also isolated as DCR mutants in the original screen (Xu and Fuqua, unpublished results). The two other putative regulators identified had not been implicated from the earlier screen. In both the Δ*uppY* and Δ*uppW* backgrounds, mutations were isolated for a homologue of the *relA*/*spoT* gene (ATU_RS05105) predicted to direct synthesis and degradation of the (p)ppGpp stringent response signal in *A. tumefaciens* (Zhang *et al*., 2004). The *relA/spoT* gene is predicted to be monocistronic. Likewise, multiple mutations in both backgrounds were also isolated in a gene annotated as *ntrY* (ATU_RS07130), encoding a predicted two-component sensor kinase homologue, located immediately upstream of a two-component response regulator annotated as *ntrX* (ATU_RS07135).

### Conservation of UPP biosynthesis genes

We used translated sequences of the UPP genes identified in this study as queries to search for homologs among the large number of genome sequences of members of the *Rhizobiaceae*. It is recognized that rhizobia and agrobacteria are polyphyletic and intermixed (Slater *et al*., 2009). Members of the *Agrobacterium* genus were historically classified into three primary biovars (I, II, and III), based on multiple attributes, and many of the wide host range *A. tumefaciens* strains including C58 are in Biovar I (Morton & Fuqua, 2012b). Recent phylogenetic studies have also divided the *A. tumefaciens* group itself into over 20 separate genomospecies (Lassalle *et al*., 2011). *A. tumefaciens* C58, the type strain we use in this study, is in Genomospecies 8 (G8, also given the species designation *A. fabrum*). There is now a wealth of genomic data for comparison across the agrobacteria and among the wider *Rhizobiaceae* that allows great insights into the genetic conservation and ecological context among these bacteria (Weisberg *et al*., 2020).

A comparison of the UPP genes identified in the current study across representative genome sequences reveals multiple interesting trends. Several of the genes are widely conserved in sequence and synteny (Fig. S7, S8). First, the *uppA-F* gene cluster is conserved largely throughout the agrobacterial genomospecies, with UppA less well conserved but still present (Fig. S7). This gene cluster is fully or partially conserved across the *Rhizobiaceae*, including among bacteria such as *Rhizobium leguminosarum* and *Rhodopseudomonas palustris* that are both recognized to make a UPP-type structure (Fritts *et al*., 2017, Williams *et al*., 2008). Beyond the *Rhizobiaceae* the gene cluster is also partially conserved more widely in *Brucella abortus* and other mammalian pathogens in the Alphaproteobacteria group. The *uppA-F* cluster shares some similarity to the <u>h</u>oldfast <u>s</u>ynthesis (*hfs*) gene cluster from *C. crescentus*, but is clearly divergent between bacteria in the Orders *Rhizobiales* and the *Caulobacterales* (Fritts *et al*., 2017).

Other UPP genes that are well conserved across the agrobacteria genomospecies are UppV and UppQ, the flippase and acetyltransferase encoded within a presumptive operon, and the UppP outer membrane protein. The sugar nucleotide enzyme ExoC and a GalU homolog (ATU_RS17570) encoded convergently with *uppP* (but not identified in our genetic screens) are also well conserved across the agrobacteria and the rhizobia. Strikingly, the *uppG-O* and *uppXY* gene clusters, and *uppW* gene are well conserved within the genomospecies considered Biovar I (G1, G2, G3, G4, G5, G7, G7-2, G8, G9, G13), with a few exceptions for specific genes within the clusters, but are largely absent outside of these taxa.

## Discussion

In this study, we identified and analyzed a large number of genes required for biosynthesis of the *A. tumefaciens* unipolar polysaccharide known as the UPP, and in so doing, have discovered that this structure is composed of two genetically and biochemically discrete polysaccharides.

### Genetic screens provisionally identify UPP biosynthetic genes

One of the two separable biosynthetic pathways is UppW-dependent, and drives synthesis of the UPP species that labels with the WGA lectin specific for GlcNAc residues (UPP_GlcN_). The other pathway is UppY-dependent and directs production of the UPP species that labels with the DBA and PNA lectins specific for GalNAc residues (UPP_GalN_). Our genetic approach defined a large number of genes implicated in UPP synthesis, identifying multiple components of a presumptive Wzx/Wzy pathway including formation of sugar nucleotide precursors (Fig. 12), several regulators and a collection of genes of unknown function. These screens were remarkably specific for UPP deficiencies, and creation of in-frame deletions in many of the genes identified from the screen revealed clear defects in UPP-dependent phenotypes in wild type and re-constructed screening backgrounds. For our first DCR screen in which the *uppY* and *uppW* genes were intact, the majority of genes identified were predicted polysaccharide-related functions (Fig. 2 and Table S1), with the exception of *uppP*, which has the sequence of an outer membrane protein with no predicted role in polysaccharide biosynthesis, and two genes of unknown function, *uppK* and *uppN*. Only three of the disrupted genes identified in DCR mutants within the genetic clusters predicted to encode Wzx-Wzy-type pathway components, failed to impact UPP-dependent phenotypes when the corresponding in-frame deletions were generated (Δ*uppK*, Δ*uppO*, Δ*uppH*) in otherwise wild type *A. tumefaciens*. Transposon insertions in *uppH* were polar on the downstream glycosyltransferase *uppG*, although the evidence suggests the UppH glycoside hydrolase recognizes the UPP structure. The incongruity between the *uppK* and *uppO* transposon mutants and in-frame deletion mutants in these genes is however not explained by simple polarity, and the basis for these differences requires further investigation.

**Figure 12.**
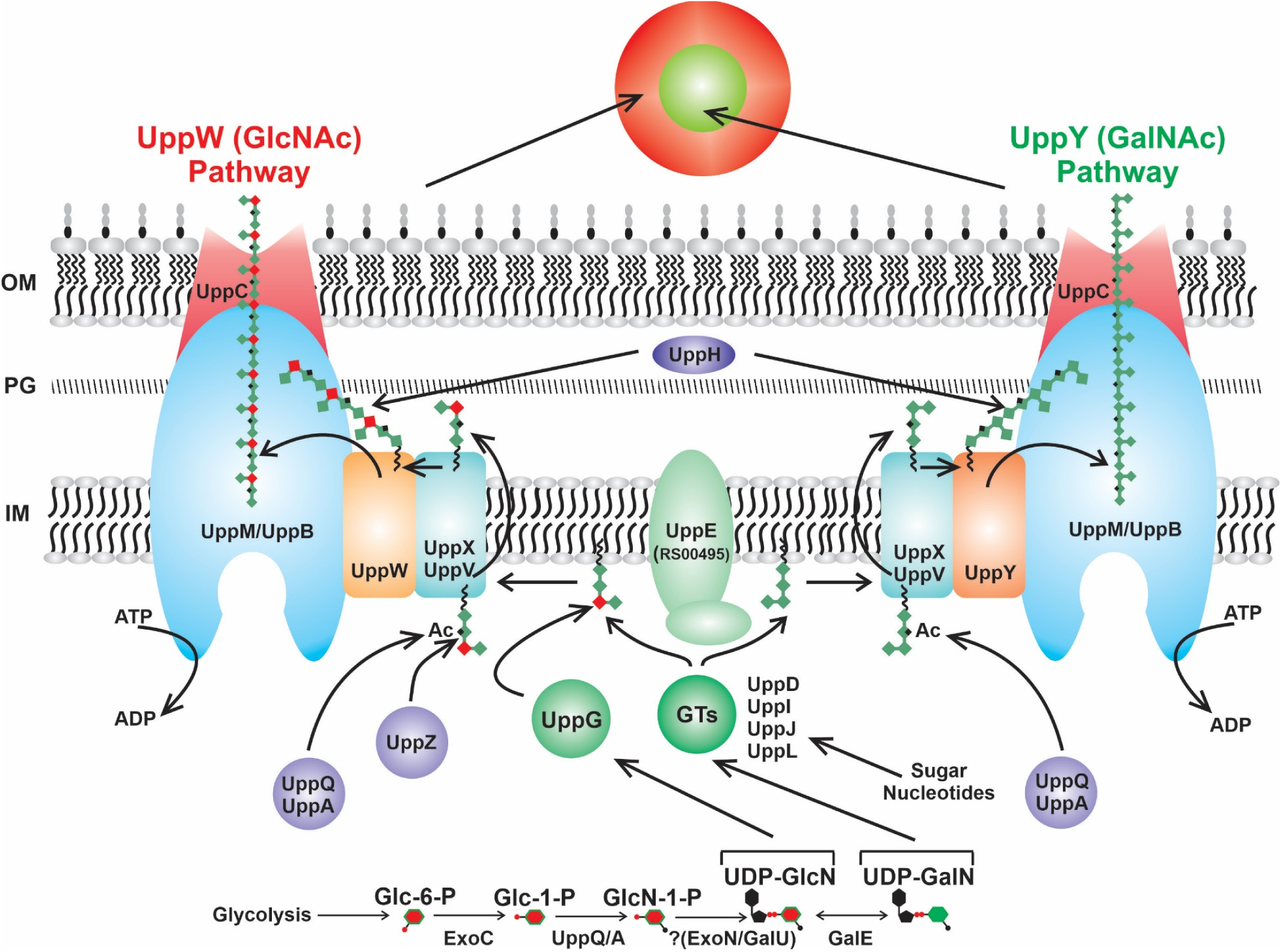
Model of UPP biosynthesis pathway in *A. tumefaciens.* Dual Wzx/Wzy polysaccharide biosynthesis pathway showing proteins that exhibit specificity for UPP_GlcN_ or UPP_GalN_, and proteins involved in synthesizing both polysaccharides. Color-coding is consistent with the legend in Fig. 2. IM – inner membrane; PG-Peptidoglycan; OM – outer membrane; GT – glycosyl transferase; Ac – acetylation.

### Initial steps in UPP precursor synthesis

The genes we have identified represent all of the requisite components of a complete Wzx/Wzy-type polysaccharide biosynthesis pathway (Whitfield *et al*., 2020), including homologs to flippases (Wzx-type proteins, UppX and UppV) and polymerases (Wzy-type proteins, UppY and UppW) (Fig. 12). Our preliminary model is that UppE functions as the WbaP initiating glycosyltransferase for UPP biosynthesis, or PHPT (polyisoprenylphosphate hexose-1-phosphate transferase), adding the first monosaccharide to an undecaprenol pyrophosphate (Und-PP) lipid carrier. Our prior studies found that under phosphorus (P_i_) limitation, the requirement for *uppE* could be supplanted by another PHPT-type protein Atu0102 (Xu *et al*., 2012), but under P_i_-replete conditions as used in the current study, the role for UppE is clear. Interestingly, multiple PHPT homologs are redundantly functional in holdfast biogenesis in *C. crescentus* (Toh *et al*., 2008).

As with other polysaccharide biosynthetic pathways, precursors for this and subsequent monosaccharide additions are supplied by cytoplasmic sugar nucleotide pathways that in *A. tumefaciens* and its relatives, depend upon ExoC to convert glucose-6-phosphate (Glc-6-P) to glucose-1-phosphate (Glc-1-P) (Kamoun *et al*., 1989). As GlcNAc appears to be a major component of the UPP, we predict Glc-1-P is acetylated to form *N*-acetyl glucosamine-1-phosphate (GlcN-1-P) by one of the putative acetyl transferases, UppA or UppQ (Fig. 12), mutations in either of which cause severe UPP defects. The GlcNAc would be conjugated to UDP, to form UDP-GlcNAc, although it is possible that UDP linkage occurs prior to acetylation. Our genetic screens did not identify any UTP-glucose-1-phosphate uridylyltransferases as DCR mutants, but the genes annotated *exoN* (ATU_RS18920) and *galU* (ATU_RS16570), which is encoded adjacent to *uppP* are both predicted to catalyze this reaction and are non-essential (Curtis & Brun, 2014), so it is possible they are functionally redundant for this activity. The GalE homolog (ATU_RS19470) identified with a general UPP defect in our pathway-specific screen (Fig. 11, 12) is predicted to reversibly epimerize UDP-GlcNAc to UDP-GalNAc, an interesting activity since these sugar moieties are likely to be defining residues for the UPP_GlcN_ and UPP_GalN_ polysaccharides, respectively. Furthermore, sugar-nucleotide precursors for UPP synthesis are likely to be generated through these basal sugar-nucleotide pathways, although no additional genes annotated to function in this capacity were identified.

Following the Wzx-Wzy general model, after addition of the initiating monosaccharide to the Und-PP carrier to form monoglycosylated Und-PP, other glycosyltransferases utilize sugar-nucleotide precursors to add monosaccharides to form the mature subunit (Whitfield et al., 2020). Generally, the first of these monosaccharide additions is mediated by a glycosyl transferase homologous to WecG/TagA, generating the diglycosylated lipid II intermediate, which can then be acted upon by additional glycosyl transferases (Ginsberg *et al*., 2006) Five predicted glycosyltransferases were identified in one or both of our genetic screens (*uppD*, *uppG*, *uppI*, *uppJ* and *uppL*). UppL is predicted to encode a glycosyltransferase (CAZy Group GT26) homologous to WecG and TagA, and is strictly required for UPP synthesis. It is also the best match to the HfsJ glycosyl transferase of *C. crescentus*, a WecG/TagA type protein required for holdfast synthesis and a point in the pathway at which holdfast production is regulated (Fiebig *et al*., 2014). We hypothesize that UppL generates the Lipid II diglycosyl intermediate for UPP biosynthesis. The four additional GTs can then act to assemble the UPP repeating subunit (Fig. 12).

The *uppI, uppJ, and uppL* genes clearly all play roles in UPP synthesis, as they were consistently identified in our DCR mutant screens and in-frame deletions have significant impact on UPP production and attachment. The *uppD* gene was identified as a DCR mutant in our initial screen with *uppY* and *uppW* intact (in all four ECR screening backgrounds) and the Δ*uppD* mutant is clearly biofilm deficient and greatly diminished for UPP production. However, it was not identified in the pathway-specific screen, suggesting that the requirement for UppD may be less stringent. Given our evidence for a UPP_GlcN_-specific role for UppG, we propose that it adds a particular residue that acts to divert a portion of the growing subunits to the UppW-dependent pathway, whereas UPP subunits in which this additional residue is not incorporated proceed through the UppY-dependent pathway. Additional modifications to the UPP subunits may include more acetylation given the three potential acetyl transferases we have identified, UppA, UppQ, and UppZ.

### Translocation, polymerization and export of the UPP

A defining step in Wzx-Wzy pathway is translocation of the mature repeating subunit linked to Und-PP from the cytoplasmic face of the inner membrane to its periplasmic face (Islam and Lam, 2014). Mutants in *uppX*, encoding a predicted flippase with 10-12 transmembrane domains, have a severe UPP deficiency and were identified repeatedly in our screens. In addition, the *uppV* gene also encodes a flippase homolog, that was consistently identified (albeit less frequently than *uppX*, see Figs. 2 and 11 and Tables S2 and S3) and also manifests a pronounced UPP defect. Our initial prediction that each flippase would be specific for one of the two polymerases did not hold true, as each flippase manifests defects for both UppW- and UppY-dependent polysaccharides. We therefore tentatively assign both UppX and UppV as flippases in our model, with perhaps UppX playing the primary role (Fig. 12).

Once in the periplasm, the UPP subunits initiate or add to the growing polysaccharide chain by the activity of either UppW or UppY, which are strictly required for UPP_GlcN_ and UPP_GalN_, respectively (Fig. 12). We predict that the specific monosaccharide composition of the subunit dictates which polymerase drives chain elongation. Addition of the subunit to the growing chain associated with its cognate polymerase, releases the Und-PP which is then available to serve for the multiple other pathways that depend on it, such as for synthesis of peptidoglycan and other polysaccharides.

Based on the Wzx/Wzy paradigm, the UPP polysaccharide then associates with a periplasmic co-polymerase (PCP, Wzc) and is ushered to the outer membrane polysaccharide secretin (OPX, Wza) for export to the cell exterior (Whitfield *et al*., 2020). The details of this process and the specific role for the PCP remain ill-defined for any biosynthetic pathway, but it is clear that the PCP and OPX proteins form a large multimeric complex that spans the outer membrane, and the periplasm, projecting into the cytoplasm (Collins *et al*., 2007). Our genetic analysis suggests that the predicted PCP-2a proteins, UppM and UppB, fulfill the PCP (Wzc) requirement for both UPP polysaccharides and that UppC functions as the OPX (Wza) secretin (Fig. 12). Other important processing may occur in the periplasm, such as the activity of the periplasmic glycoside hydrolase UppH (although this is not required for UPP production) and perhaps linkage of the UPP polysaccharide to LPS via the O-antigen ligase UppF (also see below). What retains the externalized UPP_GlcN_ and UPP_GalN_ at discrete locations at a single pole remains to be determined. Our preliminary results suggest that several of the UPP biosynthetic proteins identified here localize to the same pole as the UPP (Natarajan and Fuqua, unpublished) suggesting that at least some of the steps of biosynthesis occur proximal to the eventual site of UPP elaboration, similar to what has been observed for holdfast biogenesis in *C. crescentus* (Javens *et al*., 2013). A role for the PodJ polar localization factor in proper localization of the UPP to the pole is supported by its identification in our genetic screens (Fig. 11).

The pathway specific screen identified a significant number of genes in which multiple independent transposon insertions resulted in the DCR phenotype indicative of UPP deficiencies, that had not been identified in the initial screen with *uppY* and *uppW* intact. Some of these genes are annotated to be related to polysaccharide biosynthesis (*rfbB*, *rfbD*, and *exoZ*), predicted regulators (*relA*, *ntrY*) or the polarity determinant *podJ.* It is unclear what role the other genes with annotations either that predict functions outside of polysaccharide synthesis or hypothetical genes, might play in UPP synthesis. Although the isolation of multiple independent mutants supports their impact on UPP synthesis, more extensive genetic analysis is required to conform their roles.

### Anchoring and cohesiveness of the UPP

Interestingly, individual in-frame deletions of all of the *uppA-F* genes result in UPP deficiencies although no mutations were identified for *uppA* or *uppF* in any of the DCR genetic screens performed. UppA encodes a cytoplasmic GNAT-type acetyl transferase.

For several bacterial polysaccharides, the degree and distribution of acetylation is thought to alter the adhesiveness of the polysaccharide (Vuong *et al*., 2004, Wan *et al*., 2013). UppA may modulate adhesiveness of the UPP, but its loss may not visibly impact the CR-staining properties of the UPP. There is a strict requirement for the other cytoplasmic GNAT-type protein UppQ for the ECR phenotype, and so we speculate that UppA performs a non-critical role such as UPP modification, but we do not know with certainty, so both UppA and UppQ are depicted to modify the UPP (Fig. 12). The *uppZ* gene identified in the pathway-specific screen performed with the Δ*uppY* mutant, also encodes a predicted acetylase, but this was only identified in the Δ*uppY* mutant suggesting specificity for the UPP_GlcN_ polysaccharide.

UppF is of particular interest, as it encodes a homolog of O-antigen ligases that function in attaching the O-antigen portion of LPS to the LPS core sugars during biogenesis of this outer membrane component (Ruan *et al*., 2018). The dramatic biofilm and attachment deficiencies for the Δ*uppF* mutant suggest a profound UPP defect, but it was not identified in either of the DCR screens. In *C. crescentus,* the holdfast is attached to the tip of the stalk via the <u>h</u>oldfast <u>a</u>nchor (Hfa) proteins, that are thought to form a β-amyloid-type fiber (Hardy *et al*., 2010). However, there are no clear homologs of the Hfa proteins encoded by *A. tumefaciens* and we speculate that instead the UPP may be linked to LPS on the cell surface. Given its combination of phenotypes, UppF is a promising candidate for linking the UPP to LPS.

### The polysaccharide component of the *A. tumefaciens* UPP

As with the *C. crescentus* holdfast, the UPP structure was first detected using lectin reagents that are generally specific for polysaccharides (Tomlinson & Fuqua, 2009). Lectin labeling remains the most facile and reliable way to identify the *A. tumefaciens* UPP microscopically, although it is also visible in TEM and SEM without specific labeling (Xu *et al*., 2013). As discussed above, our genetic analyses have identified many genes that are required for UPP biosynthesis, and the majority of these can be linked to polysaccharide-related functions, specifically for Wzx-Wzy-type pathways. We have also identified several regulators and the polar localization factor PodJ as important for UPP biosynthesis. Our pathway-specific screen identified a significant number of genes of unknown function, which may or may not be related to polysaccharides, and remain to be investigated further. The UppP protein, predicted to encode an outer membrane porin (distinct from polysaccharide secretins such as UppC), is not a typical component of a Wzx/Wzy pathway, but is required for UPP production. Despite these few exceptions, the prominent role for Wzx/Wzy proteins and other polysaccharide related proteins (e.g. sugar nucleotide precursors) strongly support the conclusion that the UPP is predominantly comprised of polysaccharide.

We have added to this evidence by finding that the extracellular addition of the purified UppH glycoside hydrolase degrades all visible lectin-binding UPP material on pre-attached cells, and can inhibit biofilm formation when co-incubated with planktonic *A. tumefaciens*. The UppH protein is annotated in the *A. tumefaciens* C58 genome as a xylosidase (Goodner *et al*., 2001, Wood *et al*., 2001), and a recent study, tentatively designating the gene *xynA*, found that it was required for growth on minimal medium with exogenous xylan as a carbon source (Mathews *et al*., 2019). Our observations on susceptibility of both UPP_GlcN_ and UPP_GalN_ to UppH (XynA) activity suggest that the enzyme hydrolyzes important glycosidic linkages common to both UPP polysaccharides. Based on its activity on exogenous xylan and its annotation, we hypothesize that these vulnerable positions within the UPP contain xylose residues. A recent chemical characterization of the *C. crescentus* holdfast revealed that it contains xylose residues, among other monosaccharides (Hershey *et al*., 2019). The Δ*uppH* mutant does not have a detectable attachment phenotype, but this is a relatively common observation among periplasmic glycoside hydrolases associated with polysaccharide synthesis pathways (Baker *et al*., 2016). The roles for these enzymes in polysaccharide biosynthesis remain ill-defined despite their utility as purified reagents with high specificity for their cognate polysaccharides, and their anti-biofilm potential. We propose that UppH is a component of the UPP biosynthetic machinery that is active on the polysaccharide component of this structure. Interestingly, although exogenous UppH incubation clearly removes the UPP as visualized with lectin labeling and TEM, pre-attached cells remain on surfaces after treatment, and there is incomplete biofilm inhibition in the presence of UppH. This may reflect a curing process of the UPP, as has been observed for the *C. crescentus* holdfast (Hernando-Perez *et al*., 2018) to a glycosidase resistant, sterically blocked form, or alternatively the function of an as yet unidentified component that is resistant to the UppH enzyme.

Given the number of genes encoding Wzx-Wzy components that are required for both UPP_GlcN_ and UPP_GalN_ synthesis, and their mutual susceptibility to the UppH glycoside hydrolase, we hypothesize that they have closely related structures and perhaps a common core motif. Two related UPP polysaccharides would be similar to the *Vibrio cholerae* VPS polysaccharide which occurs as two distinct species, one which contains α-d-Glucose among other monosaccharides, and another less abundant species where this residue is α-d-GlcNAc (Yildiz *et al*., 2014). The lectin specificity of the two UPP species suggests that one species contains GlcNAc and the other GalNAc, but this remains to be proven by chemical characterization of the two different polysaccharides. Interestingly, the only annotated biosynthetic genes other than the polymerase genes *uppW* and *uppY* that are specific for each polysaccharide, are *uppG,* encoding a glycosyltransferase *uppZ*, an *exoZ* homolog, both involved in UPP_GlcN_ synthesis but not UPP_GalN_, and potentially the *rfbA* and *rfbD* homologs specific for UPP_GalN_ and not UPP_GlcN_. The observation that the Δ*uppG* mutant synthesizes a very small UPP, but still detectable with WGA, suggests that it is not solely responsible for addition of GlcNAc residues. We speculate that UppG may either contribute to addition of GlcNAc residues, or other monosaccharide residues that then dictate the efficiency of polymerization by UppW, and eventual elaboration of UPP_GlcN_. Wzy-type polymerases can be highly specific, and subtle differences in the linkages and monosaccharides can dictate recognition between particular Wzy-type proteins (Hong *et al*., 2015). The role of the *uppZ* in UPP synthesis remains to be investigated more fully.

### Physical separation of UPP_GlcN_ and UPP_GalN_ at cell poles

The sectored, bullseye pattern observed for lectin labeling when cells producing the UPP are incubated with both the DBA and WGA lectins suggests that the UPP_GlcN_ and UPP_GalN_ polysaccharide species are localized differently relative to each other (Fig. 6). The DBA labeling is usually proximal to the tip of the cell, whereas the WGA labeling, while still unipolar, is more diffuse and visibly surrounding the DBA-labeled material. This pattern suggests that these polysaccharides may be localized to specific positions around the pole. Alternatively, they may be extruded from the same point on the pole, but are temporally synthesized in stages. How the roles for these multiple polysaccharide species are integrated during the surface attachment process, and how their biophysical properties may differ are current topics of investigation.

### Mixed conservation among the UPP biosynthesis genes

The six gene *uppABCDEF* cluster we initially identified is fully or partially conserved across members of the *Rhizobiales*. Among the expanded UPP biosynthesis genes we have identified here, some show similar wide distribution, such as the *uppVQ* genes. However, several genes are restricted to the Biovar I group including the G8 genomospecies of *A. tumefaciens* C58. This is intriguing as many of the genes in these clusters encode functions that are required for both UPP_GlcN_ and UPP_GalN_ such as the *uppX* flippase and the *uppL* glycosyltransferase. Conversely, the *uppY* and *uppW* genes encode proteins that have specificity for UPP_GalN_ and UPP_GlcN_, respectively. *R. palustris* produces a UPP-type structure that labels with WGA and presumably contains GlcNAc residues, but only the *uppA-F* cluster is conserved (with the *uppC* secretin in a different position), and a flippase has yet to be identified (Fritts *et al*., 2017). In *C. crescentus*, the holdfast was initially defined by WGA binding and is now know to contain GlcNAc, xylose, and other monosaccharides (Hershey *et al*., 2019), and *A. tumefaciens* homologs to the *hfs* genes reside in the *uppA-F* cluster, as well as others outside, such as the glycosyl transferases *uppJ* and *uppL*, which are similar to *hfsL* and *hfsJ*, respectively. In *R. leguminosarum,* the *uppA-F* cluster (designated as *gms* genes for glucomannan synthase) is conserved and syntenous with that from *A. tumefaciens*, but the glucomannan unipolar polysaccharide it directs production of is composed of glucose and mannose and is recognized by host pea plants via a specific lectin (Laus *et al*., 2006, Williams *et al*., 2008). Clearly, diversification of these genes has led to production of different polysaccharides, with niche-specific functions. Over evolutionary history, several *A. tumefaciens* lineages within the Biovar I type strains have apparently bifurcated their UPP into two different chemical species, UPP_GlcN_ and UPP_GalN_, by the activity of an expanded and intertwined gene set that drives their synthesis. The UPP genes in general provide a fascinating example for diversification of cell surface polysaccharides, features that are arguably very much exposed to environmental selective pressures. The specific roles for the expanded UPP genes in the ecological context of the *A. tumefaciens* Biovar I lineages, including but not restricted to plant interactions, is an area of future interest.

## Experimental Procedures

### Strains, plasmids, and reagents

The bacterial strains and plasmids used in this study are listed in Table S4. Oligonucleotide sequences are available upon request. *A. tumefaciens* strains were grown at 28°C or 30°C on liquid or solid LB (10 g of Bacto-tryptone, 5 g of yeast extract and 5 g of NaCl per liter, pH 7.0) or AT minimal medium (Tempé *et al*., 1977) with 0.5% (w/v) glucose and 15 mM ammonium sulfate to make ATGN. For biofilm assays, 22 µM FeSO_4_ was added to the liquid ATGN medium immediately before inoculation, and for *sacB* counter-selection, 2.5-5% (w/v) sucrose replaced glucose as the sole carbon source to make ATSN. For ATGN-CR, the CR dye was dissolved in Milli-Q water (Millipore Sigma, Burlington, MA) to a concentration of 20 mg/ml, then passed through a 0.2 µm filter to remove aggregates. 250 µl of this stock solution was added per 100 ml ATGN solid medium to make a final concentration of 50 µg/ml ATGN-CR. *E. coli* strains were grown at 37°C in LB medium. Motility assays were performed in 0.3% ATGN agar in 60 mm plates, inoculated in the center of the medium with the tested strain, and incubated at 30° C. Plasmids were introduced into chemically competent cell preparations of *E. coli*, and into *A. tumefaciens* via conjugation or electroporation (Morton & Fuqua, 2012a). Oligonucleotide primers were obtained from Integrated DNA Technologies, Coralville, IA, and single-primer extension DNA sequencing was performed by ACGT, Inc., Wheeling, IL. Chemicals, antibiotics, and culture media were obtained from Fisher Scientific and Sigma-Aldrich. When necessary, antibiotics were added to the medium as follows: 100 µg/ml ampicillin (Ap), 50 µg/ml gentamycin (Gm) and 50 µg/ml kanamycin (Km) for *E. coli*, and 300 µg/ml Gm and Km for *A. tumefaciens*. 250 µM Isopropyl-β-D-thiogalactopyranoside (IPTG) was used to induce expression of plasmid borne genes unless otherwise stated. All fluorescently conjugated lectins were obtained from Invitrogen; FITC-labeled WGA (fl-WGA), Alexafluor 594-labeled WGA (af-WGA), FITC-labeled DBA (fl-DBA),.

### Controlled expression plasmids

Wild type coding sequences of genes to be expressed were amplified from *A. tumefaciens* C58 genomic DNA (unless otherwise stated), and cloned into the LacI^Q^ encoding, IPTG inducible plasmid pSRK carrying either a gentamycin (pSRKGm) or kanamycin (pSRKKm) resistance cassette (Khan *et al*., 2008) using primers corresponding to each gene either by amplification with Phusion or Q5 polymerases (NEB) and ligation or isothermal assembly. Amplicons were introduced into the appropriately cleaved plasmid and verified by PCR, restriction enzyme digestion and sequencing.

### Site specific mutagenesis

A slightly modified QuikChange mutagenesis protocol from Stratagene was used to engineer site specific mutations in *uppG* and *uppH*. Briefly, two complementary primers were designed with the required base pair mutation(s) flanked by ∼15 nt wildtype sequences on either side. A PCR reaction (16 cycles) using NEB Phusion polymerase was used to generate cut plasmid derivatives with the modified sequences. Correct amplification was verified via gel electrophoresis. 1 µl of the methylation-dependent restriction enzyme *Dpn*I was used to digest the methylated wildtype plasmid. Digested product was transformed into *E. coli* and plasmid clones were purified and sequenced to confirm the presence of the desired mutation.

### In-frame markerless deletions

In-frame markerless deletions were constructed as previously described (Morton & Fuqua, 2012a, Xu *et al*., 2013). Upstream and downstream ∼ 500 bp regions flanking the genes to be deleted were amplified from *A. tumefaciens* C58 genomic DNA using specific primers (P1 and P2 for upstream regions, P3 and P4 for downstream regions) and NEB Phusion polymerase. Care was taken to ensure the 5’ and 3’ ends of the genes were not altered, and that translational coupling between linked genes was retained. Primers P2 and P3 were designed with reverse complementarity in the 5’ sequences allowing for single overlap extension (SOE) amplification of both genes into a single amplified product as described (Morton & Fuqua, 2012a). The combined flanking regions were ligated into the appropriately cut suicide vector pNPTS138 using isothermal assembly with the NEBuilder HiFi DNA Assembly Cloning Kit to create a deletion construct which was subsequently transformed into chemically competent S17-1/λpir *E. coli*. The pNPTS138 vector confers kanamycin resistance (Km^R^) and sucrose sensitivity (Suc^S^) due to the presence of *sacB*. Introduction of this plasmid into *A. tumefaciens* was via conjugation with S17-λpir *E. coli*. The ColE1 origin of pNPTS138 does not replicate in *A. tumefaciens*, and a single crossover event allows plasmid integration into the chromosome, which was confirmed by patching transformants on to ATGN-Km and ATSN to screen for Km^R^Suc^S^ strains. Strains that had excised the plasmid were isolated by parallel patching on to ATSN and ATGN-Km to screen for Km^S^Suc^R^ derivatives. Deletion of the target genes was confirmed by diagnostic PCR using primers (P5 and P6) flanking the site of deletion.

### Congo red growth assays

*A. tumefaciens* strains were grown in ATGN until mid-exponential phase, normalized to an OD_600_ of 0.05, and 2 µl spotted on to ATGN-CR plates with or without IPTG. Plates were incubated at 30°C. Photographs of spots were taken following incubation for 48 h.

### Biofilm assays

Strains of *A. tumefaciens* were grown and analyzed as previously described (Morton & Fuqua, 2012c). Briefly, PVC coverslips were inserted vertically into the wells of 12-well polystyrene plates (Corning Inc.) and UV sterilized. The wells were subsequently filled with 3 ml mid-exponential phase cultures (with or without IPTG) normalized to OD_600_ 0.05. The 12-well plates were incubated in static conditions for 48 or 72 h at room temperature or 30°C inside a plastic Tupperware containing an open bottle of saturated K_2_SO_4_ solution to maintain humidity. Following incubation, the coverslips were rinsed with sterile water and stained with 0.1% w/v crystal violet (CV) solution 0.1% w/v for 20 min. Stained coverslips were rinsed with sterile water and then submerged in weigh boats containing 1 ml 33% acetic acid to solubilize the CV. Quantification was performed by measuring soluble A_600_ CV absorbance using a Biotek Synergy HT microplate reader and Gen 5 software (Biotek, Winooski, VT). Values were normalized to culture growth by dividing by final culture OD_600_.

### UPP production assays

UPP was visualized using the WGA lectin conjugated to Alexa fluor 594 (af-WGA, Invitrogen) and FITC (fl-DBA). 1 ml of mid-exponential phase *A. tumefaciens* cultures were spun down (7000 x g), washed twice with ATGN, pelleted, and resuspended in 100 µl ATGN. 1 µl of 1 mg/ml the fluorescent lectin stock was added to the suspensions and incubated at 28°C for 20-40 min. 1-2 µl labeled suspensions were placed on microscope slides containing agarose pads (1-1.5% agarose, 100-200 µl). Coverslips were placed on top of the pads, and samples were visualized by fluorescence microscopy using either a Nikon E800 camera with a 100X oil immersion objective or a Nikon Eclipse Ti-2 microscope with a Hamamatsu ORCAFlash 4.0 camera and a 60X oil immersion objective and NIS-Elements Software. Images were analyzed with Nikon Elements software and Fiji (Schindelin *et al*., 2012).

### Short term attachment and lectin binding assays

Short term binding assays were carried out as previously described (Morton & Fuqua, 2012c). Briefly, glass coverslips were placed horizontally inside the wells of 6-well plates and submerged in 2 ml mid-exponential phase cultures of *A. tumefaciens* strains. Plates were incubated statically 2 h at room temperature to allow cells to swim to and bind the coverslip. Following incubation, cultures were aspirated, and coverslips washed twice in ATGN to remove unattached cells. Coverslips were then placed over 100 µl of 1:100 dilution (10 μg/ml final) fl-WGA, afWGA, or fl-DBA, or two in combination for dual lectin binding. Coverslips were allowed to stain 1 h, then rinsed twice in ATGN before placing them either on a microscope slide or an agarose pad (1-1.5% agarose, 100-200 µl). Visualization was performed as with standard UPP production assays. For high resolution single cell microscopy, spinning disk confocal microscopy (SDCM) was used (Yokogawa CSU10 confocal, Nikon TE2000U microscope, Photometrics Cascade II 512B camera) with Metamorph software (Molecular Devices Corp., Sunnyvale, CA).

### Electron microscopy

Electron microscopy was carried out at the Indiana Molecular Biology Institute Microscopy Center. For Transmission Electron Microscopy (TEM) and colloidal gold-labeled lectin, 1 ml of mid-exponential phase culture was pelleted and resuspended in 200 µl ATGN. 5 µl of 20 mg/ml suspensions of 20 nm colloidal gold conjugated WGA or 10 nm gold conjugated PNA (EY Laboratories) were added to the suspension and incubated at room temperature for 20 min. The labeled suspension was pelleted at 6000 x g and washed twice with 1 ml 10 mM Tris-Cl, pH 8.0 before resuspending in 10 mM Tris-Cl, pH 8.0. 10 µl of the resuspension were placed on a formvar-coated copper grid for 5 min and stained with 10 µl of 1% uranyl acetate for another 5 min. Filter paper was used to remove the excess uranyl acetate, and the grid was air dried for 5 min before viewing using the JEOL JEM-1010 Transmission Electron Microscope at 80 kV.

### Purification of a His_6_-UppH protein

The *uppH* coding sequence (Atu2371) was amplified from genomic DNA and fused downstream of the His_6_-tag coding sequence in pET-15b cleaved at the *Nde*I and *Bam*HI restriction sites. The resulting plasmid pRN39 was transformed into chemically competent *E. coli* BL21-CodonPlus (DE3)-RIL cells (Stratagene) and grown at 37°C with shaking in 200 ml LB supplemented with Ap (100 µg/ml). At mid-exponential phase, His_6_-UppH expression was induced by addition of 200 μM IPTG and incubating for 4 h. Cells were collected by centrifugation at 5,600 x g for 20 minutes and stored at -20°C. 2 g of cells were thawed on ice, resuspended in 5 ml extraction buffer (50 mM TES [(N-tris(hydroxymethyl)methyl-2-aminoethanesulfonic acid)], 10 mM MgCl_2_,10 mM DTT, pH 7.0) and lysed using an M-110L Microfluidizer Processor (Microfluidics, Westwood, MA). Following 10 min centrifugation (9,900 x g), a significant fraction of the overexpressed tagged UppH was insoluble, but a fraction remained soluble based on SDS-PAGE analysis. The supernatant with the soluble His_6_-UppH was loaded on to a 3 ml TALON Metal Affinity Resin (Clontech) equilibrated in binding buffer (50 mM Na_2_HPO_4_, 300 mM NaCl, 10% glycerol, 10 mM imidazole, pH 8). 4 ml of binding buffer containing increasing concentrations of imidazole (50 mM, 250 mM, 500 mM, and 1M) were used to carry out step gradient elution. UppH was present in the 250 mM imidazole fraction based on SDS-PAGE analysis and MALDI-MS of the excised band. The purified protein was dialyzed into 50 mM Na_2_HPO_4_, 100 mM NaCl, 10% glycerol, pH 8. Protein concentration was determined by the molar absorptivity using bovine serum albumin as a standard and the enzyme was aliquoted and stored at -80°C for future use. The same approach was used to purify His_6_-UppH E152Q E266Q.

### Transposon mutagenesis and *en-masse* sequencing

*Mutagenesis:* donor *E. coli* SM10-λpir carrying the Himar1 plasmid pFD1 was grown in LB overnight until turbid. Recipient *A. tumefaciens* strains CDGS-Upp+ Δ*pruA* Δ*uppY* and CDGS-Upp+ Δ*pruA* Δ*uppW* were also grown in LB until turbid. Turbid cultures were subcultured 1:10 in fresh LB and incubated until they reached mid-exponential phase (∼4-6 h). 1 ml each of the donor and recipient strains were collected in Eppendorf tubes, spun down at 5,600 x g for 1 min, and the pellets resuspended in 50 µl LB. Subsequently, 50 µl of the donor strain was combined with 50 µl of each of the recipient strains. Entire matings (100 µl) were spotted on 0.2 µm cellulose acetate filter paper on LB plates. Plates were incubated overnight at 28°C to allow for conjugative transfer. Cells were resuspended from the filter paper in 30% glycerol, serially diluted in 1/10 increments, and frozen at -80°.

*Screening and gDNA extraction:* To optimize for visualization of CR phenotypes, solid medium was prepared using 1.5 ml of 2% stock solution of CR per 100 ml ATGN. Plates also contained 300 µg/ml Km for selection. 100 µl of cell suspensions were plated on ATGN-CR-Kan plates at 1:100 dilution for CDGS-Upp+ Δ*pruA* Δ*uppY*, and 1:1000 dilution for CDGS-Upp+ Δ*pruA* Δ*uppW*. Plates were incubated 48-72 h and screened for decreased CR (DCR) colony phenotypes. DCR colonies from the different strain backgrounds were picked, streaked or patched on fresh ATGN-CR-Kan plates and incubated at 28°C. PCR was used to confirm that the DCR mutants were *A. tumefaciens* and not Km^R^ contaminants with a DCR phenotype. Biofilm assays on randomly selected DCR mutants showed deficiencies to varying degrees. Confirmed DCR strains were inoculated in 2 ml ATGN-Kan, and 250 µl from each stationary-phase culture was pooled together into 1 ml batches. This was done separately for the two *A. tumefaciens* strain backgrounds. gDNA was extracted from pooled DCR cultures from the two strain backgrounds using (Promega gDNA extraction kit) and stored at 4°C. Over 50,000 colonies were screened collectively in both strain backgrounds, and roughly 0.5% were DCR mutants in each background.

*Sequencing library preparation:* Illumina library construction was carried out for the two separate gDNA pools using the terminal deoxynucleotidyl transferase (TdT) tailing method (Deng and Wu, 1983). Briefly, a Covaris instrument was used to shear gDNA to 200-700 bp fragments, and a Tape Station instrument was used to estimate sheared gDNA concentration (for this protocol, gDNA concentration was at least 69 ng/µl). Several consecutive PCR reactions were carried out to perform TdT tailing, to append adaptors sequences needed for Illumina sequencing, and to append adaptors with unique barcodes to the separate libraries corresponding to the two *A. tumefaciens* strain backgrounds. Final reaction products were combined to be multiplexed and the library was further purified by passing through a PCR purification column (Edge Biosystems).

The library was submitted to the IU Center for Genomics and Bioinformatics (CGB) for sequencing. Sequencing was performed on a NextSeq 500 using a 75-cycle single-end run. Reads were then mapped to the *Agrobacterium tumefaciens* C58 reference genome using bowtie2 (v2.3.5.1) with the following special parameters: --local -k 2. Only reads that originated within the transposon (and started with AACCTGTTA allowing a single mismatch) and had uniquely mapped were kept and tallied to quantify transposon insertion rates.

### Multi-locus sequence analysis (MLSA)

Publicly available genome sequences from the *Agrobacterium*/*Rhizobium* group as well as select *Sinorhizobium*, *Ensifer*, and *Kaistia* strains were downloaded from NCBI GenBank on September 12, 2019. AutoMLSA with the default parameters and IQ-TREE v. 1.6.12 with the parameters “-bb 1000 -alrt 1000" were used to generate a multi-locus sequence analysis (MLSA) tree (Davis *et al*., 2016, Nguyen *et al*., 2015). Twenty-three translated gene sequences (d*naK, prfC, hemN, hamI, truA, glyS, coxC*, ATU_RS03855, ATU_RS07710, *hom, rplB, rpoC, rpoB, murC, hemF*, ATU_RS12850, *acnA, leuS, cgtA, secA, plsC, aroB, lysC*) from the strain *A. fabrum* C58 (NCBI accession: GCA_000092025.1) were used as reference sequences for autoMLSA (Zhang *et al*., 2012). Protein sequences from the genome of *A. tumefaciens* (fabrum) C58 for the loci ATU_RS02360, ATU_RS02370, ATU_RS06100, ATU_RS06105, ATU_RS06110, ATU_RS06115, ATU_RS06120, ATU_RS06125, ATU_RS06130, ATU_RS11495, ATU_RS11500, ATU_RS11550, ATU_RS11555, ATU_RS11560, ATU_RS11565, ATU_RS11570, ATU_RS11575, ATU_RS11580, ATU_RS11585, Atu2378 (no annotation in NCBI), ATU_RS11590, ATU_RS16315, ATU_RS16320, ATU_RS17560, ATU_RS17565, ATU_RS17570, ATU_RS19030, ATU_RS19035, ATU_RS19040, ATU_RS26530, and ATU_RS16325 were used as queries in BLASTP v. 2.6.0+ searches of individual genomes; and the best blast hit was taken (Camacho *et al*., 2009). Top hit sequences were then used as queries in BLASTP searches against a database of proteins from the C58 reference genome. Blast hits were filtered to those with >50% query coverage and >40% amino acid ID, and the best hit to C58 were identified. If the best hits of both searches were reciprocal best hits, the presence of that gene was confirmed in a genome. Phylogenies and heat maps were visualized using the R package ggtree (Yu *et al*., 2017). Gene synteny figures were generated using clinker v. 0.0.9 with the default options (Gilchrist & Chooi, 2021).

## Supporting information

Supplementary Infoamation

## Acknowledgements

We wish to acknowledge and Loralyn Cozy and Megan Lohmiller for help with initial work on the UPP, and Yves Brun and Ellen Quardokus for helpful discussions and suggestions. Ankur Dalia generously provided protocols and troubleshooting advice for transposon sequencing. Barry Stein was a great help with electron microscopy, performed at the Indiana University (IU) Electron Microscopy Center. We thank the Department of Botany and Plant Pathology at Oregon State University for its support of computing infrastructure. T.D. was supported through an Indiana University-Purdue University Collaborations in Life Science and Informatics Research grant. M.E.H. received support from the IU Genetics, Cellular and Molecular Sciences NIH Training Grant (T32 GM007757). This project was supported by grants to C.F. from the NIH (RO1 GM120337), and the Indiana University META-Cyt Program funded in part by a major endowment from the Lilly Foundation, and to J.H.C. by the National Science Foundation (DEB-1738028). The funders had no role in study design, data collection and analysis, decision to publish, or preparation of the manuscript.

## Author contributions

C.F. formulated the research plan; M.O., R.N., G.H., J.X., I.R., J.K., T.D., M.H. and C.F. performed the experiments, J.C. and A.W. performed genomic sequence analyses; C.F., M.O. and G.H. wrote the paper.

## Data availability

High-throughput, Illumina-based transposon sequencing data is being deposited in the GEO database (([dataset] Onyeziri and Fuqua, 2021). All other data that support the findings of this study are available from the corresponding author upon reasonable request.

## References

Anderson-Furgeson, J.C., Zupan, J.R., Grangeon, R., and Zambryski, P.C. (2016) Loss of PodJ in *Agrobacterium tumefaciens* leads to ectopic polar growth, branching, and reduced cell division. J Bacteriol 198: 1883–1891.

Baker, P., Hill, P.J., Snarr, B.D., Alnabelseya, N., Pestrak, M.J., Lee, M.J., Jennings, L.K., Tam, J., Melnyk, R.A., Parsek, M.R., Sheppard, D.C., Wozniak, D.J., and Howell, P.L. (2016) Exopolysaccharide biosynthetic glycoside hydrolases can be utilized to disrupt and prevent *Pseudomonas aeruginosa* biofilms. Science advances 2: e1501632.

Berne, C., Ellison, C.K., Ducret, A., and Brun, Y.V. (2018) Bacterial adhesion at the single-cell level. Nat Rev Microbiol 16: 616–627.

Braun, A.C., and Elrod, R.P. (1946) Stages in the life history of *Phytomonas tumefaciens*. J Bacteriol. 52: 695–702.

Breton, C., Snajdrova, L., Jeanneau, C., Koca, J., and Imberty, A. (2006) Structures and mechanisms of glycosyltransferases. Glycobiology 16: 29R–37R.

Brown, P.J., de Pedro, M.A., Kysela, D.T., Van der Henst, C., Kim, J., De Bolle, X., Fuqua, C., and Brun, Y.V. (2012) Polar growth in the Alphaproteobacterial order Rhizobiales. Proc Natl Acad Sci U S A 109: 1697–1701.

Camacho, C., Coulouris, G., Avagyan, V., Ma, N., Papadopoulos, J., Bealer, K., and Madden, T.L. (2009) BLAST+: architecture and applications. BMC Bioinformatics 10: 421.

Cangelosi, G.A., Hung, L., Puvanesarajah, V., Stacey, G., Ozga, D.A., Leigh, J.A., and Nester, E.W. (1987) Common loci for *Agrobacterium tumefaciens* and *Rhizobium meliloti* exopolysaccharide synthesis and their roles in plant interactions. J. Bacteriol. 169: 2086–2091.

Collins, R.F., Beis, K., Dong, C., Botting, C.H., McDonnell, C., Ford, R.C., Clarke, B.R., Whitfield, C., and Naismith, J.H. (2007) The 3D structure of a periplasm-spanning platform required for assembly of group 1 capsular polysaccharides in *Escherichia coli*. Proc Natl Acad Sci U S A 104: 2390–2395.

Curtis, P.D., and Brun, Y.V. (2014) Identification of essential alphaproteobacterial genes reveals operational variability in conserved developmental and cell cycle systems. Mol Microbiol 93: 713–735.

Cuthbertson, L., Mainprize, I.L., Naismith, J.H., and Whitfield, C. (2009) Pivotal roles of the outer membrane polysaccharide export and polysaccharide copolymerase protein families in export of extracellular polysaccharides in gram-negative bacteria. Microbiol Mol Biol Rev 73: 155–177.

Davis, E.W., Weisberg, A.J., Tabima, J.F., Grunwald, N.J., and Chang, J.H. (2016) Gall-ID: tools for genotyping gall-causing phytopathogenic bacteria. PeerJ 4: e2222.

De Castro, C., Molinaro, A., Lanzetta, R., Silipo, A., and Parrilli, M. (2008) Lipopolysaccharide structures from *Agrobacterium* and *Rhizobiaceae* species. Carbohydr Res 343: 1924–1933.

Feirer, N., Kim, D., Xu, J., Fernandez, N., Waters, C.M., and Fuqua, C. (2017) The *Agrobacterium tumefaciens* CheY-like protein ClaR regulates biofilm formation. Microbiology 163: 1680–1691.

Feirer, N., Xu, J., Allen, K.D., Koestler, B.J., Bruger, E.L., Waters, C.M., White, R.H., and Fuqua, C. (2015) A pterin-dependent signaling pathway regulates a dual-function diguanylate cyclase-phosphodiesterase controlling surface attachment in *Agrobacterium tumefaciens*. mBio 6: e00156.

Fiebig, A., Herrou, J., Fumeaux, C., Radhakrishnan, S.K., Viollier, P.H., and Crosson, S. (2014) A cell cycle and nutritional checkpoint controlling bacterial surface adhesion. PLoS Genet 10: e1004101.

Flemming, H.C., Wingender, J., Szewzyk, U., Steinberg, P., Rice, S.A., and Kjelleberg, S. (2016) Biofilms: an emergent form of bacterial life. Nat Rev Microbiol 14: 563–575.

Fritts, R.K., LaSarre, B., Stoner, A.M., Posto, A.L., and McKinlay, J.B. (2017) A Rhizobiales-specific unipolar polysaccharide adhesin contributes to *Rhodopseudomonas palustris* biofilm formation across diverse photoheterotrophic conditions. Appl Environ Microbiol 83.

Fuqua, C., (2008) Agrobacterium-host attachment and biofilm formation. In: Agrobacterium: From Biology to Biotechnology. T. Tzfira & V. Citovsky (eds). New York, NY: Springer Science + Business Media LLC, pp. 243–277.

Gelvin, S.B. (2003) *Agrobacterium*-mediated plant transformation: the biology behind the "gene-jockeying" tool. Microbiol. Mol. Biol. Rev. 67: 16–37.

Gilchrist, C.L.M., and Chooi, Y.H. (2021) Clinker & clustermap.js: Automatic generation of gene cluster comparison figures. Bioinformatics.

Ginsberg, C., Zhang, Y.H., Yuan, Y., and Walker, S. (2006) In vitro reconstitution of two essential steps in wall teichoic acid biosynthesis. ACS Chem Biol 1: 25–28.

Goodner, B., Hinkle, G., Gattung, S., Miller, N., Blanchard, M., Qurollo, B., Goldman, B.S., Cao, Y., Askenazi, M., Halling, C., Mullin, L., Houmiel, K., Gordon, J., Vaudin, M., Iartchouk, O., Epp, A., Liu, F., Wollam, C., Allinger, M., Doughty, D., Scott, C., Lappas, C., Markelz, B., Flanagan, C., Crowell, C., Gurson, J., Lomo, C., Sear, C., Strub, G., Cielo, C., and Slater, S. (2001) Genome sequence of the plant pathogen and biotechnology agent *Agrobacterium tumefaciens* C58. Science 294: 2323–2328.

Hardy, G.G., Allen, R.C., Toh, E., Long, M., Brown, P.J., Cole-Tobian, J.L., and Brun, Y.V. (2010) A localized multimeric anchor attaches the *Caulobacter* holdfast to the cell pole. Mol Microbiol 76: 409–427.

Heckel, B.C., Tomlinson, A.D., Morton, E.R., Choi, J.H., and Fuqua, C. (2014) *Agrobacterium tumefaciens exoR* controls acid response genes and impacts exopolysaccharide synthesis, horizontal gene transfer, and virulence gene expression. J Bacteriol 196: 3221–3233.

Hernando-Perez, M., Setayeshgar, S., Hou, Y., Temam, R., Brun, Y.V., Dragnea, B., and Berne, C. (2018) Layered structure and complex mechanochemistry underlie strength and versatility in a bacterial adhesive. mBio 9.

Hershey, D.M., Porfirio, S., Black, I., Jaehrig, B., Heiss, C., Azadi, P., Fiebig, A., and Crosson, S. (2019) Composition of the holdfast polysaccharide from *Caulobacter crescentus*. J Bacteriol 201.

Hinz, A.J., Larson, D.E., Smith, C.S., and Brun, Y.V. (2003) The *Caulobacter crescentus* polar organelle development protein PodJ is differentially localized and is required for polar targeting of the PleC development regulator. Mol Microbiol. 47: 929–941.

Holden, H.M., Rayment, I., and Thoden, J.B. (2003) Structure and function of enzymes of the Leloir pathway for galactose metabolism. J Biol Chem 278: 43885–43888.

Hong, Y., Liu, M.A., and Reeves, P.R. (2018) Progress in our understanding of Wzx flippase for translocation of bacterial membrane lipid-linked oligosaccharide. J Bacteriol 200.

Hong, Y., Morcilla, V.A., Liu, M.A., Russell, E.L.M., and Reeves, P.R. (2015) Three Wzy polymerases are specific for particular forms of an internal linkage in otherwise identical O units. Microbiology (Reading) 161: 1639–1647.

Javens, J., Wan, Z., Hardy, G.G., and Brun, Y.V. (2013) Bypassing the need for subcellular localization of a polysaccharide export-anchor complex by overexpressing its protein subunits. Molecular Microbiology 89: 350–371.

Kamoun, S., Cooley, M.B., Rogowsky, P.M., and Kado, C.I. (1989) Two chromosomal loci involved in production of exopolysaccharide in *Agrobacterium tumefaciens*. J Bacteriol 171: 1755–1759.

Khan, S.R., Gaines, J., Roop, R.M., 2nd, and Farrand, S.K. (2008) Broad-host-range expression vectors with tightly regulated promoters and their use to examine the influence of TraR and TraM expression on Ti plasmid quorum sensing. Appl Environ Microbiol 74: 5053–5062.

Lassalle, F., Campillo, T., Vial, L., Baude, J., Costechareyre, D., Chapulliot, D., Shams, M., Abrouk, D., Lavire, C., Oger-Desfeux, C., Hommais, F., Gueguen, L., Daubin, V., Muller, D., and Nesme, X. (2011) Genomic species are ecological species as revealed by comparative genomics in *Agrobacterium tumefaciens*. Genome Biol Evol 3: 762–781.

Laus, M.C., Logman, T.J., Lamers, G.E., Van Brussel, A.A., Carlson, R.W., and Kijne, Y.V. (2006) A novel polar surface polysaccharide from *Rhizobium leguminosarum* binds host plant lectin. Mol Microbiol. 59: 1704–1713.

Li, G., Brown, P.J., Tang, J.X., Xu, J., Quardokus, E.M., Fuqua, C., and Brun, Y.V. (2012) Surface contact stimulates the just-in-time deployment of bacterial adhesins. Mol. Microbiol. 83: 41–51.

Li, Y.G., and Christie, P.J. (2018) The *Agrobacterium* VirB/VirD4 T4SS: mechanism and architecture defined through *in vivo* mutagenesis and chimeric systems. Curr Top Microbiol Immunol 418: 233–260.

Lombard, V., Golaconda Ramulu, H., Drula, E., Coutinho, P.M., and Henrissat, B. (2014) The carbohydrate-active enzymes database (CAZy) in 2013. Nucleic Acids Res 42: D490–495.

Mathews, S.L., Hannah, H., Samagaio, H., Martin, C., Rodriguez-Rassi, E., and Matthysse, A.G. (2019) Glycoside hydrolase genes are required for virulence of *Agrobacterium tumefaciens* on *Bryophyllum daigremontiana* and tomato. Appl Environ Microbiol 85.

Morton, E.R., and Fuqua, C. (2012a) Genetic manipulation of Agrobacterium. Curr Protoc Microbiol: Unit 3D 2.

Morton, E.R., and Fuqua, C. (2012b) Laboratory maintenance of Agrobacterium. Curr Protoc Microbiol Chapter 1: Unit3D 1.

Morton, E.R., and Fuqua, C. (2012c) Phenotypic analyses of Agrobacterium. Curr Protoc Microbiol Chapter 3: Unit 3D 3.

Nguyen, L.T., Schmidt, H.A., von Haeseler, A., and Minh, B.Q. (2015) IQ-TREE: a fast and effective stochastic algorithm for estimating maximum-likelihood phylogenies. Mol Biol Evol 32: 268–274.

Oglesby, L.L., Jain, S., and Ohman, D.E. (2008) Membrane topology and roles of *Pseudomonas aeruginosa* Alg8 and Alg44 in alginate polymerization. Microbiology (Reading) 154: 1605–1615.

[dataset] Onyeziri, M.C. and Fuqua, C. Gene Expression Omnibus (GE), GEO identifier pending. All other data that support the findings of this study are available from the corresponding author upon reasonable request.

Poindexter, J.S. (1981) The caulobacters: ubiquitous unusual bacteria. Microbiol. Rev. 45: 123–179.

Reuber, T.L., and Walker, G.C. (1993) The acetyl substituent of succinoglycan is not necessary for alfalfa nodule invasion by *Rhizobium meliloti* Rm1021. J Bacteriol 175: 3653–3655.

Romling, U., and Galperin, M.Y. (2015) Bacterial cellulose biosynthesis: diversity of operons, subunits, products, and functions. Trends Microbiol 23: 545–557.

Ruan, X., Monjaras Feria, J., Hamad, M., and Valvano, M.A. (2018) *Escherichia coli* and *Pseudomonas aeruginosa* lipopolysaccharide O-antigen ligases share similar membrane topology and biochemical properties. Mol Microbiol 110: 95–113.

Schindelin, J., Arganda-Carreras, I., Frise, E., Kaynig, V., Longair, M., Pietzsch, T., Preibisch, S., Rueden, C., Saalfeld, S., Schmid, B., Tinevez, J.Y., White, D.J., Hartenstein, V., Eliceiri, K., Tomancak, P., and Cardona, A. (2012) Fiji: an open-source platform for biological-image analysis. Nat Methods 9: 676–682.

Slater, S.C., Goldman, B.S., Goodner, B., Setubal, J.C., Farrand, S.K., Nester, E.W., Burr, T.J., Banta, L., Dickerman, A.W., Paulsen, I., Otten, L., Suen, G., Welch, R., Almeida, N.F., Arnold, F., Burton, O.T., Du, Z., Ewing, A., Godsy, E., Heisel, S., Houmiel, K.L., Jhaveri, J., Lu, J., Miller, N.M., Norton, S., Chen, Q., Phoolcharoen, W., Ohlin, V., Ondrusek, D., Pride, N., Stricklin, S.L., Sun, J., Wheeler, C., Wilson, L., Zhu, H., and Wood, D.W. (2009) Genome sequences of three agrobacterium biovars help elucidate the evolution of multichromosome genomes in bacteria. J Bacteriol 191: 2501–2511.

Tempé, J., Petit, A., Holsters, M., Van Montagu, M., and Schell, J. (1977) Thermosensitive step associated with transfer of the Ti plasmid during conjugation: possible relation to transformation in crown gall. Proc. Natl. Acad. Sci. USA 74: 2848–2849.

Thompson, M.A., Onyeziri, M.C., and Fuqua, C. (2018) Function and regulation of *Agrobacterium tumefaciens* cell surface structures that promote attachment. Curr Top Microbiol Immunol 418: 143–184.

Toh, E., Kurtz, H.D., Jr., and Brun, Y.V. (2008) Characterization of the *Caulobacter crescentus* holdfast polysaccharide biosynthesis pathway reveals significant redundancy in the initiating glycosyltransferase and polymerase steps. J Bacteriol. 190: 7219–7231.

Tomlinson, A.D., and Fuqua, C. (2009) Mechanisms and regulation of polar surface attachment in *Agrobacterium tumefaciens*. Curr Opin Microbiol 12: 708–714.

Vetting, M.W., LP, S.d.C., Yu, M., Hegde, S.S., Magnet, S., Roderick, S.L., and Blanchard, J.S. (2005) Structure and functions of the GNAT superfamily of acetyltransferases. Arch Biochem Biophys 433: 212–226.

Vuong, C., Kocianova, S., Voyich, J.M., Yao, Y., Fischer, E.R., DeLeo, F.R., and Otto, M. (2004) A crucial role for exopolysaccharide modification in bacterial biofilm formation, immune evasion, and virulence. J Biol Chem 279: 54881–54886.

Wan, Z., Brown, P.J., Elliott, E.N., and Brun, Y.V. (2013) The adhesive and cohesive properties of a bacterial polysaccharide adhesin are modulated by a deacetylase. Mol Microbiol 88: 486–500.

Weisberg, A.J., Davis, E.W.mTabima, J., Belcher, M.S., Miller, M., Kuo, C.H., Loper, J.E., Grunwald, N.J., Putnam, M.L., and Chang, J.H. (2020) Unexpected conservation and global transmission of agrobacterial virulence plasmids. Science 368.

Whitfield, C., Wear, S.S., and Sande, C. (2020) Assembly of bacterial capsular polysaccharides and exopolysaccharides. Annu Rev Microbiol 74: 521–543.

Williams, A., Wilkinson, A., Krehenbrink, M., Russo, D.M., Zorreguieta, A., and Downie, J.A. (2008) Glucomannan-mediated attachment of *Rhizobium leguminosarum* to pea root hairs is required for competitive nodule infection. J Bacteriol. 190: 4706–4715.

Wood, D.W., Setulab, J.C., Kaul, R., Monks, D.E., Kitajima, J.P., Okura, V.K., Zhou, Y., Chen, L., Wood, G.E., Almeida, N.F.J., Woo, L., Chen, Y., Paulsen, I.T., Eisen, J.A., Karp, P.D., Dovee, D.S., Chapman, P., Clendenning, J., Deatherage, G., Gillet, W., Grant, C., Kutyavin, T., Levy, R., Li., M.-J., McClellund, E., Palmieri, A., Raymond, C., Rouse, G., Saenphimmachak, C., Wu, Z., Romero, P., Gordon, D., Zhang, S., Yoo, H., Tao, Y., Biddle, P., Jung, M., Krespan, W., Perry, M., Gordon-Kamm, B., Liao, L., Kim, S., Hendrick, C., Zhao, Z.-Y., Dolan, M., Chumley, F., Tingey, S.V., Tomb, J.-F., Gordon, M., Olson, M.V., and Nester, E.W. (2001) The genome of the natural genetic engineer *Agrobacterium tumefaciens* C58. Science 294: 2317–2323.

Xu, J., Kim, J., Danhorn, T., Merritt, P.M., and Fuqua, C. (2012) Phosphorus limitation increases attachment in *Agrobacterium tumefaciens* and reveals a conditional functional redundancy in adhesin biosynthesis. Res Microbiol 163: 674–684.

Xu, J., Kim, J., Koestler, B.J., Choi, J.H., Waters, C.M., and Fuqua, C. (2013) Genetic analysis of *Agrobacterium tumefaciens* unipolar polysaccharide production reveals complex integrated control of the motile-to-sessile switch. Mol Microbiol 89: 929–948.

Yildiz, F., Fong, J., Sadovskaya, I., Grard, T., and Vinogradov, E. (2014) Structural characterization of the extracellular polysaccharide from *Vibrio cholerae* O1 El-Tor. PLoS One 9: e86751.

Yu, G., Smith, D.K., Zhu, H., Guan, Y., and Lam, T.T.-Y. (2017) ggtree: an r package for visualization and annotation of phylogenetic trees with their covariates and other associated data. *Meth*. Ecol. Evol. 8: 28–36.

Zhang, H.B., Wang, C., and Zhang, L.H. (2004) The quormone degradation system of *Agrobacterium tumefaciens* is regulated by starvation signal and stress alarmone (p)ppGpp. Mol Microbiol 52: 1389–1401.

Zhang, Y.M., Tian, C.F., Sui, X.H., Chen, W.F., and Chen, W.X. (2012) Robust markers reflecting phylogeny and taxonomy of rhizobia. PLoS One 7: e44936.

