## Supplementary Infoamation for "Dual Adhesive Unipolar Polysaccharides Synthesized by Overlapping Biosynthetic Pathways in *Agrobacterium tumefaciens*"

Supplementary Tables: S1-S4

Supplementary References:

Supplementary Figure Legends:

Supplementary Figures: S1-S8

**Table S1. Putative UPP Biosynthetic Genes and Proteins – Properties and Mutant Phenotypes**

| ATU RS Number (Atu) | Gene Name | Protein Similarity | P-sach Nom.† | Length (aa) | Local.‡ | Motifs | ΔMutant %WT (S.D.) (Comp.)§ | Mutant Lectin (WGA) |
|---|---|---|---|---|---|---|---|---|
| 06125 (Atu1240) | *uppA* | GNAT | BcsL | 408 | Cyt | | 12 (1) (+) | None |
| 06120 (Atu1239) | *uppB* | GumC MRP | Wzc (PCP-2a) | 753 | CM | 2 TM | 10 (2) (+) | None |
| 06115 (Atu1238) | *uppC* | Secretin | Wza (OPX) | 190 | OM | β-barrel Lipidation Site (23) | 2 (<1) (+/-) *uppCD*(+) | None |
| 06110 (Atu1237) | *uppD* | GT4 PimA-like | NA | 382 | Cyt | GT Cat | 11 (<1) (+) | None |
| 06105 (Atu1236) | *uppE* | PHPT | WbaP | 457 | CM | 3 | 3 (+) | None |
| 06100 (Atu1235) | *uppF* | RfaL O-antigen Ligase | Wzy_C | 413 | CM | 12 TM | 6 (+) | None |
| 11550 (Atu2370) | *uppG* | GT4 | NA | 361 | Cyt | GT Cat., DxD (142-4) | 33(8) (+) | Small |
| 11555 (Atu2371) | *uppH* | GH10 XynA | NA | 355 | Peri | N-term Tat SS GH Cat. E152, E266 | 101 (18) (NA) | Normal |
| 11560 (Atu2372) | *uppI* | GT2 | NA | 315 | Cyt | GT Cat. | 44 (13) (+) | ND |
| 11565 (Atu2373) | *uppJ* | GT2 | NA | 314 | Cyt | GT Cat. | 13 (6) (+) | None |
| 11570 (Atu2374) | *uppK* | Con Hyp. | NA | 113 | Cyt | 2 TM | 92 (25) (NA) | ND |
| 11585 (Atu2377) | *uppM* | GumC | Wzc (PCP-2a) | 476 | CM | 2 TM | 39 (7) (+) | Weak |

| | | | | | | | | |
|---|---|---|---|---|---|---|---|---|
| No Annot. (Atu2378) | *uppN* | Con Hyp. | | 59 | Cyt | | 2 (<1) (-) | None |
| 11590 (Atu2379) | *uppO* | NoeK Phospho-mannomutase | | 483 | Cyt | | 177 (6) NA | Normal |
| 02360 (Atu0480) | *uppX* | RfbX Flippase | Wzx | 426 | CM | 12 TM | 8 (5) (+) | None |
| 02370 (Atu0481) | *uppY* | RfaL Polymerase | Wzy | 431 | CM | 10 TM | 104 (18) (NA) | Normal |
| 16320 (Atu3523) | *uppV* | RfbX Flippase | Wzx | 477 | CM | 12 TM | 37(22) (+) | None |
| 16325 (Atu8188) | *uppQ* | GNAT | BcsL | 423 | Cyt | | 4 (<1) (+) | None |
| 11495 (Atu2356) | *uppW* | RfaL Polymerase | Wzy | 477 | CM | 10 TM | 126 (29) (NA) | None |
| 11500 (Atu2358) | *uppZ* | OafA AT | NA | 334 | CM | 10 TM | ND | ND |
| 17565 (Atu3777) | *uppP* | Outer Membrane Protein | NA | 497 | OM | β-Barrel Porin Type II | 3 (1) (+) | None |
| 19035 (Atu4074) | *exoC* | Phospho-glucomutase | NA | 542 | Cyt | | ND | None |
| 19470 (Atu4166) | *exoB* | GalE | NA | 328 | Cyt | | ND | ND |

†Polysaccharide biosynthesis nomenclature – see (Whitfield, 2006)

‡ Predicted cellular localization: Cyt, cytoplasm; CM, cytoplasmic membrane; Peri, periplasm; OM, outer membrane

§ Ratio of solubilized CV $A_{600}/OD_{600}$ values of in-frame deletion mutants to % wild type C58 (S.D. standard deviation) and complementation status. (+), full complementation; (+/-), partial complementation;

NA – not applicable; ND – not determined

**Table S2.** DCR Transposon Mutant Insertions

| Gene ID (Atu RS) | Annotated gene product | Number of Insertions Per ECR Mutant | | | | |
|---|---|---|---|---|---|---|
| | | Δ *claR* (ATU_ RS08010) | Δ*visR* (ATU_ RS02585) | Δ*pruA* (ATU_ RS05580) | Δ*dcpA* (ATU_ RS16190) | Total |
| 02360 | polysaccharide biosynthesis protein,wzx | 1 | 5 | 12 | 3 | 21 |
| 06105 | polysaccharide biosynthesis glycosyltransferase | 1 | 6 | 8 | 1 | 16 |
| 06110 | glycosyl transferase | 1 | 1 | 3 | 1 | 6 |
| 06115 | polysaccharide export protein | 3 | 3 | 7 | 1 | 14 |
| 06120 | exopolysaccharide production protein | 0 | 0 | 2 | 0 | 2 |
| 11550 | Glycosyl transferase | 2 | 0 | 5 | 0 | 7 |
| 11555 | endo-1,4-beta-xylanase | 2 | 2 | 3 | 0 | 7 |
| 11560 | UDP-hexose transferase | 4 | 3 | 1 | 1 | 9 |
| 11565 | UDP-hexose transferase | 0 | 0 | 3 | 0 | 3 |
| 11570 | hypothetical protein | 0 | 0 | 4 | 0 | 4 |
| 11575 | UDP-hexose transferase | 0 | 2 | 1 | 0 | 3 |
| 11585 | succinoglycan biosynthesis transport protein | 1 | 0 | 6 | 0 | 7 |
| No Annot. | hypothetical protein | 1 | 0 | 2 | 0 | 3 |
| 11590 | phosphomannomutase | 0 | 0 | 3 | 1 | 4 |
| 17560 | hypothetical protein | 1 | 0 | 0 | 0 | 1 |
| 17565 | conserved hypothetical protein | 1 | 0 | 6 | 1 | 8 |
| 16320 | polysaccharide biosynthesis protein,wzx | 0 | 0 | 1 | 1 | 2 |
| 16325 | GNAT acetyltransferase | 1 | 3 | 8 | 3 | 15 |
| 19035 | phosphoglucomutase, exoC | 3 | 1 | 4 | 0 | 8 |
| Total | | 22 | 26 | 79 | 13 | 140 |

**Table S3.** Pathway-specific genetic screen

| **ATU RS Number (Atu)** | | **Independent transposon mutants** | |
|---|---|---|---|
| | **Gene name** | **Δ*uppY1*** | **Δ*uppY2*** |
| **Polymerase Genes** | | | |
| 02370 (Atu0481) | *uppY* | 0 | 8 |
| 11495 (Atu2356) | *uppW* | 15 | 0 |
| **Isolated in both screens (annotated)** | | | |
| 06115 (Atu1238) | *uppC* | 13 | 17 |
| 06105 (Atu1236) | *uppE* | 13 | 7 |
| 11560 (Atu2372) | *uppI* | 9 | 8 |
| 11565 (Atu2373) | *uppJ* | 1 | 3 |
| 11575 (Atu2375) | *uppL* | 1 | 6 |
| 11585 (Atu2377) | *uppM* | 14 | 27 |
| No annotated (Atu2378) | *uppN* | 1 | 2 |
| 17565 (Atu3777) | *uppP* | 7 | 1 |
| 16325 (Atu8188) | *uppQ* | 9 | 7 |
| 02360 (Atu0480) | *uppX* | 17 | 14 |
| 16320 (Atu3523) | *uppV* | 3 | 1 |
| 19035 (Atu4047) | *exoC* | 7 | 21 |
| 19470 (Atu4166) | *galE/exoB* | 5 | 6 |
| 05105 (Atu1030) | *relA* | 5 | 9 |
| 07130 (Atu1447) | *ntrY* | 6 | 4 |
| 08400 (Atu1715) | *exoR* | 6 | 4 |
| 22800 (Atu4855) | *traA* | 1 | 2 |
| **Isolated in both screens (unannotated)** | | | |
| 15030 (Atu3255) | NA | 1 | 2 |
| 22740 (Atu4842) | NA | 1 | 2 |
| **Isolated in Δ*uppW* screen (annotated)** | | | |
| 06315 (Atu1281) | *nuoL* | 0 | 3 |
| 06730 (Atu1364) | *clpA* | 0 | 2 |
| 21635 (Atu4615) | *rfbA* | 0 | 2 |
| 21640 (Atu4616) | *rfbD* | 0 | 3 |

| | | | |
|---|---|---|---|
| **Isolated in Δ*uppW* screen (unannotated)** | | | |
| 03020 (Atu0613) | NA | 0 | 4 |
| 09720 (Atu1990) | NA | 0 | 3 |
| 13900 (Atu3023) | NA | 0 | 3 |
| 16905 (Atu3640) | NA | 0 | 6 |
| 21600 (Atu4606) | NA | 0 | 3 |
| 22700 (Atu4833) | NA | 0 | 2 |
| 24105 (Atu5033) | NA | 0 | 3 |
| **Isolated in Δ*uppY* screen (annotated)** | | | |
| 11550 (Atu2370) | *uppG* | 11 | 0 |
| 11555 (Atu2371) | *uppH* | 3 | 0 |
| 11500 (Atu2358) | *uppZ* | 3 | 0 |
| 02460 (Atu0499) | *podJ* | 9 | 0 |
| **Isolated in Δ*uppY* screen (unannotated)** | | | |
| 05430 (Atu1097) | NA | 4 | 0 |

### Table S4. Strains and plasmids used in this study

| Strain or plasmid | Relevant features | Source |
|---|---|---|
| ***E. coli*** | | |
| S17-1/λ*pir* | λ-pir, Tra+ donor for plasmid conjugation, $Sm^R$ | (Simon *et al.*, 1983) |
| SM10/ λ*pir* | λ-pir, Tra+ donor for Mariner *Himar 1* transposon conjugative delivery, $Km^R$ | (Miller & Mekalanos, 1988) |
| DH5α/λ*pir* | λ-pir cloning strain | (Chiang & Rubin, 2002) |
| TOP 10 F' | Cloning strain | Invitrogen |
| BL21 (DE3) | T7 expression strain | Stratagene |
| ***A. tumefaciens*** | | |
| C58 | Wild-type, nopaline-type, genomospecies G8 (*fabrum*) | (Lassalle *et al.*, 2011) |
| C58-JX110 | Δ*crdS* Δ*celH-D* Δ*exoA* Δ*chvAB,* (CDGS-) | (Xu *et al.*, 2012) |
| C58-JX137 | Δ*pruA* (ATU_RS05580, Atu1130) | (Xu *et al.*, 2013) |
| C58-JX138 | Δ*dcpA* (ATU_RS16190, Atu3495) | (Xu *et al.*, 2013) |
| C58-JX136 | Δ*claR* (ATU_RS08010, Atu1631) | (Xu *et al.*, 2013) |
| C58-JX117 | Δ*visR* (ATU_RS02585, Atu0525) | (Xu *et al.*, 2013) |
| C58-PMM26 | Δ*uppABCDEF* (ATU_RS06110-06125, Atu1235-40) | This study |
| C58-PMM27 | Δ*uppA* (ATU_RS06125, Atu1240) | This study |
| C58-PMM25 | Δ*uppB* (ATU_RS06120, Atu1239) | This study |
| C58-PMM22 | Δ*uppC* (ATU_RS06115, Atu1238) | This study |
| C58-PMM24 | Δ*uppD* (ATU_RS06110, Atu1237) | This study |
| C58-PMM13 | Δ*uppE* (ATU_RS06105, Atu1236) | (Xu *et al.*, 2012) |
| C58-PMM19 | Δ*uppF* (ATU_RS06100, Atu1235) | This study |
| C58-PMM32 | Δ*exoA* Δ*exoQ* (ATU_RS15360, Atu3325) | This study |
| C58-JX190 | Δ*uppG* (ATU_RS11550, Atu2370) | This study |
| C58-RN10 | Δ*uppH* (ATU_RS11555, Atu2371) | This study |
| C58-MCO051 | Δ*uppI* (ATU_RS11560, Atu2372) | This study |
| C58-JX122 | Δ*uppJ (*ATU_RS11565, Atu2373*)* | This study |
| C58-MCO048 | Δ*uppK* (ATU_RS11570, Atu2374) | This study |
| C58-MCO003 | Δ*uppL* (ATU_RS11575, Atu2375) | This study |
| C58-MCO052 | Δ*uppM* (ATU_RS11585, Atu2377) | This study |
| C58-MCO050 | Δ*uppN* (No annotation, Atu2378) | This study |
| C58-MCO049 | Δ*uppO* (ATU_RS11590, Atu2379) | This study |
| C58-JX179 | Δ*uppP* (ATU_RS17565, Atu3777) | This study |
| C58-JX177 | Δ*uppQ* (ATU_RS16325, Atu8188) | This study |
| C58-RN11 | Δ*uppV* (ATU_RS16320, Atu3523) | This study |
| C58-PMM29 | Δ*uppW* (ATU_RS11495, Atu2356) | This study |
| C58-JX121 | Δ*uppX* (ATU_RS02360, Atu0480) | This study |
| C58-JX180 | Δ*uppY* (ATU_RS02370, Atu0481) | This study |
| C58-MCO067 | Δ*uppX* Δ*uppV* | This study |
| C58-JX185 | Δ*uppY* Δ*uppW* | This study |
| C58-PMM32 | Δ*exoQ* (ATU_RS15360, Atu3325), Δ*exoA* (ATU_RSRS18935, Atu4053) | This study |
| C58-JX184 | Δ*exoQ* Δ*uppY* Δ*exoA* | This study |
| C58-JX186 | C58-JX110 (CDGS-), Δ*pruA* Δ*uppG* | This study |
| C58-MCO070 | C58-JX151 (CDGS-), Δ*pruA*, Δ*uppY* | This study |
| C58-MCO071 | C58-JX151 (CDGS-), Δ*pruA*, Δ*uppW* | This study |
| C58-MCO072 | C58-JX151 (CDGS-), Δ*pruA*, Δ*uppY*, Δ*uppW* | This study |
| C58-MCO073 | Δ*uppK*, Δ*pruA* | This study |
| C58-MCO075 | Δ*uppM*, Δ*pruB* | This study |

| Plasmids | | |
|---|---|---|
| pGEM-T Easy | PCR cloning vector, $Ap^R$ | Promega |
| pCR2.1 TOPO | PCR cloning vector, $Ap^R$ | Invitrogen |
| pNPTS138 | ColE1 suicide plasmid; *sacB*; $Km^R$ | (Morton & Fuqua, 2012) |
| pET-15b | $His_6$-expression vector, $Ap^R$ | EMD Biosciences |
| pFD1 | Mariner *Himar1* suicide delivery plasmid, $Km^R$, $Ap^R$ | (Lampe *et al.*, 1999) |
| pBBR-MCS5 | Broad host range expression $P_{lac}$ expression vector, $Gm^R$ | (Kovach *et al.*, 1995) |
| pSRKGm | Broad host range expression $P_{lac}$ expression vector plasmid, $lacI^Q$, $Gm^R$ | (Khan *et al.*, 2008) |
| pSRKKm | Broad host range expression $P_{lac}$ expression vector plasmid, $lacI^Q$, $Km^R$ | (Khan *et al.*, 2008) |
| pPM179 | pSRKKm carrying $P_{lac}$-*uppA*, $Km^R$ | This study |
| pPM178 | pSRKKm carrying $P_{lac}$-*uppB*, $Km^R$ | This study |
| pRN005 | pSRKGm carrying $P_{lac}$-*uppC*, $Gm^R$ | This study |
| pRN007 | pSRKGm carrying $P_{lac}$-*uppCD*, $Gm^R$ | This study |
| pPM177 | pSRKKm carrying $P_{lac}$-*uppD*, $Km^R$ | This study |
| pPM128 | pBBR-MCS5 carrying $P_{lac}$-*uppE*, $Gm^R$ | This study |
| pGH679 | pBBR-MCS5 carrying $P_{lac}$-*uppF*, $Gm^R$ | This study |
| pJX176 | pSRKGm carrying $P_{lac}$-*uppG*, $Gm^R$ | This study |
| pJX523 | pSRKGm carrying $P_{lac}$-$uppG^{D142A}$, $Gm^R$ | This study |
| pRN041 | pSRKGm carrying $P_{lac}$-*uppH*, $Gm^R$ | This study |
| pRN39 | pET-15b carrying T7-$His_6$-*uppH* | This study |
| pRN55 | pET-15b carrying T7-$His_6$-*uppH* E152Q E266Q | This study |
| pMCO017 | pSRKGm carrying $P_{lac}$-*uppI*, $Gm^R$ | This study |
| pJX115 | pSRKGm carrying $P_{lac}$-*uppJ*, $Gm^R$ | This study |
| pMCO018 | pSRKGm carrying $P_{lac}$-*uppM*, $Gm^R$ | This study |
| pMCO016 | pSRKGm carrying $P_{lac}$-*uppN*, $Gm^R$ | This study |
| pRN003 | pSRKGm carrying $P_{lac}$-*uppP*, $Gm^R$ | This study |
| pRN004 | pSRKGm carrying $P_{lac}$-*uppQ*, $Gm^R$ | This study |
| pRN042 | pSRKGm carrying $P_{lac}$-*uppV*, $Gm^R$ | This study |
| pMCO006 | pSRKGm carrying $P_{lac}$-*uppW,* $Gm^R$ | This study |
| pRN50 | pSRKGm carrying $P_{lac}$-*uppX*, $Gm^R$ | This study |
| pMCO005 | pSRKGm carrying $P_{lac}$-*uppY,* $Gm^R$ | This study |
| pJX612 | pSRKGm carrying $P_{lac}$-*pleD,* $Gm^R$ | This study |

### Supplementary References

Chiang, S.L., and Rubin, E.J. (2002) Construction of a mariner-based transposon for epitope-tagging and genomic targeting. *Gene* **296**: 179-185.

Khan, S.R., Gaines, J., Roop, R.M., 2nd, and Farrand, S.K. (2008) Broad-host-range expression vectors with tightly regulated promoters and their use to examine the influence of TraR and TraM expression on Ti plasmid quorum sensing. *Appl Environ Microbiol* **74**: 5053-5062.

Kovach, M.E., Elzer, P.H., Hill, D.S., Robertson, G.T., Farris, M.A., Roop, R.M.I., and Peterson, K.M. (1995) Four new derivatives of the broad-host-range cloning vector pBBR1MCS, carrying different antibiotic resistance cassettes. *Gene* **166**: 175-176.

Lampe, D.J., Akerley, B.J., Rubin, E.J., Mekalanos, J.J., and Robertson, H.M. (1999) Hyperactive transposase mutants of the *Himar1* mariner transposon. *Proc. Natl. Acad. Sci. USA* **96**: 11428-11433.

Lassalle, F., Campillo, T., Vial, L., Baude, J., Costechareyre, D., Chapulliot, D., Shams, M., Abrouk, D., Lavire, C., Oger-Desfeux, C., Hommais, F., Gueguen, L., Daubin, V., Muller, D., and Nesme, X. (2011) Genomic species are ecological species as revealed by comparative genomics in *Agrobacterium tumefaciens*. *Genome Biol Evol* **3**: 762-781.

Miller, V.L., and Mekalanos, J.J. (1988) A novel suicide vector and its use in construction of insertion mutations:osmoregulation of outer membrane proteins and virulence determinants in *Vibrio cholerae*. *J. Bacteriol.* **170**: 2575-2583.

Morton, E.R., and Fuqua, C. (2012) Genetic manipulation of *Agrobacterium*. *Curr Protoc Microbiol*: Unit 3D 2.

Simon, R., Priefer, U., and Puhler, A. (1983) A broad host range mobilization system for *in vivo* genetic engineering: transposon mutagenesis in gram negative bacteria. *Bio/Technology (Nature Publishing Company)* **Nov.**: 784-791.

Whitfield, C. (2006) Biosynthesis and assembly of capsular polysaccharides in *Escherichia coli*. *Annu. Rev. Biochem.* **75**: 39-68.

Xu, J., Kim, J., Danhorn, T., Merritt, P.M., and Fuqua, C. (2012) Phosphorus limitation increases attachment in *Agrobacterium tumefaciens* and reveals a conditional functional redundancy in adhesin biosynthesis. *Res Microbiol* **163**: 674-684.

Xu, J., Kim, J., Koestler, B.J., Choi, J.H., Waters, C.M., and Fuqua, C. (2013) Genetic analysis of *Agrobacterium tumefaciens* unipolar polysaccharide production reveals complex integrated control of the motile-to-sessile switch. *Mol Microbiol* **89**: 929-948.

**Supplementary Figure Legends**

**Figure S1. Biofilm formation and lectin labeling of genes identified in the DCR genetic screen.** (A) $A_{600}$ of acetic acid-solubilized CV absorbance from 48 h biofilm assays as a ratio to $OD_{600}$ from the planktonic phase of the same culture. Ratio for wild type set to 100% and mutant derivatives expressed as percent wild type. Bars represent the mean of triplicate assays and error bars are standard deviation. Bars with the same letter have means that are not significantly different from: a, WT C58; b, Δ*uppX* c, Δ*uppV*; d, *uppO* (Ordinary one-way ANOVA, *P* value < 0.05). Color coding of bars matches gene map (Fig. 2) and model (Fig. 12). (B) af-WGA labeling of *A. tumefaciens* C58 WT and selected mutant derivatives on 1.5% agar pads using a Nikon E800 epifluorescence microscope under 100x oil immersion using NIS-Elements software. Arrows indicate labeled UPP for WT.

**Figure S2. Complementation and related phenotypes of selected genes identified in the DCR genetic screen.** (A) $A_{600}$ of acetic acid-solubilized CV absorbance from 48 h biofilm assays Δ*uppL,* Δ*uppH and* Δ*uppN* and their plasmid-harboring derivatives as a ratio to $OD_{600}$ from the planktonic phase of the same culture in the presence of 500 μM IPTG. Gene names in parentheses indicate $P_{lac}$ expression constructs. (B) Biofilm assays with Δ*uppK* mutant in C58 WT and in Δ*pruA* mutant. (C) Biofilm assays for Δ*uppB*, Δ*uppM*, and the Δ*uppB* Δ*uppM* mutants performed as above. Ratio for wild type set to 100% and mutant derivatives expressed as percent wild type. Bars represent the mean of triplicate values, error bars are standard deviation. * comparison with wild type, and **comparison between Δ*uppM* and Δ*uppB* Δ*uppM;* standard *t*-test ($P$ = 0.05 or lower).

**Figure S3. WGA Lectin-binding and Congo red phenotypes of *uppG* deletion mutant.** (A) af-WGA labeling of *A. tumefaciens* WT and ΔATU_RS11550 (Δ*uppG)* on 1.5% agar pads using a Nikon E800 epifluorescence microscope under 100x oil immersion using NIS-Elements software. (B) Quantification of UPP fluorescence intensities in WT and Δ*uppG*. 100 cells counted per for each strain. (C) Congo red phenotypes of Δ*uppG* and its derivatives ectopically expressing the gene and the catalytic site *upp*G$^{AXD}$ mutant from $P_{lac}$ in a CDGS- Δ*pruA* background; 48 h incubation with 500 μM IPTG.

**Figure S4. *exoC* mutant phenotypes in *A. tumefaciens.*** Comparison of C58 WT, an *exoC* mutant with a *Himar1* insertion (*exoC::Mn*) and other mutant derivatives. (A) Congo red phenotypes of the indicated strains harboring a $P_{lac}$-*pleD* plasmid capable of driving production of cdGMP in the presence of 500 μM IPTG. (B) af-WGA lectin binding assay of WT and *exoC*::Mn on 1.5 % agar pads with 500 μM IPTG using a Nikon E800 epifluorescence microscope under 100x oil immersion using NIS-Elements software. (C) Calcofluor white assay (200 μg/mL) for succinoglycan production assay on LB. (D) Swim assay on 0.3% ATGN motility agar after 24 h incubation.

**Fig. S5. UppH inhibits biofilm formation.** $A_{600}$ of acetic acid-solubilized CV absorbance from 48 h biofilm inoculated with the addition of the indicated range of purified $His_6$-UppH (0.025-5.0 μM) and incubated at 30ºC. No growth inhibition was observed with addition of the enzyme so raw $A_{600}$ values were plotted. Bars represent the mean of triplicate measurements, error bars are standard deviation. All treatments were significantly decreased relative to wild type (Ordinary one-way ANOVA, $P < 0.05$) with a consistent downward trend with increasing enzyme concentration.

**Figure S6. Strategy for UPP pathway-specific screen.** In contrast to individual mutant characterization and isolation of the interrupted sequences, transposon mutants with the DCR phenotype were pooled and included in *en masse*, bar-coded sequencing of transposon right borders. Sequences from each parent background were bar-coded to enable multiplex sequencing.

**Fig. S7. Multi-locus sequence analysis (MLSA) phylogeny of agrobacteria/rhizobia genomes for genes identified in DCR screens.** The presence or absence of key polysaccharide genes in each strain are indicated by a heat map. Genes present in a strain (based on reciprocal best blast hits to C58) are blue, genes that are absent are white. Light blue indicates genes that are present but with low (<50%) protein sequence identity to C58. Support for tree branches is indicated by color. Branches with Ultrafast bootstrap > 95% and SH-aLRT test > 80% are colored black, unsupported branches are gray. Clades containing agrobacteria biovars and BV1 genomospecies, as well as other key lineages, are labeled.

**Figure S8. Comparison of synteny and gene content of various *upp* loci across lineages.** The presence and organization of key *upp* genes identified in the DCR screens are shown as arrows. Arrow direction indicates strand. Genes are colored by locus, genes colored gray are not present in reference strain C58 and are variably present. Each locus is found in a different location across the C58 reference genome and loci are not shown in order. Lines connecting genes of different strains indicate homology, darker colors indicate higher percent identity.

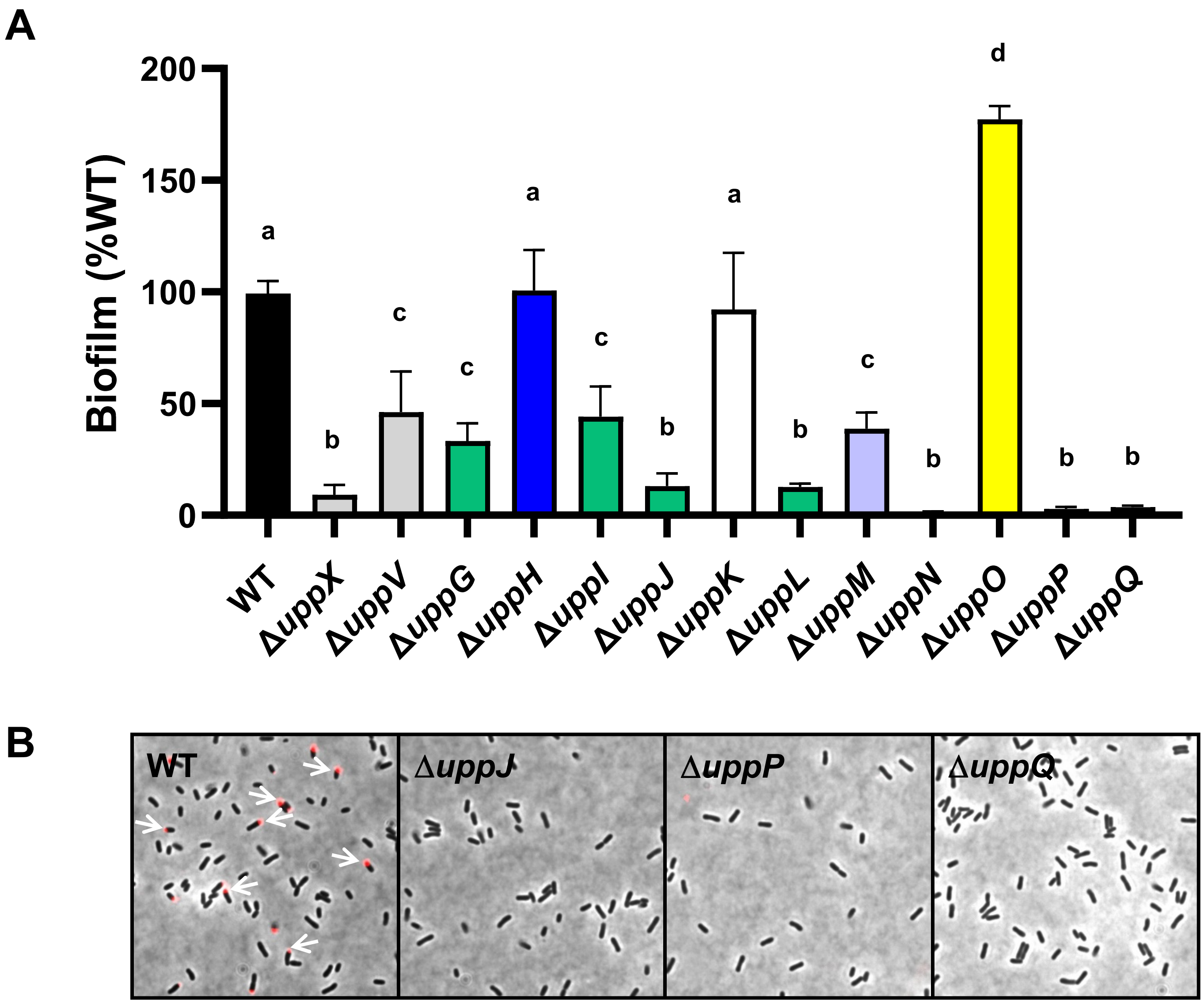

Figure S1 - Onyeziri et al.

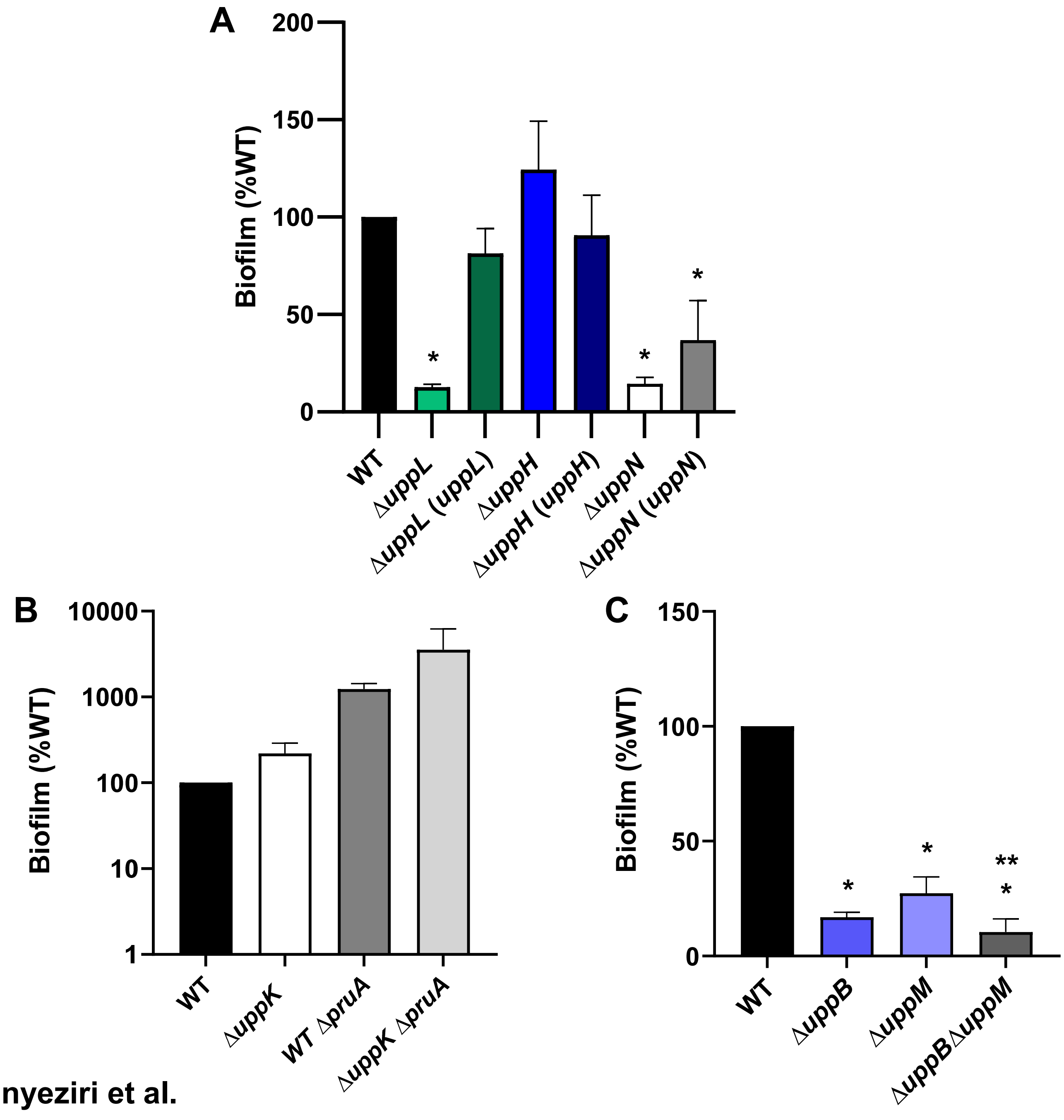

Figure S2 - Onyeziri et al.

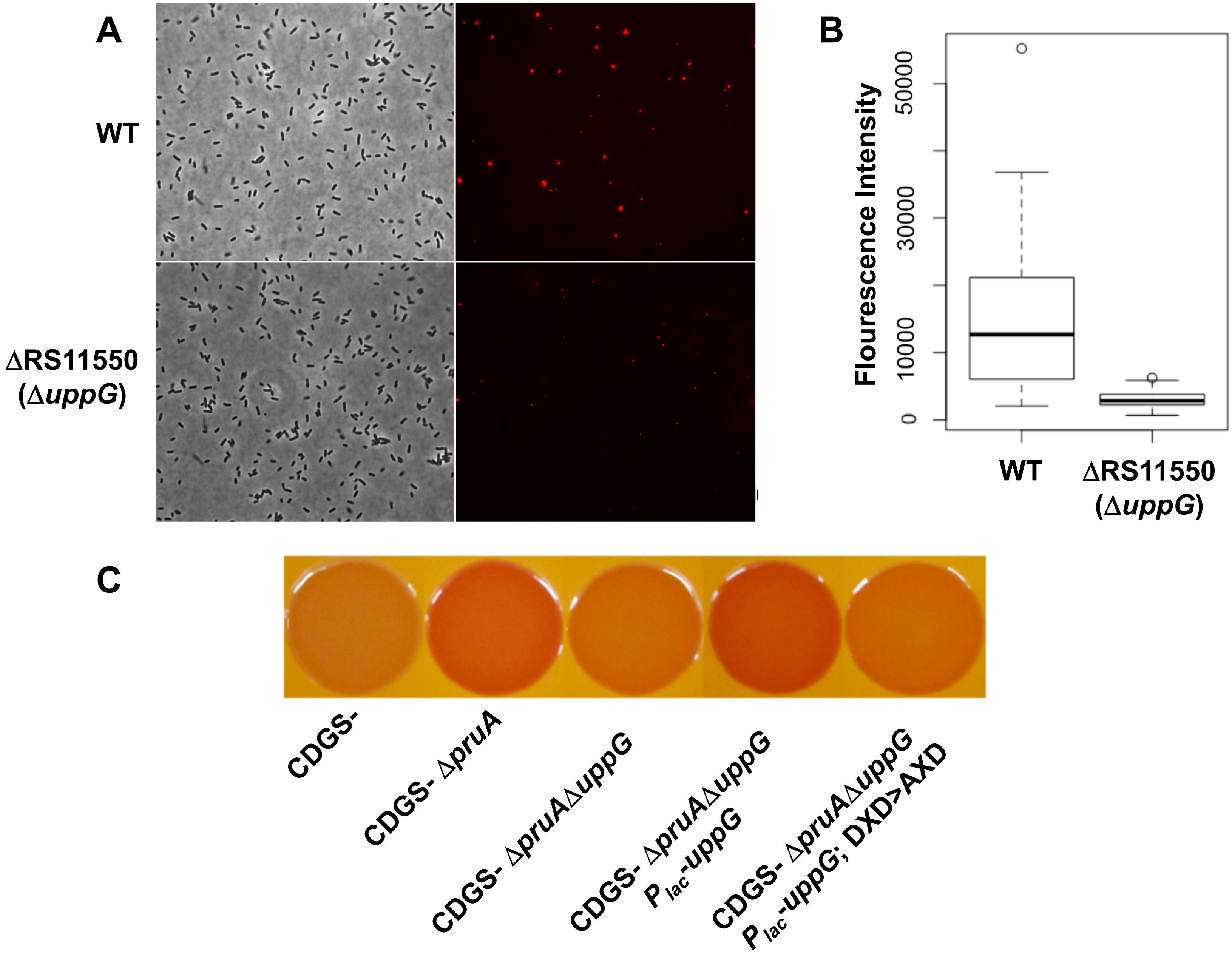

Figure S3 - Onyeziri et al.

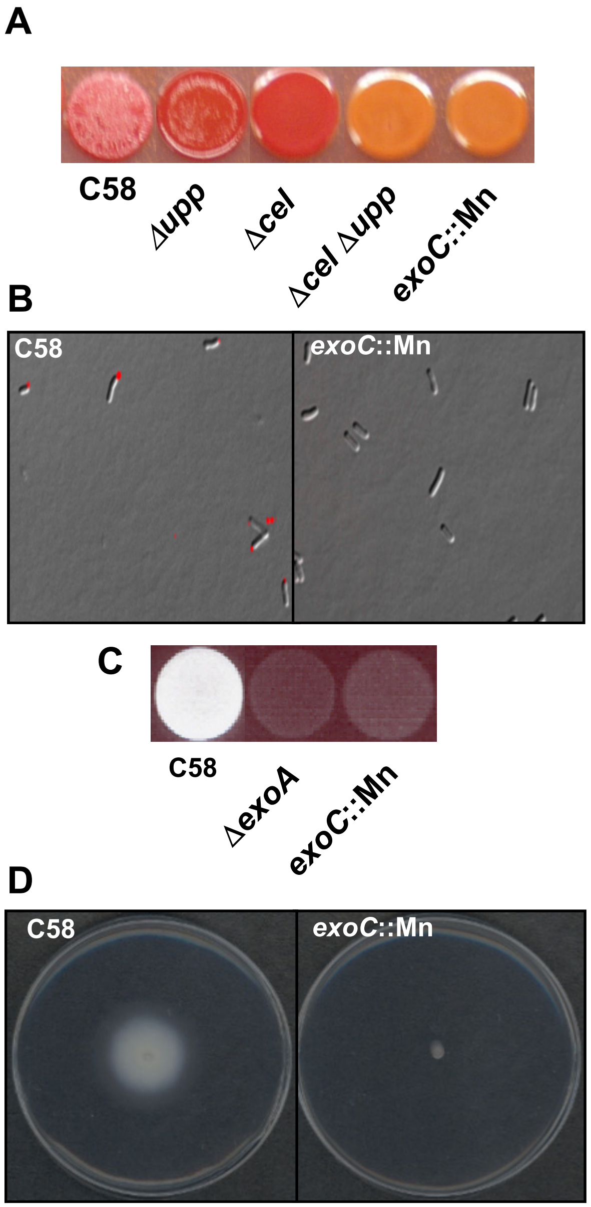

**Figure S4 - Onyeziri et al.**

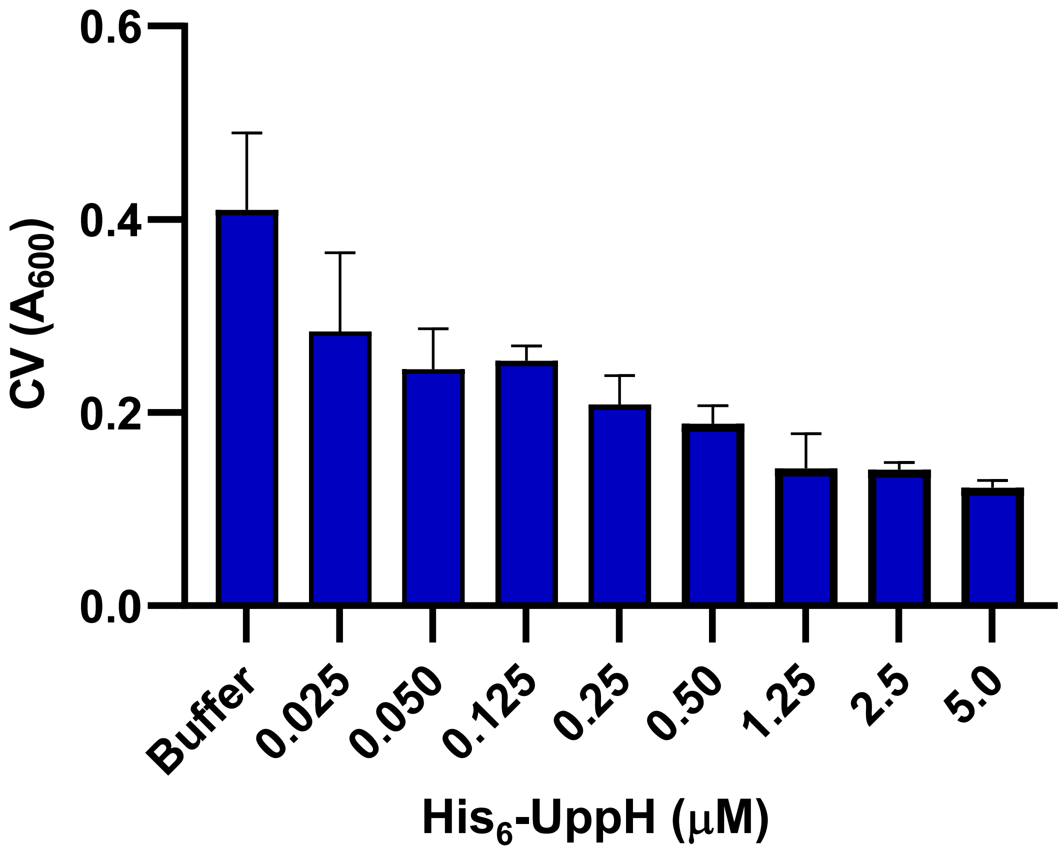

**Figure S5 - Onyeziri et al.**

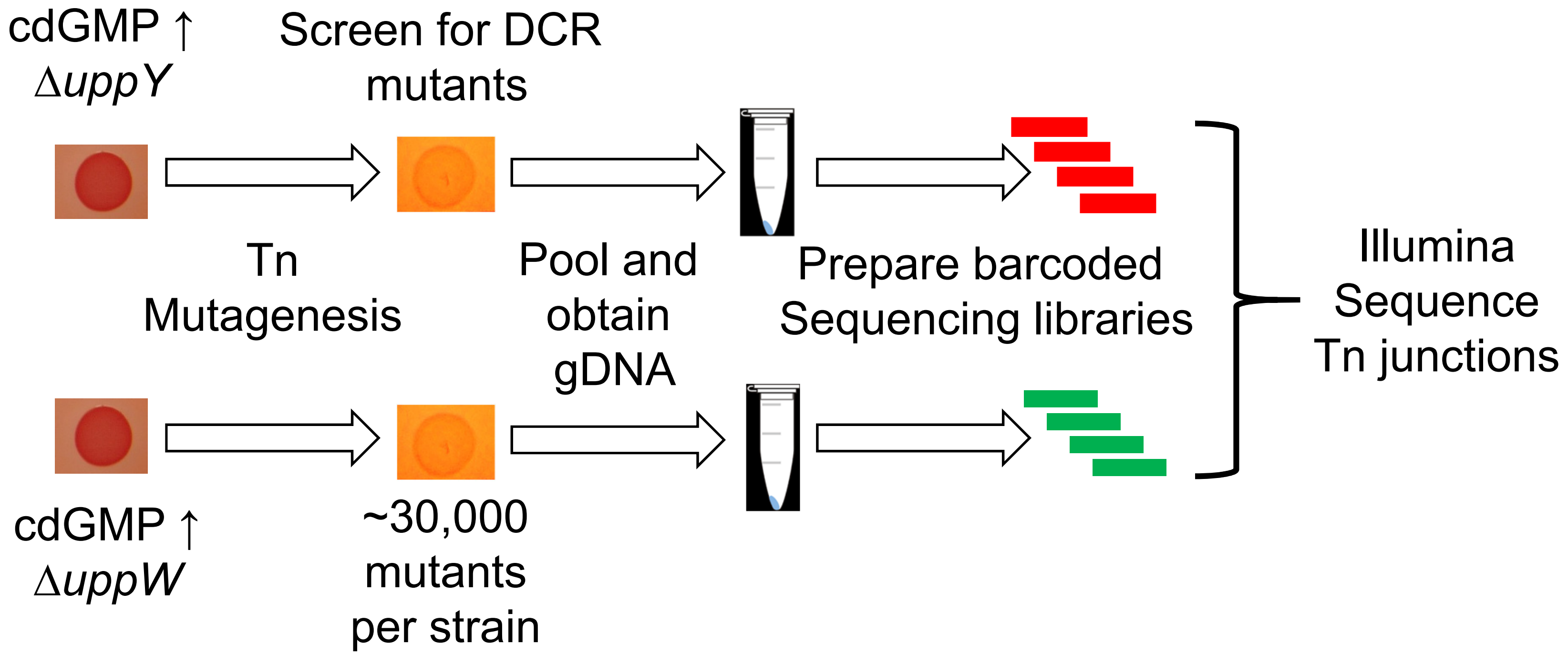

**Figure S6 - Onyeziri et al.**

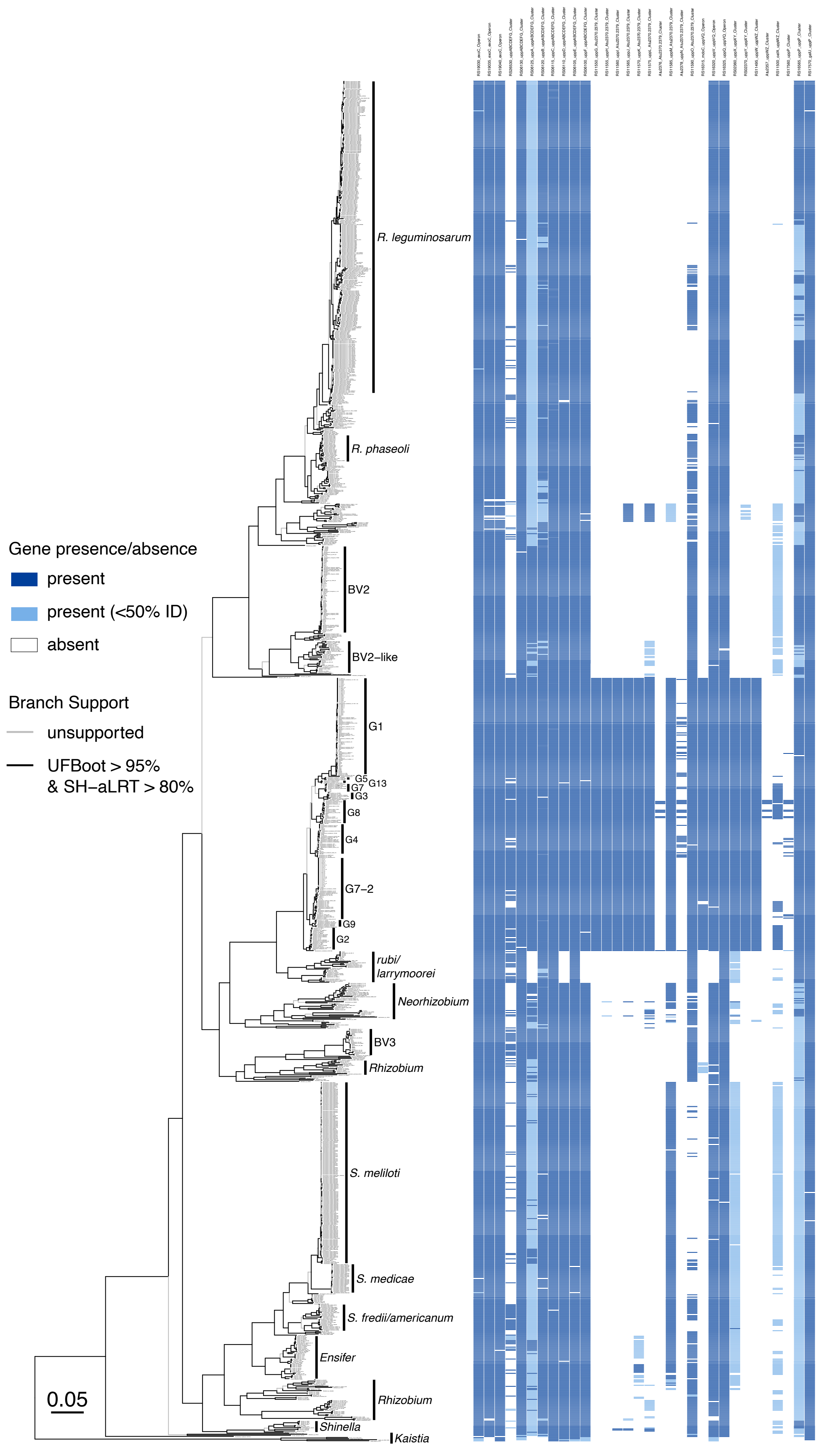

RS19030_exoC_Operon
RS19035_exoC_exoC_Operon
RS19040_exoC_Operon
RS26530_uppABCDEFG_Cluster
RS06130_uppABCDEFG_Cluster
RS06125_uppA_uppABCDEFG_Cluster
RS06120_uppB_uppABCDEFG_Cluster
RS06115_uppC_uppABCDEFG_Cluster
RS06110_uppD_uppABCDEFG_Cluster
RS06105_uppE_uppABCDEFG_Cluster
RS06100_uppF_uppABCDEFG_Cluster
RS11550_uppG_Atu2370.2379_Cluster
RS11555_uppH_Atu2370.2379_Cluster
RS11560_uppI_Atu2370.2379_Cluster
RS11565_uppJ_Atu2370.2379_Cluster
RS11570_uppK_Atu2370.2379_Cluster
RS11575_uppL_Atu2370.2379_Cluster
Atu2376_Atu2370.2379_Cluster
RS11585_uppM_Atu2370.2379_Cluster
Atu2378_uppN_Atu2370.2379_Cluster
RS11590_uppO_Atu2370.2379_Cluster
RS16315_mdoC_uppVQ_Operon
RS16320_uppV_uppVQ_Operon
RS16325_uppQ_uppVQ_Operon
RS02360_uppX_uppXY_Cluster
RS02370_uppY_uppXY_Cluster
RS11495_uppW_uppWZ_Cluster
Atu2357_uppWZ_Cluster
RS11500_oafA_uppWZ_Cluster
RS17560_uppP_Cluster
RS16565_uppP_uppP_Cluster
RS17570_galU_uppP_Cluster
R. leguminosarum
R. phaseoli
BV2
BV2–like
G1
G5
G13
G7
G3
G8
G4
G7–2
G9
G2
rubi/
larrymoorei
Neorhizobium
BV3
Rhizobium
S. meliloti
S. medicae
S. fredii/americanum
Ensifer
Rhizobium
Shinella
Kaistia
Gene presence/absence
present
present (<50% ID)
absent
Branch Support
unsupported
UFBoot > 95%
& SH–aLRT > 80%
0.05

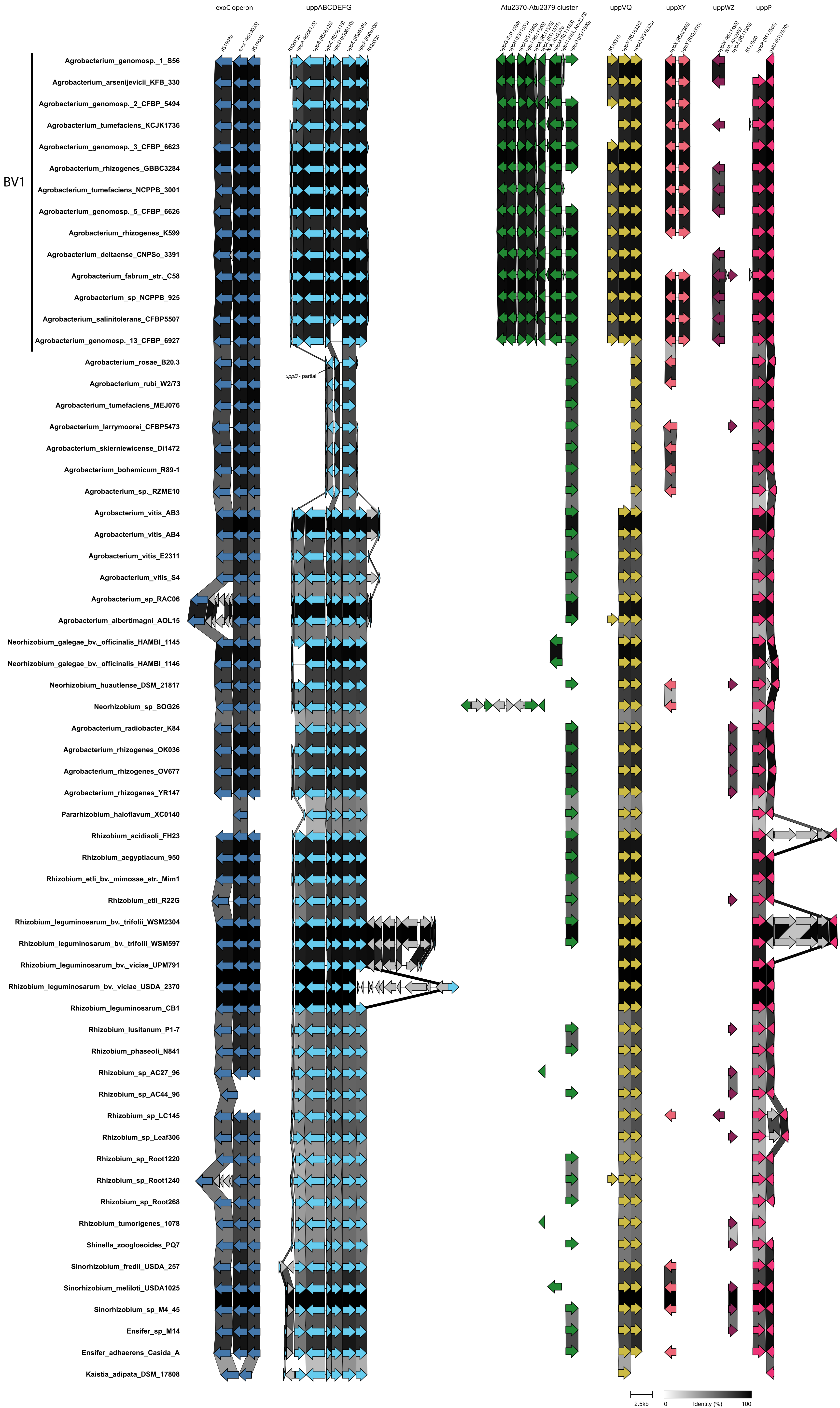

exoC operon
uppABCDEFG
Atu2370-Atu2379 cluster
uppVQ
uppXY
uppWZ
uppP
RS19030
exoC (RS19035)
RS19040
RS06130
uppA (RS06125)
uppB (RS06120)
uppC (RS06115)
uppD (RS06110)
uppE (RS06105)
uppF (RS06100)
RS26530
uppG (RS11550)
uppH (RS11555)
uppI (RS11560)
uppJ (RS11565)
uppK (RS11570)
uppL (RS11575)
N/A, Atu2376
uppM (RS11585)
uppN (N/A, Atu2378)
uppO (RS11590)
RS16315
uppV (RS16320)
uppQ (RS16325)
uppX (RS02360)
uppY (RS02370)
uppW (RS11495)
N/A, Atu2357
uppZ (RS11500)
RS17560
uppP (RS17565)
galU (RS17570)
BV1
Agrobacterium_genomosp._1_S56
Agrobacterium_arsenijevicii_KFB_330
Agrobacterium_genomosp._2_CFBP_5494
Agrobacterium_tumefaciens_KCJK1736
Agrobacterium_genomosp._3_CFBP_6623
Agrobacterium_rhizogenes_GBBC3284
Agrobacterium_tumefaciens_NCPPB_3001
Agrobacterium_genomosp._5_CFBP_6626
Agrobacterium_rhizogenes_K599
Agrobacterium_deltaense_CNPSo_3391
Agrobacterium_fabrum_str._C58
Agrobacterium_sp_NCPPB_925
Agrobacterium_salinitolerans_CFBP5507
Agrobacterium_genomosp._13_CFBP_6927
Agrobacterium_rosae_B20.3
uppB - partial
Agrobacterium_rubi_W2/73
Agrobacterium_tumefaciens_MEJ076
Agrobacterium_larrymoorei_CFBP5473
Agrobacterium_skierniewicense_Di1472
Agrobacterium_bohemicum_R89-1
Agrobacterium_sp._RZME10
Agrobacterium_vitis_AB3
Agrobacterium_vitis_AB4
Agrobacterium_vitis_E2311
Agrobacterium_vitis_S4
Agrobacterium_sp_RAC06
Agrobacterium_albertimagni_AOL15
Neorhizobium_galegae_bv._officinalis_HAMBI_1145
Neorhizobium_galegae_bv._officinalis_HAMBI_1146
Neorhizobium_huautlense_DSM_21817
Neorhizobium_sp_SOG26
Agrobacterium_radiobacter_K84
Agrobacterium_rhizogenes_OK036
Agrobacterium_rhizogenes_OV677
Agrobacterium_rhizogenes_YR147
Pararhizobium_haloflavum_XC0140
Rhizobium_acidisoli_FH23
Rhizobium_aegyptiacum_950
Rhizobium_etli_bv._mimosae_str._Mim1
Rhizobium_etli_R22G
Rhizobium_leguminosarum_bv._trifolii_WSM2304
Rhizobium_leguminosarum_bv._trifolii_WSM597
Rhizobium_leguminosarum_bv._viciae_UPM791
Rhizobium_leguminosarum_bv._viciae_USDA_2370
Rhizobium_leguminosarum_CB1
Rhizobium_lusitanum_P1-7
Rhizobium_phaseoli_N841
Rhizobium_sp_AC27_96
Rhizobium_sp_AC44_96
Rhizobium_sp_LC145
Rhizobium_sp_Leaf306
Rhizobium_sp_Root1220
Rhizobium_sp_Root1240
Rhizobium_sp_Root268
Rhizobium_tumorigenes_1078
Shinella_zoogloeoides_PQ7
Sinorhizobium_fredii_USDA_257
Sinorhizobium_meliloti_USDA1025
Sinorhizobium_sp_M4_45
Ensifer_sp_M14
Ensifer_adhaerens_Casida_A
Kaistia_adipata_DSM_17808
2.5kb
0
Identity (%)
100
